# Patient-specific functional brain architecture explains cortical patterns of tau PET in Alzheimer’s disease

**DOI:** 10.1101/2025.10.02.679969

**Authors:** Harry H. Behjat, Jacob W. Vogel, Olof Strandberg, Nicola Spotorno, Nicolai Franzmeier, Léa Chauveau, Jonathan Rittmo, Sara Stampacchia, Lyduine E. Collij, Alexa Pichet Binette, Yu Xiao, Pavanjit Chaggar, Zeyu Zhu, the Alzheimer’s Disease Neuroimaging Initiative, Danielle van Westen, Erik Stomrud, Sebastian Palmqvist, Dimitri Van De Ville, Ruben Smith, Niklas Mattsson-Carlgren, Oskar Hansson, Rik Ossenkoppele

**Affiliations:** Clinical Memory Research Unit, Department of Clinical Sciences Malmö, Faculty of Medicine, Lund University, Lund, Sweden; Neuro-X Institute, École Polytechnique Fédérale de Lausanne, Geneva, Switzerland; Neurodegenerative Research Unit, Department of Clinical Sciences Malmö, Faculty of Medicine, SciLifeLab, LU, Lund, Sweden; Radiology and Nuclear Medicine, Amsterdam UMC, location VUmc, Amsterdam, the Netherlands; Brain Imaging, Amsterdam Neuroscience, Amsterdam, the Netherlands; Department of Physiology and Pharmacology, Université de Montréal, Montréal, Quebec, Canada; Centre de Recherche de l’Institut Universitaire de Gériatrie de Montréal, Montréal, Quebec, Canada; Department of Diagnostic Radiology, Department of Clinical Sciences Lund, Lund University, Lund, Sweden; Memory Clinic, Skåne University Hospital, SE-205 02 Malmö, Sweden; Department of Radiology and Medical Informatics, University of Geneva, Geneva, Switzerland; Alzheimer Center Amsterdam, Neurology, Vrije Universiteit Amsterdam, Amsterdam UMC, Amsterdam, the Netherlands; Amsterdam Neuroscience, Neurodegeneration, Amsterdam, the Netherlands; Institute for Stroke and Dementia Research (ISD), University Hospital, LMU Munich, Munich, Germany; Department of Psychiatry and Neurochemistry, The Sahlgrenska Academy, Institute of Neuroscience and Physiology, University of Gothenburg, Gothenburg, Sweden; Munich Cluster for Systems Neurology (SyNergy), Munich, Germany

## Abstract

The spatial distribution of tau pathology, a key correlate of neurodegeneration and cognitive decline in Alzheimer’s disease (AD), varies markedly across individuals. While tau is thought to spread along brain networks, the role of interindividual variability in accounting for these patterns remains underexplored. Using resting-state fMRI and tau-PET from 805 BioFINDER participants across the AD continuum, with replication in ADNI (n=361) and A4 (n=336), we studied whether subject-specific functional connectivity (FC) profiles enhance the characterization of tau deposition patterns. A hybrid approach integrating individual and group-average FC explained individual tau-PET topographies better than either FC representation alone, particularly in symptomatic individuals and at finer spatial resolutions. Hybrid FC also better captured individual tau topographies than canonical tau-PET maps derived from cohort-level data. These effects were specific to tau and not similarly observed for β-amyloid, and the explanatory advantage of FC-based models increased with spatial granularity. Furthermore, baseline hybrid FC explained follow-up tau-PET topography better than template FC, suggesting that individualized baseline connectivity contains information about future tau-PET progression. The main FC model-comparison findings replicated in ADNI and A4. Collectively, these findings show that individual functional brain architecture is associated with heterogeneity in tau-PET topography. While not establishing a causal propagation mechanism, our findings are consistent with network-spread models. This work advances the methodological characterization of tau-PET heterogeneity in AD and highlights functional connectivity as a potentially informative marker of individual tau-PET trajectories.

## 1. Introduction

Alzheimer’s disease (AD), the leading cause of dementia, is characterized by the progressive accumulation of extracellular β-amyloid plaques and intracellular neurofibrillary tangles composed of hyperphosphorylated tau proteins (Scheltens et al., 2021). β-amyloid plaques can be detected in the brain decades prior to symptom onset (Hansson, 2021; Palmqvist et al., 2017), and are detected in 20-40% of cognitively unimpaired elderly (Villemagne et al., 2013; Hedden et al., 2013; Jansen et al., 2015; Jack et al., 2017; Jansen et al., 2022). In contrast, tau neurofibrillary tangles bear a strong temporal and spatial link to neurodegeneration and cognitive decline (Villemagne et al., 2018; Bejanin et al., 2017). The formation of tau tangles, generally follows a hierarchical spatial progression, beginning in the transentorhinal cortex and spreading to the hippocampus, association cortices, and eventually to primary sensory regions (Braak and Braak, 1991). This stereotypical pattern of tau progression has been observed both postmortem (Scheff et al., 2006; Braak et al., 2011; Marquié et al., 2017; Ravikumar et al., 2024) and in vivo using tau positron emission tomography (PET) (Schöll et al., 2016; Johnson et al., 2016; Sanchez et al., 2021; Therriault et al., 2022; Leuzy et al., 2023; Ossenkoppele et al., 2025). However, more recent studies show that tau burden patterns exhibit notable heterogeneity across individuals with AD, often diverging from the stereotypical Braak staging system (Ferreira et al., 2020; Vogel et al., 2021; Young et al., 2022). Understanding the biological mechanisms underlying variation in individual spatial patterns of tau progression is crucial, as these may have significant clinical implications. For instance, both postmortem and clinical imaging studies have demonstrated that neocortical-predominant (hippocampal-sparing) tau-PET patterns are linked to earlier symptom onset and faster cognitive decline, while limbic-predominant patterns are associated with later onset and slower decline (Murray et al., 2011; Ossenkoppele et al., 2020). Inter-individual variations in premorbid brain structure are thought to be a key contributor to variations in tau spread (Jacobs et al., 2018; Bocancea et al., 2023), a notion that is especially supported by an accumulating body of studies that report strong associations between the spatial structure of brain networks and the trajectory of tau spread in AD (Hoenig et al., 2018; Ossenkoppele et al., 2019; Franzmeier et al., 2020; Vogel et al., 2020).

Connectome-based disease spreading models have emerged as a powerful framework for simulating tau propagation through human brain networks, incorporating both functional and structural connectivity in predicting tau distribution (Raj et al., 2021; Vogel et al., 2023). These group-level models have demonstrated that tau may spread along neuronal connections, measured by diffusion tensor imaging, and clarify how various brain regions contribute to tau propagation (Fornari et al., 2019). These findings support an influential hypothesis that tau propagates in a “prion-like” manner, spreading from an epicenter, typically in the (trans)entorhinal cortex, along synaptically connected regions, facilitated by the brain’s network structure (Clavaguera et al., 2009; de Calignon et al., 2012; Goedert et al., 2017; Kaufman et al., 2018). Human studies and animal models show greater tau accumulation in regions strongly connected to initially affected areas, indicating that tau spread follows functional and anatomical connectivity patterns (Liu et al., 2012; Frost and Diamond, 2010; Yu et al., 2021).

Resting-state functional MRI (fMRI) is a valuable technique for mapping brain functional architecture, particularly in deriving functional connectivity networks that capture coherent brain activity between regions. Several studies have shown that tau-PET patterns can be predicted or simulated using group-averaged functional connectivity networks (Sepulcre et al., 2017; Cope et al., 2018; Franzmeier et al., 2019, 2020, 2022; Ossenkoppele et al., 2019; Vogel et al., 2020, 2023; Ottoy et al., 2024; Wang et al., 2024), providing evidence for a potential link between functional connectivity and tau pathology. While averaging functional connectivity patterns across individuals creates a template connectome that captures stable patterns, it fails to account for individual connectivity differences. Functional connectomes retain unique, person-specific features that allow reliable fingerprinting of individuals (Finn et al., 2015; Amico and Goñi, 2018) throughout the AD continuum, and have therefore been suggested as a means to inform personalized models of disease progression (Stampacchia et al., 2024). Importantly, if the prion-like spread hypothesis holds (Cope et al., 2018; Bejanin et al., 2017; Tucholka et al., 2018), these individual connectivity profiles may help explain inter-individual variation in tau-PET topography. In other words, the specific architecture of a person’s brain network may contribute to individual variation in not only the extent but also the pattern of tau burden, including which regions show greater involvement over time. Recognizing this individual-level variability is crucial, as it highlights connectivity not only as a population-level correlate but also as a candidate source of information for characterizing personalized disease trajectories. Consistent with this, recent studies have begun to advocate for patient-tailored connectivity models to capture the heterogeneity of tau progression across individuals (Vogel et al., 2023; Tripathi et al., 2025), with evidence that clinical subtypes of AD, such as early-versus late-onset, show distinct patterns of both tau accumulation and functional connectivity (Schöll et al., 2017; La Joie et al., 2020; Lee et al., 2022; Franzmeier et al., 2018). Recent work has begun to address this gap by incorporating individualized connectomic information into tau models. Abuwarda et al. (2025) showed that whole-brain resting-state functional connectivity can predict regional tau-PET signal in preclinical AD, particularly in later Braak-stage regions, highlighting the relevance of FC-derived features for tau-PET modelling. Complementarily, Ottoy et al. (2024) demonstrated that tau accumulation can be mapped within subject-specific functional and structural connectivity gradients, that is, low-dimensional representations of brain-network organization, with gradient-derived hubs explaining regionally distinct patterns of future tau accumulation. Building on these studies, the present work extends prior individualized-connectome approaches by explicitly comparing subject-specific, template, and hybrid FC representations; testing whether individualized FC provides added explanatory information beyond group-level template FC and canonical PET-pattern models; evaluating these effects across spatial scales; and assessing replication in two external cohorts. At the same time, this does not preclude alternative explanations such as the role of selective regional vulnerability in modulating tau accumulation patterns (Leng et al., 2021; Kampmann, 2024; Xiao et al., 2025).

This study aims to determine whether individual-level FC, derived from resting-state fMRI, enhances the statistical characterization of individual tau-PET patterns beyond what is achievable using group-level template FC. We hypothesize that individual FC patterns can account for more of the variance observed in tau-PET patterns across subjects compared to group-averaged connectivity. Importantly, we investigate this across multiple spatial resolutions (from 100 to 1000 cortical regions) using subject-space surface-based parcellations, enabling multiscale modelling of both FC and tau-PET. We further investigate whether FC more accurately reflects individual tau-PET distributions than canonical group-based on- and off-target binding patterns. Additionally, we test the specificity of this relationship by assessing whether the unique explanatory power of individual FC for tau is similarly observed in β-amyloid, for which the same network-spread hypothesis is not assumed. Finally, we test whether baseline FC helps explain follow-up tau-PET topography, thereby examining its longitudinal explanatory relevance. We additionally assess whether the main FC model-comparison findings replicate in two independent cohorts, ADNI and A4. Overall, our approach moves beyond the traditional reliance on group averages and addresses limitations of previous individualized FC studies by evaluating multiple FC representations across spatial scales and external cohorts, shedding light on how individual variations in functional connectivity relate to the spatial distribution of tau pathology along the AD continuum.

## 2. Results

Our main study included 805 elderly participants (mean [std]: 71 [9] years; 55% females) from the Swedish BioFINDER-2 study (https://biofinder.se/). Among them, 398 were cognitively normal (CN), 77 reported subjective cognitive decline (SCD), 156 were diagnosed with mild cognitive impairment (MCI), and 174 were diagnosed with AD dementia (ADD). All patients with ADD, MCI, and SCD, and a subset of the CN (n=89, 22.4%) were Aβ-positive based on CSF testing. Demographic information for each group is shown in Table 1. The data for each participant included structural and resting-state functional MRI and tau-PET. Aβ-PET was available for CN, SCD, and MCI participants but not for ADD patients. Furthermore, our longitudinal analysis included follow-up tau-PET scans from a subset of the participants (n=422, 52.4% of sample) with a mean follow-up time of 1.8*±*0.48 years.

**Table 1.** Demographics and clinical characteristics of the sample. Aβ: β-amyloid; Aβ status (positive: Aβ+, negative: Aβ−) was determined based on CSF Aβ42/Aβ40 ratio. Data are presented as mean *±*s.d. unless specified otherwise. CN: Cognitively normal; SCD: Subjective cognitive decline; MCI: mild cognitive impairment; ADD: Alzheimer’s disease dementia; MMSE: mini mental state examination; *APOE* ε4: apolipoprotein E genotype carrying at least one ε4 allele. ^a^Braak staging meta-ROI (Braak and Braak, 1991) was constructed using regions defined by (Cho et al., 2016), based on FreeSurfer labels. ^b^There are 4, 0, 0, 0, 174 missing values in the respective categories; value defined as the average SUVR of a global neocortical ROI (Ossenkoppele et al., 2022), including prefrontal, lateral temporal, parietal, anterior cingulate and posterior cingulate/praecuneus. ^c^Dementia patients did not undergo Aβ-PET.

| | CN Aβ-<br>(n=309) | CN Aβ+<br>(n=89) | SCD Aβ+<br>(n=77) | MCI Aβ+<br>(n=156) | ADD Aβ+<br>(n=174) | Overall<br>(n=805) |
| --- | --- | --- | --- | --- | --- | --- |
| Age (years) | 67.27 ± 10.95 | 76.29 ± 8.14 | 70.55 ± 6.96 | 72.74 ± 6.82 | 72.98 ± 7.70 | 70.88 ± 9.45 |
| Sex, female n (%F) | 188 (60.8) | 49 (55.1) | 37 (48.1) | 69 (44.2) | 99 (56.9) | 442 (54.9) |
| Education (years) | 12.67 ± 3.41 | 12.28 ± 3.92 | 13.09 ± 3.95 | 13.16 ± 4.48 | 12.08 ± 4.05 | 12.64 ± 3.89 |
| APOE status available n | 225 | 45 | 75 | 153 | 172 | 670 |
| APOE ε4 carriers n (% of available n) | 98 (43.0) | 34 (75.6) | 54 (72.0) | 117 (76.6) | 122 (70.9) | 425 (63.4) |
| MMSE | 29.05 ± 1.05 | 28.52 ± 1.32 | 28.55 ± 1.50 | 27.06 ± 1.89 | 21.11 ± 4.02 | 26.85 ± 3.81 |
| tau-PET SUVR (RO948), Braak Stages I-IV<sup>a</sup> | 1.14 ± 0.08 | 1.24 ± 0.18 | 1.33 ± 0.29 | 1.50 ± 0.43 | 2.12 ± 0.63 | 1.45 ± 0.52 |
| tau-PET SUVR (RO948), Braak Stages V-VI<sup>a</sup> | 1.04 ± 0.07 | 1.06 ± 0.11 | 1.09 ± 0.13 | 1.16 ± 0.20 | 1.52 ± 0.40 | 1.17 ± 0.29 |
| follow-up tau-PET, available n (female n) | 175 (101) | 46 (24) | 54 (24) | 73 (26) | 74 (44) | 422 (219) |
| tau-PET follow-up time (years) | 1.91 ± 0.53 | 1.88 ± 0.63 | 1.75 ± 0.33 | 1.74 ± 0.35 | 1.60 ± 0.41 | 1.80 ± 0.48 |
| Aβ-PET SUVR<sup>b</sup> (flutemetamol), meta ROI | 0.92 ± 0.04 | 1.34 ± 0.22 | 1.43 ± 0.23 | 1.52 ± 0.27 | Not available<sup>c</sup> | 1.20 ± 0.33 |

### 2.A. Patient-specific FC improves the explanation of individual tau-PET patterns beyond template FC

Throughout the following analyses, template FC refers to the group-derived FC matrix estimated from amyloid-negative cognitively healthy controls.

We first used regional FC–tau regression models to test whether subject-specific FC provided explanatory information beyond template FC and to obtain the region- and subject-specific coefficients later used to construct hybrid FC (Fig. 1a). For each subject, atlas resolution, and cortical seed region, tau-PET values across cortical parcels were modelled using template FC alone or template FC together with subject FC. Because these models differed in the number of regressors, all reported in-sample explained-variance values correspond to corrected *R*^2^, not raw *R*^2^. Thus, improvements in explained variance reflect additional model fit after penalizing for the inclusion of extra predictors. Across scales, the inclusion of subject FC consistently improved model fits relative to template FC alone, with the additional variance explained being most pronounced in later disease stages (Fig. 1b; Fig. S1a–b). The explained variance per region decreased with increasing atlas size, since regions became smaller; however, this decline was less steep than the decrease in region size itself, indicating a net gain in cumulative variance explained across regions (Fig. 1b, bottom; Fig. S1b). We next evaluated whether this regional FC– tau correspondence could be explained by lower-order network properties or by spatial autocorrelation in tau-PET. Degree- and strength-preserving rewired FC models produced substantially lower explained variance than empirical subject FC, indicating that the added explanatory value of subject FC depended on its specific wiring pattern rather than preserved degree or strength alone (Fig. 1c; Fig. S1c).

**Fig. 1.**
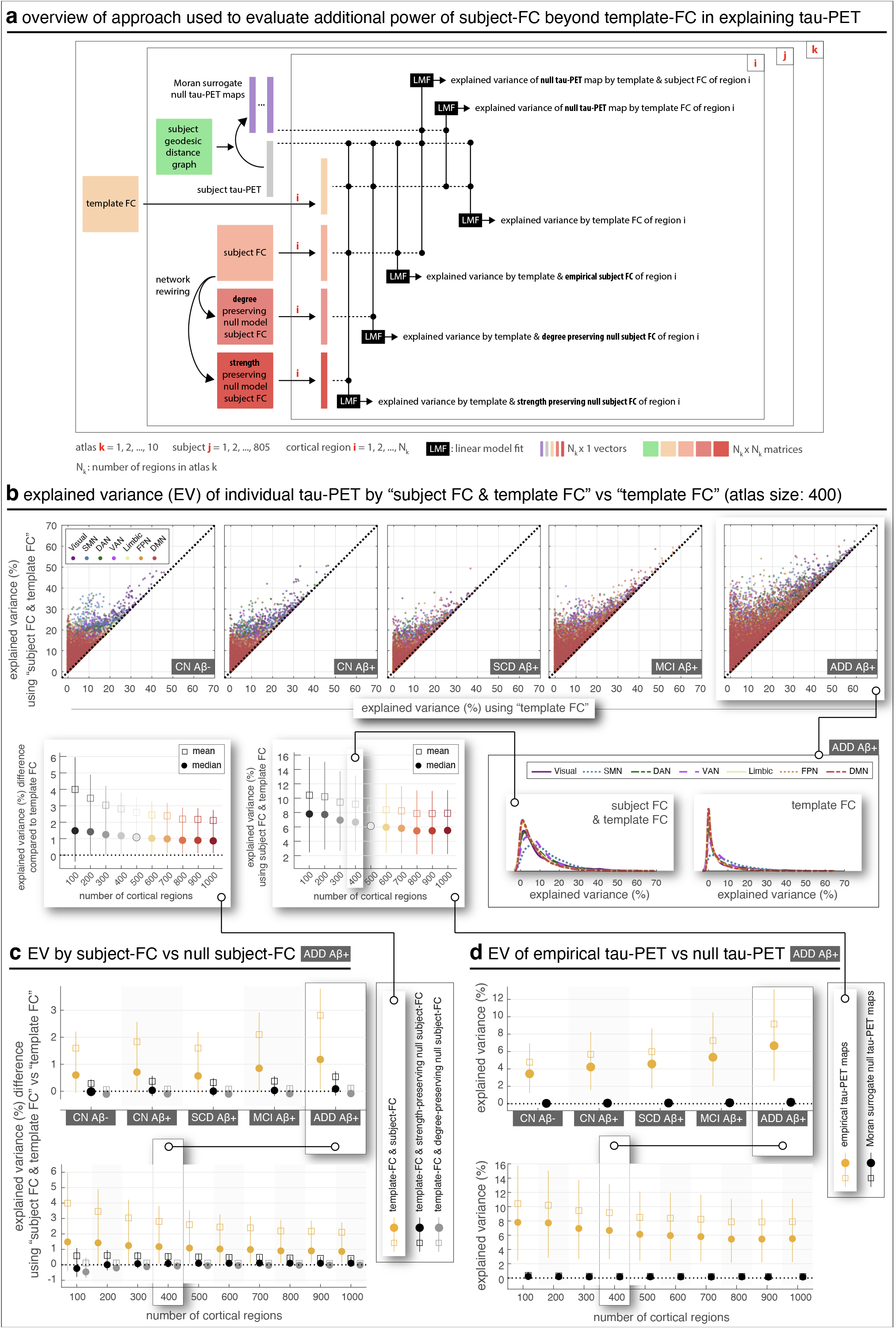
Regional subject FC provides additional explanatory power in explaining individual tau-PET beyond template FC, across spatial scales. (a) Overview of the approach used to evaluate the additional power of subject FC beyond template FC in explaining regional tau-PET patterns. By “explained variance”, we refer to the coefficient of determination (*R*^2^) from model fits; throughout this work, we report corrected *R*^2^ values to facilitate comparison between models with differing numbers of regressors. In this figure, the model using only template FC includes one regressor, whereas the model using both template FC and subject FC includes two. “Subject FC + template FC” denotes a joint regression model in which subject-specific FC and template FC were entered simultaneously as separate predictors. This differs from hybrid FC, which was constructed as a weighted combination of the two FC profiles. Rewired null models were used to assess the importance of higher-order topological structure in subject FC by preserving either nodal degree or both nodal degree and strength while randomizing the specific wiring pattern. Moran surrogate tau-PET maps were additionally used to assess whether regional FC–tau associations could be explained by spatial autocorrelation in tau-PET; these surrogate maps preserved the spatial autocorrelation structure of each empirical tau-PET map while randomizing its spatial organization. (b) Comparison of explained variance when using both subject FC and template FC versus using only template FC. The scatter plots show explained variance for individual regions and subjects. More detailed comparisons are shown in the inset on the bottom-right for group ADD and in Fig. S1a–b for the other groups. Whiskers represent the 25th to 75th percentiles, and markers indicate the mean and median, computed across regions and subjects. (c) Additional explained variance from including empirical subject FC, compared to two null models where subject FC was rewired to preserve either nodal degree (Maslov and Sneppen, 2002) or both nodal degree and strength (Milisav et al., 2025). Whiskers represent the 25th to 75th percentiles, and markers indicate the mean and median, computed across regions and subjects. (d) Explained variance obtained from empirical tau-PET maps compared with Moran surrogate tau-PET maps. The upper plot summarizes group-level explained variance at the 400-region atlas resolution, and the lower plot shows results across atlas resolutions for the ADD group. Whiskers represent the 25th to 75th percentiles, and markers indicate the mean and median, computed across regions and subjects. Template FC refers to the group-derived functional connectivity matrix estimated from amyloid-negative cognitively healthy controls; subject FC was estimated separately from each participant’s rs-fMRI data. CN: cognitively normal; SCD: subjective cognitive decline; MCI: mild cognitive impairment; ADD: Alzheimer’s disease dementia; SMN: somatomotor network; DAN: dorsal attention network; VAN: ventral attention network; FPN: frontoparietal network; DMN: default mode network.

We further tested whether the regional FC–tau associations could be explained by spatial autocorrelation in tau-PET maps. For each subject and atlas, we generated Moran spectral surrogate tau-PET maps that preserved the spatial autocorrelation structure of the empirical tau-PET map while randomizing its spatial organization. Repeating the regional FC–tau regression analyses using these surrogate tau-PET maps produced explained variance values close to zero across groups and spatial scales, which were substantially lower than those obtained using empirical tau-PET maps (Fig. 1d; Fig. S1d). Thus, the observed regional associations between FC profiles and tau-PET patterns were not attributable to shared spatial smoothness alone.

These regional analyses provided the regression weights used to construct hybrid FC and linked the regional model estimates to the gains observed in the whole-brain analyses.

To assess whether these effects differed across canonical functional networks, we conducted complementary network-level analyses across atlas resolutions and identified the regions with the strongest FC–tau associations within each network (Fig. S6 and Fig. S7, respectively). Across disease stages, the Default Mode (DMN) and Frontoparietal (FPN) networks consistently showed the strongest associations with tau-PET patterns, while the limbic network was more prominent in earlier, Aβ+ groups. Performance in networks such as the visual and limbic tended to plateau or decline at finer parcellation scales, whereas the DMN and FPN remained robust. These convergent findings across regional and network analyses reinforce the central role of DMN and FPN in capturing tau-related variability, while highlighting that other networks may be more sensitive to spatial granularity.

Using the regional coefficients described above, we constructed an individualized hybrid FC representation for each subject that integrated template-level and subject-specific connectivity information without increasing the dimensionality of the whole-brain models (Fig. 2a). We then compared how well hybrid FC, template FC, and subject FC explained individual tau-PET topography in whole-brain in-sample regression models. Across the disease continuum, hybrid FC consistently outperformed both template and raw subject FC in explaining tau-PET patterns (Fig. 2b), with particularly strong effects in symptomatic groups where tau-PET burden is highest. The total explained variance by the hybrid FC model largely depended on the spatial granularity of the design: for example, in patients with ADD, the median ranged from 40% to 70% (Fig. 2c). While template FC also improved with finer parcellations (Fig. S2b), hybrid FC consistently achieved higher explanatory power across all groups and scales. In contrast, raw subject FC showed weaker performance, except in the ADD group, highlighting the stabilizing role of combining subject- and group-level information.

**Fig. 2.**
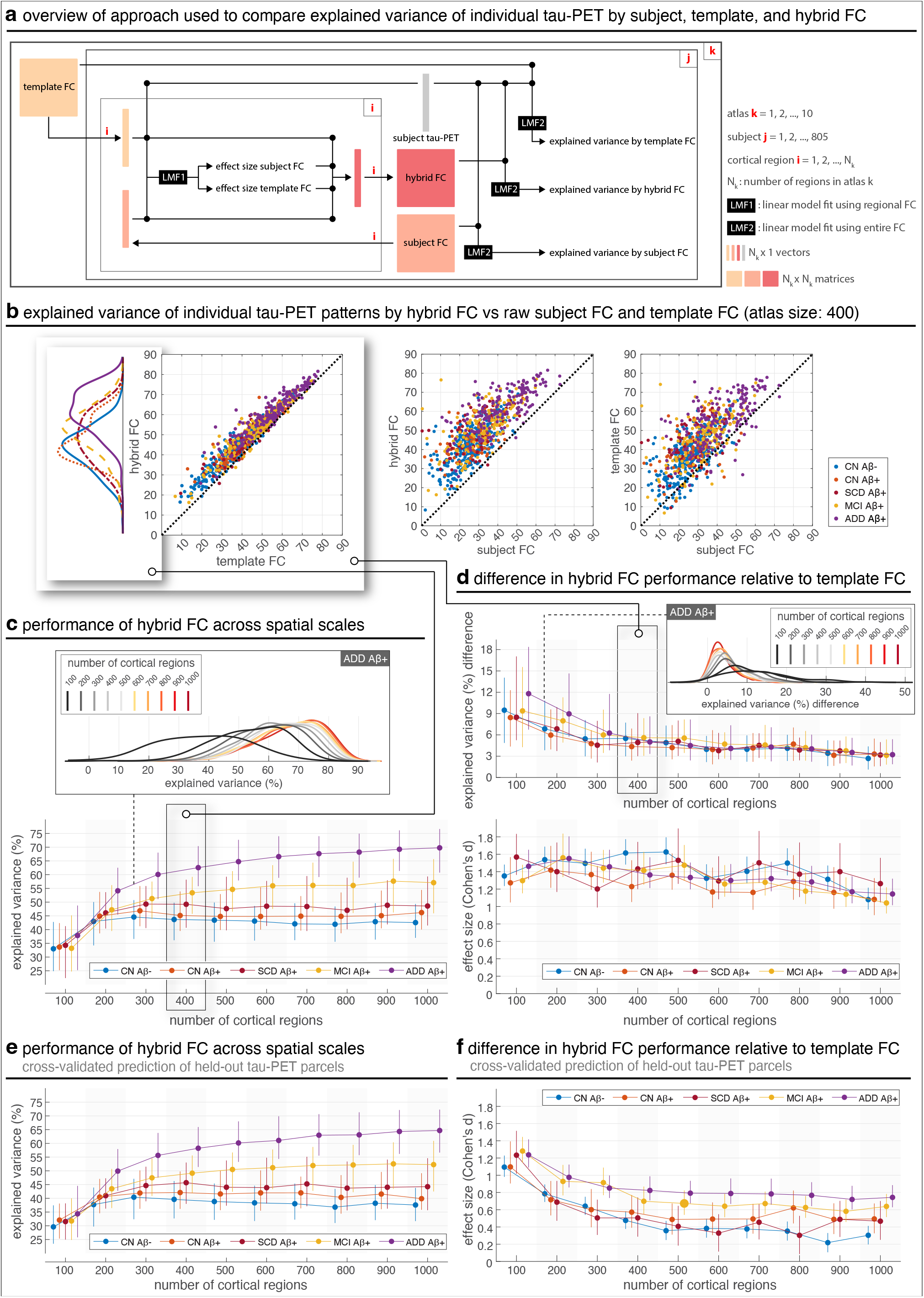
Hybrid FC explains individual tau-PET better than raw subject FC and template FC, across spatial scales. (a) Overview of the approach used to evaluate the additional explanatory power of hybrid FC beyond template FC in explaining individual tau-PET. (b) Axis values are the coefficient of determination (*R*^2^, in percentage) of individual tau-PET patterns. Axis labels specify the regressor(s) used. Each marker represents a subject. Marginal distributions for the groups are shown as kernel density plots across the different FC types. FC matrices were constructed using cortical regions defined by the Schaefer atlas (Schaefer et al., 2018) using 400 cortical regions. All reported in-sample *R*^2^ values are corrected *R*^2^ to enable comparison of models with different numbers of regressors. (c) At later stages of disease, especially MCI and ADD, in-sample explained variance using hybrid FC increased with finer-grained cortical parcellations. For the ADD group, marginal distributions of the data are compared. (d) Across clinical groups, the absolute additional in-sample explained variance of hybrid FC over template FC decreased with increasing spatial granularity, whereas the standardized effect size remained consistently large. Whiskers in the top plot represent the 25th to 75th percentiles, and markers indicate the median. Whiskers in the bottom plot show the 95% confidence interval of Cohen’s *d*. (e) Cross-validated ridge-regression prediction of held-out tau-PET parcels using hybrid FC. Performance was quantified as out-of-sample explained variance (*R*^2^), averaged across hemispheres. (f) Standardized difference in cross-validated prediction performance between hybrid FC and template FC, quantified using Cohen’s *d*. Positive values indicate better out-of-sample prediction by hybrid FC. In panels e–f, ridge models were trained and evaluated using nested, network-stratified cross-validation across held-out tau-PET parcels. Template FC refers to the group-derived functional connectivity matrix estimated from amyloid-negative cognitively healthy controls; subject FC was estimated separately from each participant’s rs-fMRI data. Hybrid FC combines template and subject-specific FC using region- and subject-specific weights. CN: cognitively normal; SCD: subjective cognitive decline; MCI: mild cognitive impairment; ADD: Alzheimer’s disease dementia.

Across groups, while the absolute difference in explained variance between hybrid and template FC gradually declined with increasing resolution (Fig. 2d, top), the difference relative to raw subject FC remained largely stable (Fig. S2c), even increasing slightly at higher resolutions in later-stage groups.

The effect size of hybrid FC’s advantage over template FC also declined somewhat with scale but remained strong across atlas sizes (Fig. 2d, bottom). Additional whole-brain comparisons showed that hybrid FC also outperformed raw subject FC, and that template FC outperformed raw subject FC, across groups and scales (Fig. S2c–f). Notably, the absolute difference in explained variance was largest at coarser parcellation resolutions and was particularly pronounced in the ADD Aβ+ group. With increasing atlas resolution, the absolute hybrid-over-template difference decreased and became more similar across clinical groups. Nevertheless, Cohen’s *d* remained consistently large across resolutions, indicating a robust advantage of hybrid FC over template FC despite the smaller absolute percentage-point differences at finer spatial scales.

Because the whole-brain FC models include many correlated predictors, we performed an additional regularized out-of-sample prediction analysis to assess whether the main model-comparison result was robust to potential overfitting and generalized to held-out tau-PET parcels. Specifically, we repeated the whole-brain analyses using ridge regression with nested cross-validation and evaluated performance using tau-PET values from held-out cortical parcels. This analysis supported the main finding: hybrid FC continued to explain individual tau-PET patterns better than template FC and subject FC alone, although out-of-sample ridge-regression estimates were, as expected, somewhat attenuated relative to the original insample estimates. Cross-validated prediction of held-out tau-PET parcels increased with spatial granularity for hybrid FC, particularly in symptomatic groups (Fig. 2e). Hybrid FC also showed consistently positive Cohen’s *d* values for the comparison with template FC across groups and atlas resolutions (Fig. 2f), indicating better out-of-sample prediction by hybrid FC, and additional comparisons with subject FC are provided in the Supplementary Information (Fig. S4).

We further assessed whether the ridge-regression coefficient patterns were stable across independent parcel subsets. These analyses were intended to assess coefficient stability of the high-dimensional models rather than to rank FC representations by *β*-stability. Split-half *β*-stability was quantified as the Pearson correlation coefficient between ridge-regression coefficient vectors estimated independently in two network-stratified halves of the cortical parcels. This analysis was performed separately for hybrid FC, template FC, and subject FC models. Across all three FC representations, *β*-stability correlations were consistently positive across clinical groups and atlas resolutions, indicating reproducible coefficient patterns under regularized estimation (Fig. S5).

Because low-magnitude BOLD FC correlations may be more susceptible to noise and physiological artifacts, we repeated the hybrid FC construction and whole-brain regression analyses after sparsifying both subject-specific and template FC matrices to retain the strongest 30% of positive connections, a threshold also used in prior related work (Franzmeier et al., 2024). Hybrid FC reconstructed from thresholded FC matrices continued to explain individual tau-PET patterns better than thresholded template FC and thresholded subject FC across clinical groups and atlas resolutions (Fig. S3). However, the hybrid-over-template effect sizes were more conservative than in the unthresholded primary analysis, suggesting that hard edge removal may discard distributed connectivity information that remains useful within the hybrid FC framework. These findings show that the main conclusion is robust to a commonly used FC thresholding approach, while supporting the use of unthresholded FC in the primary analyses.

These results demonstrate that hybrid FC provides a robust and scalable framework for capturing both canonical and individualized contributions to tau-PET patterns, particularly in later disease stages.

### 2.B. Hybrid FC better explains individual tau-PET than cohort-level canonical tau-PET patterns or individual Aβ-PET patterns, alone or combined

The results thus far have demonstrated that individual FC provides additional explanatory power beyond group-level template FC in explaining individual tau-PET patterns. We next sought to determine whether, and to what extent, FC captures unique aspects of individual tau-PET patterns beyond what can be explained by canonical tau-PET patterns, addressing a dimension that has received limited attention in previous studies. Here, canonical patterns refer to stereotypical spatial distributions of tau-PET signal across the cortex, which we derived by estimating ensemble off- and on-target tau-PET retention patterns present across the entire cohort via Gaussian mixture modelling (Fig. S8–S10). These patterns represent generalizable features of tau accumulation, distilled across individuals, and serve as a reference or baseline model of tau-PET topography.

When comparing model performance, we found that for Aβ+ individuals with MCI or ADD, hybrid FC explained individual tau-PET patterns better than canonical tau-PET patterns, while in early-stage patients, whose tau pathology is more spatially restricted, canonical tau-PET patterns provided a better explanation (Fig. 3a). However, this distinction shifted when we increased the spatial resolution of the analysis, i.e., using more fine-grained cortical regions (Fig. 3b). First, at higher resolutions of cortical parcellation (>300 regions), hybrid FC more distinctly outperformed canonical patterns in Aβ+ patients with MCI or ADD, and second, it matched or outperformed canonical patterns even in early disease stages. These findings suggest that individual FC may capture subtle, spatially localized features of tau distribution that are not reflected in cohort-level canonical patterns, particularly when analysed at higher spatial granularity. Furthermore, across all individuals, combining canonical patterns with hybrid FC explained more variance in individual tau-PET patterns than either factor alone, suggesting that the two provide complementary information about the spatial distribution of tau pathology (Fig. 3a, top-right). The combined model including hybrid FC together with canonical tau-PET patterns showed the highest corrected *R*^2^ among the models considered. Because this model includes additional predictors, we interpret this result as evidence that hybrid FC and canonical tau-PET patterns contain partially complementary explanatory information, rather than as a simple model-size advantage. Importantly, corrected *R*^2^ penalizes the additional predictors, so the increase indicates that canonical tau-PET maps contributed residual explanatory information beyond hybrid FC. These trends were further supported by results across spatial scales (Fig. S11). While canonical tau-PET patterns (especially off-target binding) consistently declined in explanatory power with increasing atlas resolution (Fig. S11a–c), hybrid FC performance increased (Fig. 2c), eventually surpassing the canonical models at finer scales. Direct comparisons confirmed that the added value of hybrid FC over canonical patterns grew with spatial granularity, particularly in Aβ+ individuals with MCI and ADD (Fig. S11e–f). These findings reinforce that canonical tau-PET patterns and individual FC capture different aspects of individual tau-PET, and that the benefit of FC-based modelling becomes increasingly evident when leveraging fine-grained spatial information.

**Fig. 3.**
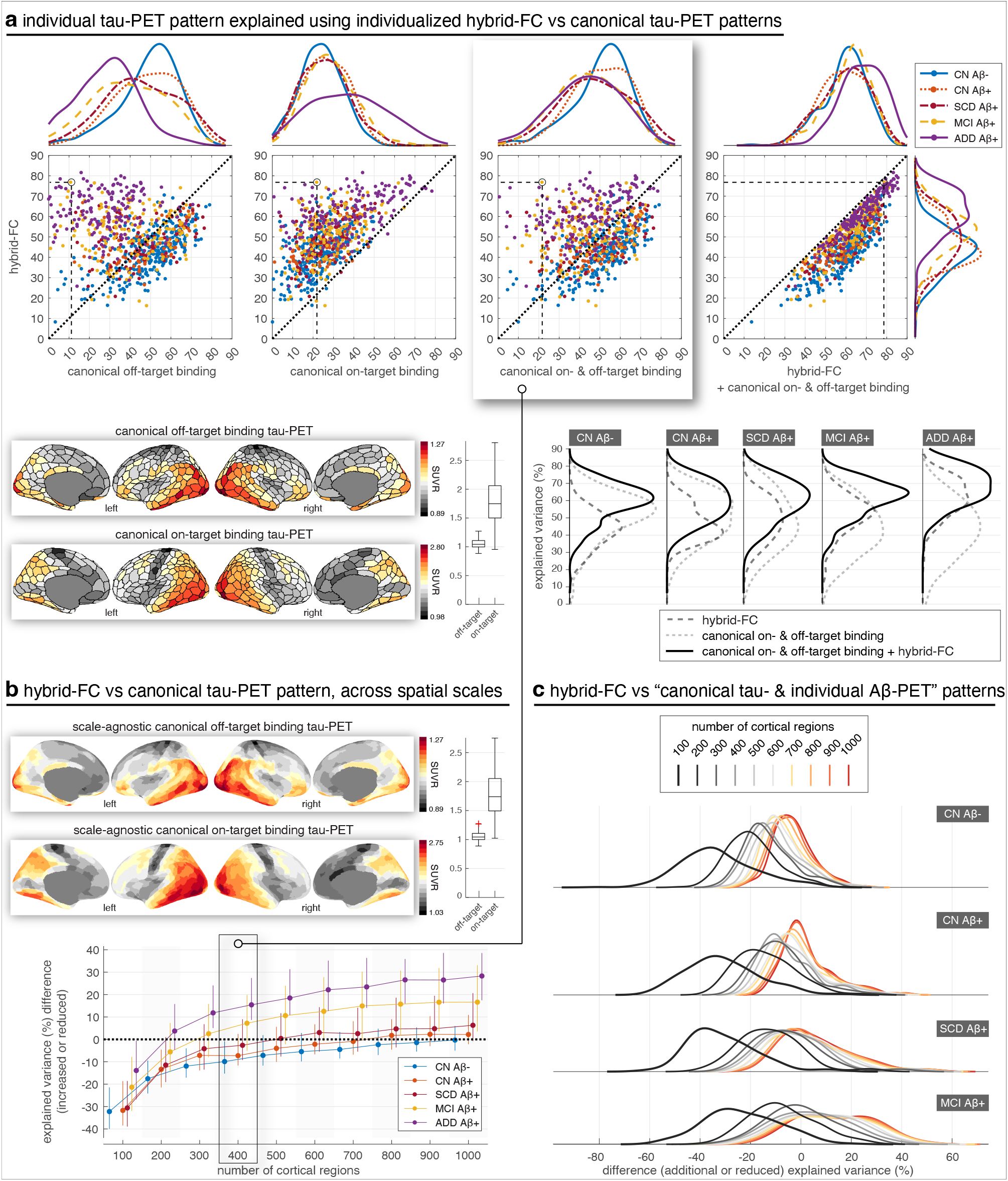
Hybrid FC explains individual tau-PET better than canonical tau-PET patterns, across spatial scales. (a) Axis values are explained variance (corrected *R*^2^, in percentage) of individual tau-PET. Axis labels specify regressor(s) used. Each marker represents a subject; e.g. for the circled Aβ+ patient with MCI, hybrid FC (y-axis across all 4 panels) explains 77% of the spatial variance in their tau-PET pattern whereas canonical off-(1st panel, x-axis) and on-target (2nd panel, x-axis) binding explain 11% and 22% of the variance, respectively; canonical off- and on-target binding together (3rd panel, x-axis) explain 22% of the variance whereas if hybrid FC is also used (4th panel, x-axis) the explained variance increases to 79%. Marginal distributions for the groups are shown as kernel density plots; in the bottom plot, marginal distributions shown next to scatter plots (top panel) are re-configured (bottom panel) to enable comparison of kernel densities associated to each group for the different regressors. FC matrices were constructed using cortical regions defined by Schaefer atlas (Schaefer et al., 2018) using 400 cortical regions. All reported *R*^2^ are corrected *R*^2^ to facilitate comparison of models with different numbers of regressors across this study. (b) The performance of hybrid FC over canonical tau-PET patterns further improves as a function of the resolution of FC, where FC built using more fine-grained functional regions performs better. The cortical projections show the scale-agnostic version of the off- and on-target tau PET binding patterns obtained by averaging the binding patterns derived for the ten different parcellation scales; the off- and on-target tau binding patterns for each spatial resolution are shown in Fig. S9 and Fig. S10 in the supplementary material, respectively. Whiskers represent the 25^th^ to 75th percentiles; markers indicate the median. The marked panel (i.e. for 400 regions) relates to data shown in the third, top panel in (a). (c) The difference in explained variance of individual tau-PET by hybrid FC relative to canonical tau-PET patterns combined with individual Aβ-PET is shown across spatial scales. The difference in performance increases as the spatial resolution increases across the three groups, with hybrid FC showing improved performance (values above zero) most dominantly in Aβ+ patients with MCI and at high spatial resolutions (parcellations with 400 cortical regions and above). The differences are generally greater in Aβ+ patients with MCI. Template FC refers to the group-derived functional connectivity matrix estimated from amyloid-negative cognitively healthy controls. Hybrid FC combines this template FC representation with each participant’s subject-specific FC using region- and subject-specific weights. CN: cognitively normal; SCD: subjective cognitive decline; MCI: mild cognitive impairment; ADD: Alzheimer’s disease dementia.

For CN individuals and patients with MCI who had Aβ-PET scans available, we further explored whether using individual Aβ-PET patterns as regressors in addition to canonical tau-PET patterns improves performance relative to that provided by hybrid FC. The results revealed that individual Aβ-PET patterns explained individual tau-PET patterns only minimally, compared to hybrid FC and canonical tau-PET patterns (Fig. S15). Even when combined with canonical tau-PET patterns they only marginally increased the performance in Aβ+ patients with MCI while the performance for the majority of Aβ+ patients with MCI remained below that of hybrid FC (Fig. 3c). Together, these results underscore the unique and complementary value of individualized FC in capturing personalized features of tau pathology that are not fully accounted for by canonical tau or amyloid-based models.

### 2.C. External replication of hybrid FC performance in ADNI and A4

To test whether the main findings generalized beyond BioFINDER, we performed external replication analyses in two independent cohorts: ADNI and A4. ADNI provided an independent cohort of 361 participants across cognitively normal, MCI, and AD dementia stages, including 149 CN A*β*-, 91 CN A*β*+, 78 MCI A*β*+, and 43 ADD A*β*+ participants. A4 provided an independent cognitively unimpaired cohort enriched for preclinical AD, including 36 CN A*β*- and 300 CN A*β*+ participants. Demographic and clinical characteristics of the ADNI and A4 replication samples are shown in Tables 2 and 3. Following the BioFINDER workflow, the replication analyses used subject-space cortical parcellations of tau-PET and resting-state fMRI data and repeated the primary model-comparison analyses across atlas resolutions. Given the breadth of analyses performed in the primary BioFINDER cohort, the replication analyses focused on the main model-comparison findings: whether hybrid FC explained individual tau-PET patterns better than template FC, and, in ADNI, whether hybrid FC outperformed canonical tau-PET patterns.

**Table 2.**
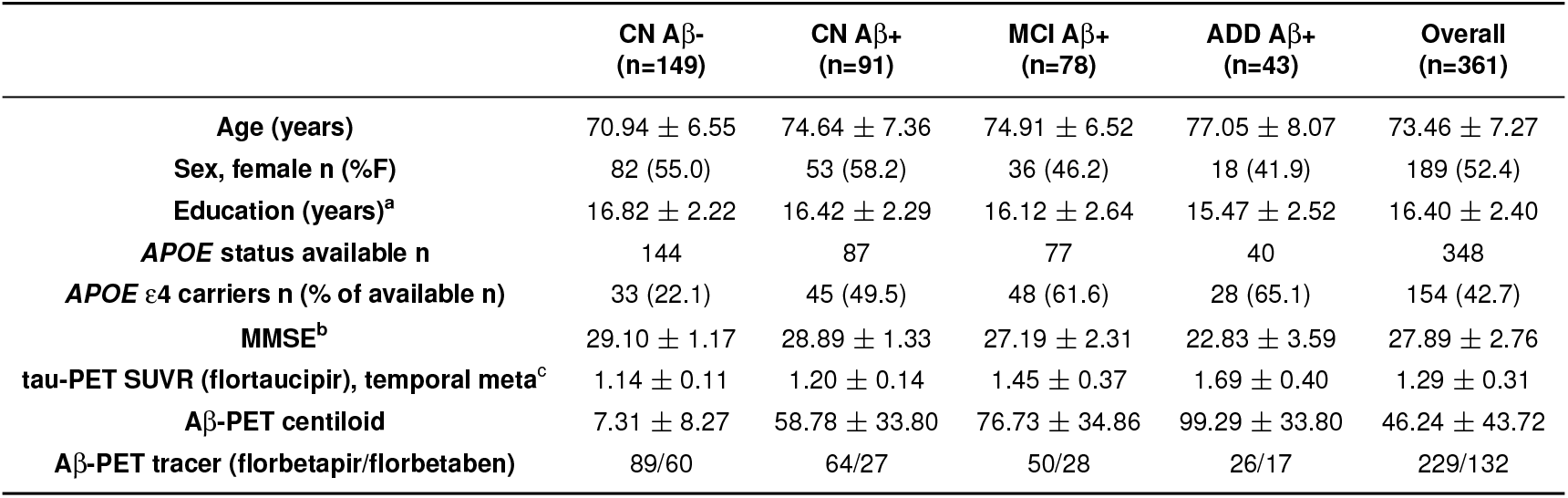
Demographics and clinical characteristics of the ADNI sample. Aβ: β-amyloid; Aβ status (positive: Aβ+, negative: Aβ−) was determined based on global amyloid-PET SUVRs using tracer-specific cut-offs at 1.11/1.08 for Florbetapir/Florbetaben, as previously established in the ADNI cohort (Landau et al., 2012; Royse et al., 2021). Data are presented as mean *±* s.d. unless specified otherwise. CN: Cognitively normal; MCI: mild cognitive impairment; ADD: Alzheimer’s disease dementia; MMSE: mini mental state examination; *APOE* ε4: apolipoprotein E genotype carrying at least one ε4 allele. ^a^There are 1, 0, 0, 0 missing values in the respective categories. ^b^There are 2, 1, 1, 1 missing values in the respective categories. ^c^There are 9, 5, 2, 0 missing values in the respective categories.

**Table 3.** Demographics and clinical characteristics of the A4 sample. Aβ: β-amyloid; Aβ status (positive: Aβ+, negative: Aβ−) was determined based on a composite amyloid-PET SUVR at cut-off 1.15 as previously suggested by the A4 imaging core (Langford et al., 2020). Data are presented as mean *±* s.d. unless specified otherwise. CN: Cognitively normal; MMSE: mini mental state examination; *APOE* ε4: apolipoprotein E genotype carrying at least one ε4 allele.

|  | CN Aβ-<br>(n=36) | CN Aβ+<br>(n=300) | Overall<br>(n=336) |
| --- | --- | --- | --- |
| Age (years) | 70.12 ± 4.23 | 72.43 ± 4.90 | 72.18 ± 4.88 |
| Sex, female n (%F) | 21 (58.3) | 172 (57.3) | 193 (57.4) |
| Education (years) | 16.31 ± 2.87 | 16.23 ± 2.75 | 16.24 ± 2.76 |
| APOE status available n | 36 | 300 | 336 |
| APOE ε4 carriers n (% of total) | 9 (25.0) | 182 (60.7) | 191 (56.8) |
| MMSE | 28.69 ± 1.45 | 28.59 ± 1.26 | 28.60 ± 1.28 |
| Aβ-PET centiloid (florbetapir) | 20.61 ± 7.75 | 68.07 ± 29.14 | 62.99 ± 31.31 |

In both ADNI and A4, hybrid FC explained individual tau-PET patterns better than template FC across atlas resolutions (Fig. 4a). This effect was evident at the 400-region atlas resolution in the subject-level scatter plots and remained positive across spatial scales, as reflected by consistently positive Cohen’s *d* values for the hybrid FC versus template FC comparison. In ADNI, hybrid FC explained variance increased with spatial granularity and was highest in symptomatic Aβ+ groups, consistent with the pattern observed in BioFINDER. The corresponding supplementary analyses further showed that hybrid FC also outperformed subject FC, while template FC outper-formed subject FC, in both ADNI and A4 (Figs. S12 and S13). Thus, the overall hierarchy of FC representations observed in BioFINDER—hybrid FC *>* template FC *>* subject FC—was reproduced in both external cohorts.

**Fig. 4.**
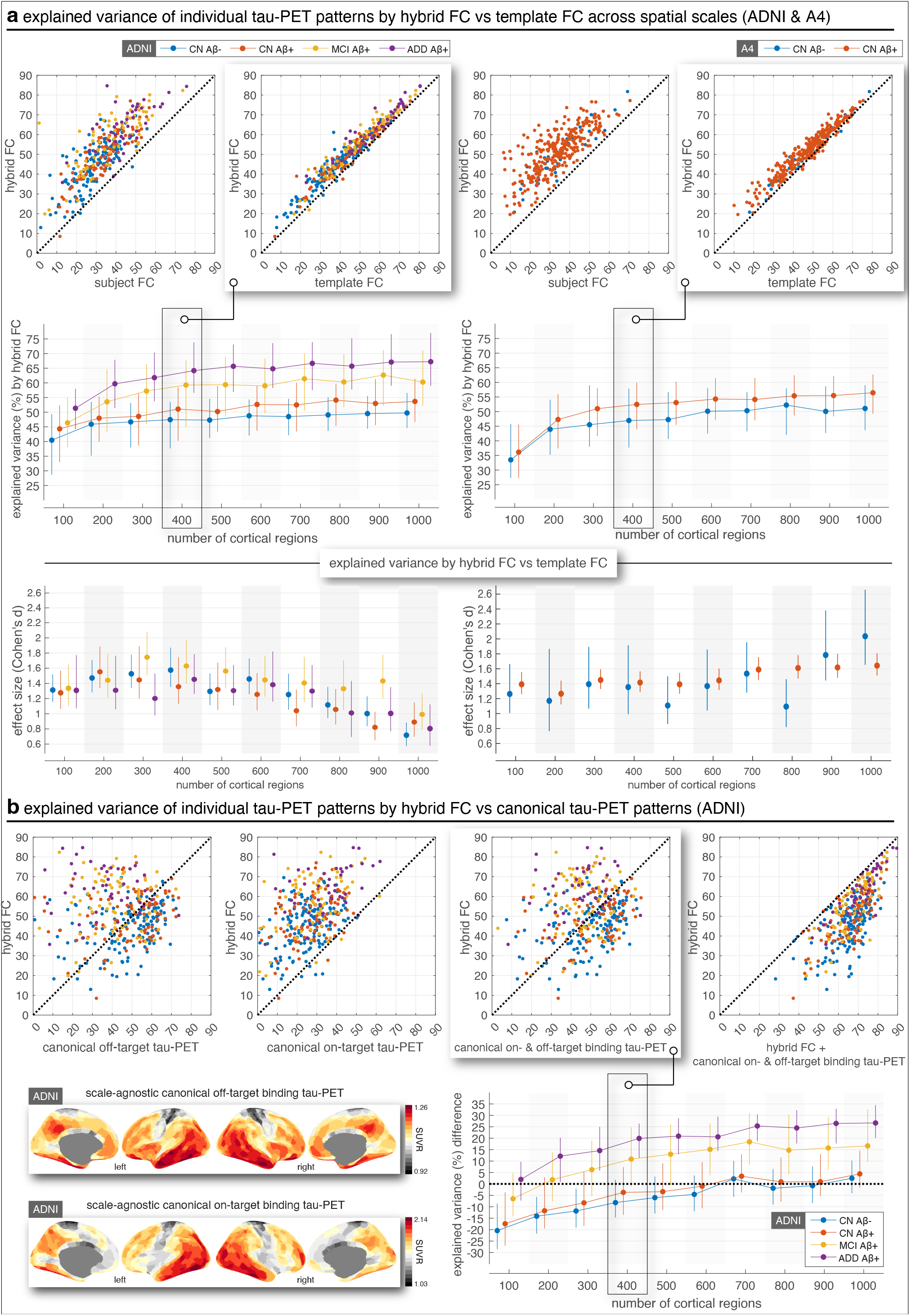
External replication of hybrid FC performance in ADNI and A4. (a) Explained variance of individual tau-PET patterns by hybrid FC compared with subject FC and template FC in two independent cohorts. Scatter plots show individual-subject explained variance at the 400-region atlas resolution. Line plots show hybrid FC explained variance across atlas resolutions and the standardized difference in explained variance between hybrid FC and template FC, quantified using Cohen’s *d*. Positive Cohen’s *d* values indicate greater explained variance by hybrid FC than template FC. (b) Replication of the comparison between hybrid FC and canonical tau-PET patterns in ADNI. Scatter plots show individual-subject explained variance at the 400-region atlas resolution for hybrid FC compared with canonical off-target binding tau-PET, canonical on-target binding tau-PET, canonical on- and off-target binding tau-PET, and the combined model including hybrid FC together with canonical on- and off-target binding tau-PET. Surface maps show the scale-agnostic canonical off-target and on-target binding tau-PET patterns estimated in ADNI. The line plot shows the difference in explained variance between hybrid FC and canonical on- and off-target binding tau-PET across atlas resolutions. Positive values indicate greater explained variance by hybrid FC. Points and error bars show group-level mean *±* 95% confidence interval across subjects for each clinical group and atlas resolution. In ADNI, groups included CN A*β*-(n=149), CN A*β*+ (n=91), MCI A*β*+ (n=78), and ADD A*β*+ (n=43). In A4, groups included CN A*β*-(n=36) and CN A*β*+ (n=300). Template FC refers to the group-derived functional connectivity matrix estimated from amyloid-negative cognitively healthy controls; subject FC was estimated separately from each participant’s rs-fMRI data. Hybrid FC combines template and subject-specific FC using region- and subject-specific weights. CN: cognitively normal; MCI: mild cognitive impairment; ADD: Alzheimer’s disease dementia.

We additionally tested whether hybrid FC outperformed canonical tau-PET patterns in ADNI. Hybrid FC outperformed canonical on-target binding tau-PET patterns across clinical groups and atlas resolutions (Fig. 4b; Fig. S14). Compared with canonical off-target binding tau-PET patterns, hybrid FC showed the clearest advantage in MCI Aβ+ and ADD Aβ+ groups, whereas differences were smaller or near zero in cognitively normal groups at finer resolutions. Similar disease-stage-dependent effects were observed when hybrid FC was compared with the combined canonical on- and off-target binding tau-PET model. We did not perform this canonical tau-PET comparison in A4 because, in contrast to the ADNI cohort, the A4 cohort only included cognitively unimpaired participants with low tau burden. Together, these external replication analyses indicate that the central BioFINDER findings replicate across independent datasets and support the generalizability of hybrid FC as a representation of individual tau-PET topography.

### 2.D. Hybrid FC better captures individual tau-PET but not Aβ-PET compared to canonical PET patterns

The key focus of this study has been to understand the explanatory power of FC in relation to individual tau-PET patterns, an endeavour motivated by the hypothesis that tau pathology spreads along neuronal communication pathways. To test the specificity of this hypothesis for tau pathology, we examined whether, and if so to what degree, FC contributes to explaining individual Aβ-PET patterns, for which this hypothesis is less commonly proposed. We therefore repeated the analysis with Aβ-PET data and compared the performance to the tau-PET analysis within the same individuals; patients with ADD were excluded as they lacked Aβ-PET data. The superior explanatory power of hybrid FC over canonical PET patterns we observed for tau-PET was not observed for Aβ-PET (Fig. 5a). Compared to the explanatory power of hybrid FC, Aβ-PET patterns were better explained by the estimated canonical off-target binding Aβ-PET pattern in Aβ-individuals, and by the canonical on- target binding pattern in Aβ+ patients. The apparent advantage of hybrid FC over the canonical off-target binding pattern in Aβ-positive individuals should therefore not be interpreted as evidence that FC strongly explains Aβ deposition. Rather, this comparison reflects that the off-target binding pattern primarily captures non-specific signal relevant to Aβ-negative individuals and is not expected to capture the dominant on-target cortical Aβ signal present in Aβ-positive individuals. For Aβ-positive groups, the relevant canonical comparator is therefore the on-target, or combined on-/off-target, Aβ-PET pattern. This contrasts with observations for individual tau-PET patterns.

**Fig. 5.**
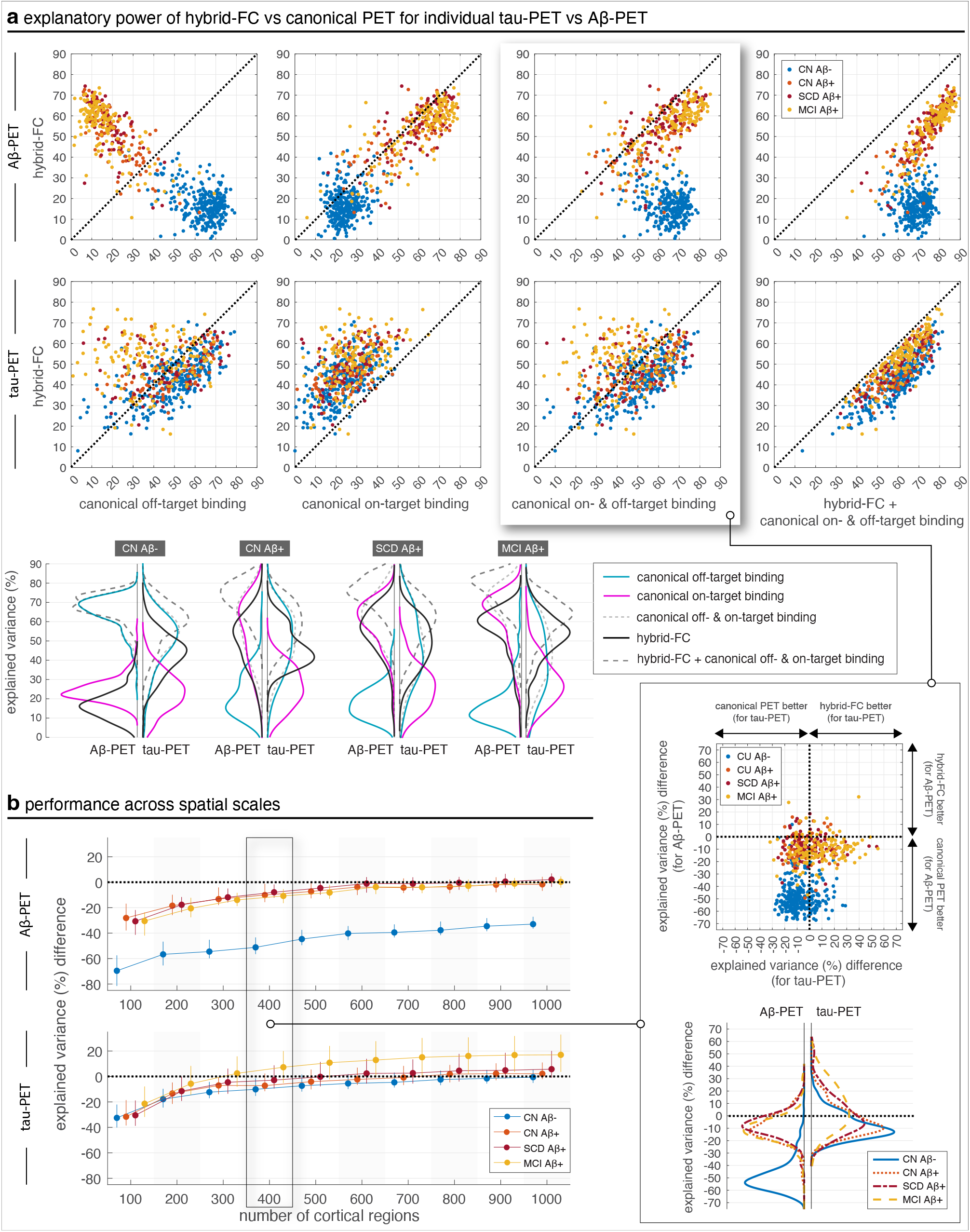
Hybrid FC explains individual tau-PET, but not Aβ-PET, better than canonical PET patterns. (a) In the first row, linear models are fit to individual Aβ-PET patterns using different regressors as specified by axes labels of each scatter plot. Each marker represents a subject and axes values are the percentage of explained variance (corrected *R*^2^ resulting from linear fit models). In the second row, linear fit models are instead fitted to individual tau-PET patterns. Marginal distributions of explained variance are compared across groups. Note that in each group explained variances are being compared for tau-PET and Aβ-PET which are different in nature in terms of the amount of spatial content they entail (Aβ-PET patterns generally exhibit less spatial complexity compared to tau-PET patterns), and, as such, comparison between the two should be made in relation to difference in performance using different regressors for each data modality (i.e., up-down shift in kernel densities for each modality) rather than comparing explained variance of tau-PET to that of Aβ-PET. (b) Explained variance using hybrid FC minus explained variance using canonical PET patterns across scales. Differences in explained variance for the data in the first, second, and fourth columns in (a), across spatial scales, are shown in Fig. S16, whereas the absolute explained variances are shown in Fig. S17. Whiskers represent the 25th to 75th percentiles; markers indicate the median. The marked panel (i.e. for 400 regions) relates to data shown in (a). Template FC refers to the group-derived functional connectivity matrix estimated from amyloid-negative cognitively healthy controls. Hybrid FC combines this template FC representation with each participant’s subject-specific FC using region- and subject-specific weights. CN: cognitively normal; SCD: subjective cognitive decline; MCI: mild cognitive impairment.

To further contextualize these findings, we compared marginal distributions of explained variance. This revealed that hybrid FC largely outperforms canonical patterns for tau-PET, but not for Aβ-PET (Fig. 5a, bottom-right inset). The difference increased as the spatial granularity of the analysis increased (Fig. 5b). For Aβ-CN and Aβ+ patients with MCI, the difference in explained variance was consistently greater for tau-PET compared to Aβ-PET, whereas for Aβ+ CN the performances were on par. Furthermore, comparison of differences in explained variances for other settings (e.g. hybrid FC minus on-target binding pattern) show that canonical patterns explain more variance in Aβ-PET than hybrid FC, especially in Aβ+ CN and patients with SCD or MCI (Fig. S16), while the reverse is true for tau-PET. This trend is reinforced by the absolute explained variances (Fig. S17), where canonical Aβ-PET patterns consistently explain more variance than hybrid FC, in contrast to tau-PET where hybrid FC explains the most variance across all groups and scales. This analysis supports a modality-specific dissociation: FC-based models explained tau-PET patterns better than canonical PET patterns, particularly at finer spatial scales, whereas Aβ-PET patterns were better explained by cohort-derived canonical PET patterns included in the same modelling framework.

### 2.E. Longitudinal tau-PET is better explained by hybrid FC than template FC

For a subset of patients with available follow-up tau-PET scans in BioFINDER (52% of patients, see Table 1), we examined the extent to which hybrid FC derived from baseline data explained the individual’s follow-up tau-PET patterns. Hybrid FC explained follow-up tau-PET with greater explained variance at finer spatial scales, plateauing around 400—600 regions (Fig. 6a), and consistently outperformed template FC across all clinical groups and spatial resolutions (Fig. 6b). Both baseline template FC and baseline hybrid FC explained variance in follow-up tau-PET patterns. However, hybrid FC consistently outperformed template FC across clinical groups and atlas resolutions (Fig. 6b). Because hybrid FC incorporates the subject-specific FC component in addition to template FC, this result indicates that individualized baseline connectivity provides additional information about future tau topography beyond the group-level template connectivity pattern. While Fig. 6a–b focus on follow-up tau-PET, we confirmed that hybrid FC achieved similar levels of explained variance for baseline tau-PET patterns in groups consisting of the same matching individuals (Fig. S18a), reinforcing the model’s robustness across time-points. Moreover, hybrid FC provided additional explanatory value over template FC, particularly at finer spatial resolutions and in later disease stages (Fig. S18b). However, despite this group-level consistency, individual-level comparisons revealed substantial variability: the correlation between how well hybrid FC explained baseline versus follow-up tau-PET patterns was modest (Fig. S18c, left), with standardized effect sizes peaking at intermediate spatial resolutions (Fig. S18c, middle). This suggests that, while hybrid FC captures stable spatial features at the cohort level, individual FC–tau correspondence varies over time, potentially reflecting heterogeneity in longitudinal tau-PET change.

**Fig. 6.**
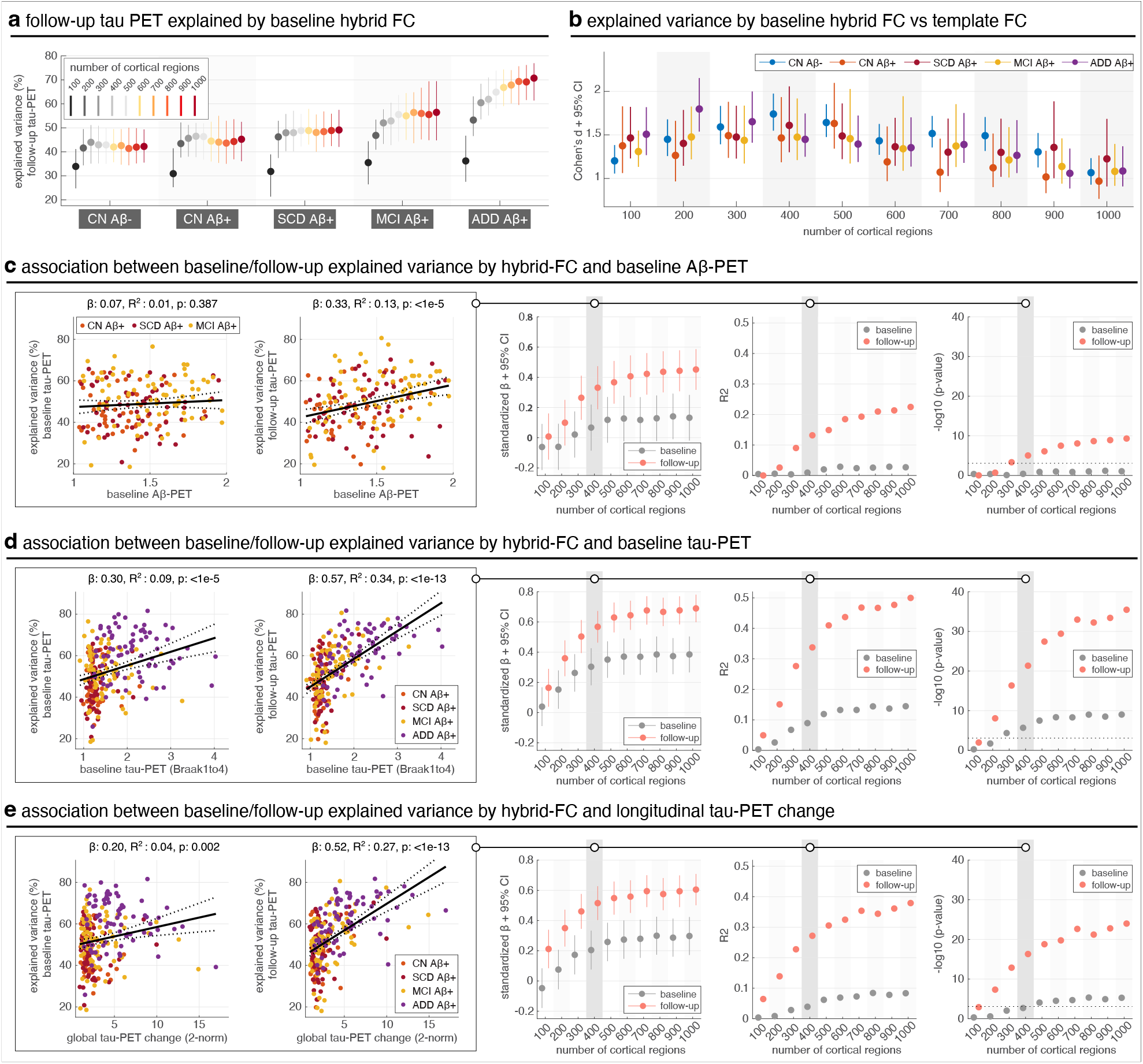
Explained variance of individual follow-up tau-PET patterns using baseline hybrid FC. (a) Baseline hybrid FC explained individual follow-up tau-PET patterns across clinical groups, with performance increasing at finer spatial scales and plateauing around 400–600 cortical regions. (b) Across all scales and groups, hybrid FC outperformed template FC in explaining follow-up tau-PET, as quantified by Cohen’s *d*. (c–e) Associations between tau-PET model performance (explained variance, EV) and baseline biomarkers: global Aβ-PET SUVR (c), baseline tau-PET (Braak I–IV) (d), and longitudinal tau-PET change (Braak I–IV) (e); for Aβ-PET, the meta ROI SUVR is the average SUVR calculated from a global neocortical ROI, including prefrontal, lateral temporal, parietal, anterior cingulate and posterior cingulate/precuneus. For each biomarker, we modelled its relationship with tau-PET EV at both baseline and follow-up using linear regression with age and sex as covariates. Predictors and outcomes were z-scored, and standardized beta coefficients (with 95% CI), *R*^2^, and p-values were extracted across spatial scales. Leftmost plots in each panel show associations for one representative spatial scale (400 regions); rightmost columns summarize results across scales. Analyses were restricted to Aβ-positive individuals (CN Aβ+, SCD Aβ+, MCI Aβ+, ADD Aβ+), except for Aβ-PET analyses (panel c), where the ADD group was excluded due to missing Aβ-PET data. The CN Aβ-group was excluded from all analyses in panels c-e to avoid skewing the regression due to consistently low biomarker levels in this group. Explained variance refers to corrected *R*^2^. Template FC refers to the group-derived functional connectivity matrix estimated from amyloid-negative cognitively healthy controls. Hybrid FC combines this template FC representation with each participant’s subject-specific FC using region- and subject-specific weights.

To better understand the factors shaping this heterogeneity, we next examined whether baseline aggregate pathological markers were associated with the model’s ability to explain tau-PET patterns at either timepoint. Explained variance at follow-up was only weakly associated with baseline Aβ-PET levels (Fig. 6c, left), and this relationship was not significant at baseline (Fig. 6c, middle/right). In contrast, baseline tau-PET levels were more strongly associated with the explained variance of follow-up tau-PET, and this relationship was stronger than at baseline across most spatial scales (Fig. 6d). These patterns suggest that hybrid FC captures aspects of functional architecture that are associated with the spatial distribution of subsequent tau-PET signal. To clarify why the association between baseline aggregate tau burden and explained variance was stronger at follow-up, we tested baseline Aβ-PET as a mediator across spatial scales (Fig. S19). Mediation was evident only for follow-up, becoming significant at higher spatial scales, and was absent for baseline at all scales. In line with this interpretation, subjects who showed greater longitudinal changes in tau-PET (i.e., greater tau spread) also exhibited higher follow-up explained variance (Fig. 6e), again more so than at baseline, and with the effect increasing and stabilizing across spatial granularity. Together, these findings support the potential utility of hybrid FC for estimating follow-up tau-PET patterns and are consistent with, but do not establish, network-based models of tau propagation. By linking baseline individual FC to follow-up tau-PET topography, our results highlight an association between baseline connectivity and later tau-PET spatial patterns.

The explanatory performance of hybrid FC varied substantially across individuals, indicating that FC–tau correspondence is not uniform even within clinical groups. To examine potential contributors to this variability, we related hybrid FC explained variance to baseline Aβ-PET, baseline tau-PET burden, longitudinal tau-PET change, and age. Baseline tau burden and longitudinal tau-PET change were associated with higher explained variance, particularly for follow-up tau-PET patterns (Fig. 6d–e), whereas baseline Aβ-PET showed only a weak association with follow-up explained variance and no significant association at baseline (Fig. 6c). Age was not associated with baseline explained variance but showed a small negative association with follow-up explained variance (*β* = − 0.15, *R*^2^ = 0.02, *p* = 0.022; Fig. S18d). Together, these analyses indicate that inter-individual variability in FC-based explanatory performance is partly related to pathology burden and longitudinal tau change, with only a limited contribution of age.

## 3. Discussion

Network-based models of tau propagation propose that the spatial distribution of pathology in Alzheimer’s disease (AD) is related to the brain’s intrinsic connectivity architecture, yet direct evidence at the individual level has been limited. We hypothesized that subject-specific functional connectivity (FC) would account for individual differences in the spatial distribution of tau pathology, such that individual differences in connectivity profiles would account for corresponding differences in tau-PET patterns. Using 1227 tau-PET scans from 805 participants in the BioFINDER-2 study, we found that individual-level FC indeed improved the explanation of tau-PET topographies compared to group-level FC alone, indicating that both shared and idiosyncratic features of network architecture provide complementary explanatory information about tau-PET topography. These effects were particularly evident in symptomatic individuals, where tau progression is most advanced, and at higher spatial resolution. Furthermore, baseline FC explained follow-up tau-PET patterns, suggesting that individualized baseline connectivity contains information relevant to later tau-PET topography and may inform future models of tau-PET progression. Taken together, these findings are consistent with the network-spread hypothesis and extend previous group-level observations to individualized FC estimates: idiosyncrasies in individual network architecture are associated with variability in tau distribution, above and beyond what can be explained by canonical connectivity or PET patterns. This work thus enhances our understanding of tau PET heterogeneity in AD, underscoring that individual variability in functional brain architecture contains information relevant to heterogeneity in tau-PET patterns across individuals.

The principal contribution of this study is methodological. We evaluated how alternative FC representations and parcellation resolutions influence the extent to which individual tau-PET topography can be statistically explained. The observed associations should therefore be interpreted as model-based correspondence between FC and tau-PET patterns, rather than as direct evidence that functional connectivity causally determines tau propagation. Although the findings are consistent with network-based accounts of tau distribution, the present observational analyses cannot establish the underlying biological mechanism.

Our results are consistent with, and add an individualized modelling perspective to, the prion-like propagation hypothesis of tau pathology, which posits that tau spreads transneuronally along functional and structural pathways. While previous studies have shown that group-averaged connectomes strongly explain stereotypical patterns of tau distribution (Franzmeier et al., 2020; Vogel et al., 2020), these models do not accurately capture subject-specific heterogeneity of tau deposition. Here, we show that individualized FC provides explanatory information about regional tau-PET patterns beyond that available from the normative template FC (Fig. 1), and that hybrid FC improves whole-brain explanation of individual tau-PET topography beyond template FC alone (Fig. 2), especially in later disease stages. The greater explanatory performance of empirical subject FC relative to degree- and strength-preserving null models indicates that the findings were not attributable solely to lower-order graph properties. Complementary Moran surrogate analyses further showed that the regional FC–tau associations were not explained by spatial autocorrelation of tau-PET maps alone (Fig. 1c–d; Fig. S1). These findings underscore the importance of considering personalized connectomic architecture to understand individual differences in tau distribution, building on fMRI-based evidence that patients with AD show individual-specific functional connectivity profiles (Millar et al., 2020; Stampacchia et al., 2024), with variability across networks such as the default mode, salience, and frontoparietal control (Adriaanse et al., 2014; Gour et al., 2014; Lehmann et al., 2015; Lin et al., 2018; Franzmeier et al., 2018; Tu et al., 2024). Together, our results highlight that variability in brain connectivity contains information relevant to individual differences in tau-PET topography.

The external replication analyses indicate that the main findings generalized across cohorts. Absolute explained-variance values differed across cohorts: hybrid FC *R*^2^ values were broadly comparable between BioFINDER and ADNI, reaching approximately 60–70% in symptomatic Aβ+ groups at finer atlas resolutions, whereas A4 showed a more compressed range, generally around 45–55%, consistent with its cognitively unimpaired, lower-tau disease-stage composition. Despite these cohort differences in absolute *R*^2^, the relative model hierarchy was largely preserved: hybrid FC consistently explained more individual tau-PET variance than template FC, and, in ADNI, also exceeded canonical tau-PET patterns. This distinction is important because absolute *R*^2^ values are likely to depend on the amount and spatial heterogeneity of tau pathology present in a cohort, whereas the model comparison tests whether individualized FC adds information beyond group-level reference patterns. The replication therefore supports the robustness of the central conclusion while also showing that the magnitude of FC–tau correspondence is partly cohort- and disease-stage dependent.

Notably, hybrid FC explained tau-PET patterns better than canonical PET patterns, but this pattern was not observed for Aβ-PET (Fig. 5a–b), suggesting relative specificity to tau, and reinforcing the notion that FC is more tightly linked to tau-PET topography than to Aβ-PET topography. Furthermore, FC explained individual tau-PET heterogeneity in later disease stages better than cohort-derived canonical tau-PET patterns (Fig. 3b), even when the latter were combined with individual Aβ-PET data (Fig. 3c). These findings raise important questions about upstream modulators of FC, such as Aβ pathology, which accumulates early in the disease and may be associated with later FC–tau correspondence. Consistent with this, we found that baseline Aβ-PET levels were more strongly associated with the ability of baseline FC to explain follow-up tau-PET patterns than concurrent tau-PET patterns, even though both tau scans were explained using the same FC data (Fig. 6c). This suggests that Aβ-related differences in functional connectivity may be associated with later FC–tau correspondence. Mediation analyses revealed that baseline Aβ explains part of the association between early tau burden and future FC-constrained tau patterns, but not concurrent patterns, with the mediated component becoming more apparent at finer parcellations (Fig. S19). This scale dependence suggests that Aβ-related influences on tau PET topography are expressed at meso-scale functional architecture captured by higher-resolution FC, while a substantial direct effect of baseline tau remains. Notably, this effect was not driven by evolving FC profiles, but rather by baseline Aβ influencing the extent to which baseline FC explained longitudinal changes in tau burden. These findings corroborate prior work showing that Aβ deposition can induce functional hyper-connectivity or dysregulation that facilitates tau spread (Jones et al., 2017; Palmqvist et al., 2017; Franzmeier et al., 2020; Pereira et al., 2019; Roemer-Cassiano et al., 2025), and they extend this notion by suggesting a temporally ordered association among baseline Aβ burden, FC, and later tau-PET topography. While our analyses focused on static FC, future efforts integrating subject-specific Aβ maps and dynamic FC may help determine whether Aβ actively modifies the functional connectome to accelerate tau transmission. Together, these findings support further investigation of individualized models linking Aβ burden, baseline tau burden, FC, and longitudinal tau-PET patterns.

The large inter-individual variability in hybrid-FC explained variance also suggests that the strength of FC–tau correspondence differs across individuals (Fig. 6a; Fig. S18a–c). This variability was not random with respect to disease biology: individuals with higher baseline tau burden and greater longitudinal tau-PET change tended to show higher explained variance, particularly for follow-up tau-PET patterns (Fig. 6d–e). Baseline Aβ-PET showed a weaker relationship (Fig. 6c), and age explained only a small proportion of variance in follow-up model performance (Fig. S18d). These findings suggest that hybrid FC is most informative when tau pathology is sufficiently expressed and spatially structured, but they also indicate that FC alone does not capture all sources of individual tau-PET heterogeneity. Other factors, including regional vulnerability, structural connectivity, local neurodegeneration, PET signal-to-noise, rs-fMRI reliability and specific individual features may also contribute to variability in explanatory performance.

Our findings highlight spatial scale as a critical factor in modelling the relationship between functional connectivity (FC) and tau pathology. While prior studies have used parcellations of 400 regions or fewer, we systematically evaluated FC and tau-PET associations across ten parcellation resolutions ranging from 100 to 1000 regions, enabling multiscale analysis of both modalities. Across analyses, higher-resolution parcellations consistently improved model performance, particularly in symptomatic individuals, where tau is more diffusely distributed and more variable (Fig. 2c; Fig. 6b). Across several analyses, the largest changes with increasing parcellation resolution occurred between the coarser atlases and approximately 400 cortical regions, with comparatively smaller changes thereafter. This pattern suggests diminishing returns in the group-level summary measures beyond an intermediate spatial resolution. However, it should not be interpreted as establishing 400 regions as a definitive cutoff or universally optimal parcellation. The present multiscale analyses were intended to assess robustness across spatial scales rather than to optimize a single atlas, and the preferred resolution will depend on the effective spatial resolution and signal-to-noise characteristics of the imaging modalities, the reliability of parcel-wise FC estimates, and the intended application. For the BioFINDER-2 data, an atlas of approximately 400 regions may offer a practical balance between spatial specificity and measurement stability, while the consistency of the principal findings from 400 to 1,000 regions indicates that they are not contingent on a single resolution. Finer parcellations may nevertheless be useful in future studies seeking to resolve more localized, potentially mesoscale, relationships between connectivity and tau pathology. For example, in patients with ADD, the median explained variance using hybrid FC increased from approximately 38% at 100 regions to nearly 70% at 1000 regions. This scale-dependent improvement was also evident when comparing hybrid FC to canonical tau-PET patterns (Fig. 3b) and A*β*-PET patterns (Fig. 5b), and was supported by regional null-model analyses showing that the observed FC–tau associations were not explained by lower-order graph properties or by spatial autocorrelation in tau-PET alone (Fig. 1c–d). Although the absolute relative advantage of hybrid over template FC decreased with increasing resolution due to performance saturation in subject FC models, particularly in CN and MCI (Fig. S2a), the difference remained positive and statistically strong across scales (Fig. 2d). These observations reflect a broader methodological point: coarser atlases introduce greater spatial averaging, obscuring local variation in both FC and tau-PET maps. To address this, we implemented a subject-space surface-based representation of the group-derived Schaefer atlas, applied in each subject’s native anatomical space. All PET and fMRI data were co-registered to the individual’s T1-weighted image and projected onto native cortical surfaces, avoiding warping to template spaces (e.g., MNI) and preserving fine-grained anatomical detail. This approach enabled accurate parcellation at high spatial resolutions and ensured consistency between modalities, resulting in richer input data for modelling. Importantly, if tau pathology indeed is constrained by brain-network architecture, then modelling this relationship at coarse spatial scales— where each region encompasses broad, heterogeneous cortical territories—may obscure spatially localized FC–tau correspondence related to individual variability. Notably, the use of hybrid FC, which combines subject-specific and template-level FC, helps mitigate the increased spatial noise and instability that can arise when estimating fine-grained FC matrices at the individual level, offering a stable yet personalized model that retains subject-specific features while enhancing signal reliability. Our results suggest that FC–tau correspondence is more readily detected at finer spatial scales within the present analytical framework, supporting the idea that spatial granularity and individualized modelling are important parameters for advancing connectome-based models of tau-PET heterogeneity.

Building on the observed correspondence between individual functional connectivity and tau pathology, this work lays the groundwork for the development of personalized, network-aware models of tau PET heterogeneity that may inform future prognostic modelling and individualized disease monitoring in Alzheimer’s disease. By showing that subject-specific FC robustly explains individual tau-PET patterns, and outperforms group-average and canonical models, particularly at finer spatial scales, we lay the groundwork for future evaluation of FC-based models for characterizing individual tau-PET heterogeneity and monitoring disease progression. Notably, the Default Mode and Frontoparietal networks consistently exhibited the highest explained variance across disease stages, emphasizing their centrality as networks in which individual FC profiles most consistently relate to tau-PET variability (Figs. S6 and S7). Importantly, these are also networks characterized by high inter-individual variability (Mueller et al., 2013; Finn et al., 2015), suggesting that they may be particularly informative settings in which subject-specific FC provides added explanatory value. These results suggest that individualized FC patterns capture not only global FC–tau correspondence, but also regional susceptibility linked to network architecture. Building on this, a critical next step will be integrating subject-specific FC with biological markers of regional vulnerability—such as excitatory neuron density (Fu et al., 2019; Leng et al., 2021), myelination gradients (Rubinski et al., 2022; Depp et al., 2023), microglial activity (Anand et al., 2022), and regionally patterned gene expression (Montal et al., 2022; Yu et al., 2024; Anand et al., 2024)—to better model why certain brain regions serve as preferential sites for tau accumulation (Xiao et al., 2025). This would allow hybrid models to not only characterize how tau-PET topography relates to an individual’s connectivity profile but also estimate where it is most likely to emerge or intensify. While the integration of such multimodal biological features lies beyond the scope of the present study, our individualized FC framework provides a scalable methodological platform for this endeavour. Together, these findings support a precision connectomics approach to AD, with potential relevance for future modelling and monitoring of tau-PET trajectories, pending further validation.

Key strengths of this study include the large BioFINDER-2 sample, longitudinal tau-PET follow-up, multimodal PET and fMRI characterization, multiscale modelling framework, and external replication in ADNI and A4. Several methodological safeguards further support the robustness of the main result. All in-sample model comparisons used corrected *R*^2^, so improvements should be interpreted as added explanatory information after penalizing model complexity rather than as raw increases in fit. The hybrid-FC advantage was preserved when FC matrices were density-thresholded (Fig. S3), regional FC– tau associations exceeded degree- and strength-preserving FC null models and Moran surrogate tau-PET maps (Fig. 1c– d; Fig. S1), and hybrid FC outperformed template FC under nested cross-validated ridge-regression prediction of held-out tau-PET parcels, with reproducible coefficient patterns across split-half analyses (Fig. 2e–f; Figs. S4 and S5). Together, these analyses indicate that the central model hierarchy was not driven by a single modelling decision, by spatial autocorrelation alone, or by in-sample model complexity alone.

However, several limitations should be acknowledged. First, FC estimated via resting-state fMRI is an indirect proxy for neuronal signalling and is sensitive to physiological noise, dynamic fluctuations, and scanner artifacts (Logothetis, 2008; Leonardi and Van De Ville, 2015; Lurie et al., 2020). It also fails to distinguish direct from polysynaptic pathways (Honey et al., 2009). These issues may reduce the mechanistic interpretability of FC as a marker of directed communication, especially given cautions against overinterpreting static FC estimates in isolation, as dynamic FC measures may conflate true neural variability with confounding sources (Liegeois et al., 2017; Lurie et al., 2020). A related interpretational consideration is that template FC was estimated from cognitively unimpaired Aβ-negative individuals as a relatively disease-insensitive normative reference, whereas subject FC and hybrid FC may contain both stable individual connectivity differences and disease- or age-related connectivity alterations. Thus, the advantage of subject-derived FC over template FC demonstrates added explanatory value of individualized connectivity information, but does not isolate whether this information reflects premorbid organization, disease-related FC changes, or both. Relatedly, we used a cognitively unimpaired A*β*-negative template FC as a normative reference rather than clinical-stage-specific template FC matrices. Therefore, part of the added value of subject-derived FC may reflect disease-or stage-related deviations from the healthy-control template. Future work could compare normative, disease-stage-specific, and individualized FC templates to determine how much explanatory information is attributable to group-level disease-stage effects versus person-specific connectivity variation. Nonetheless, the correspondence between our findings and results from animal models (Basheer et al., 2024), in which microscale neuronal dynamics can be more directly assessed, suggests a degree of cross-species consistency in the observed relationship between connectivity and tau pathology.

Second, while we demonstrate that subject-specific FC explains tau-PET topography, we did not explicitly distinguish between tau presence and load (Vogel et al., 2020; Xiao et al., 2025) in our models, nor did we incorporate orthogonal biological features that could modulate vulnerability, such as gene expression or blood flow. Third, limited spatial resolution of PET imaging compared to MRI may introduce noise, especially at higher granularity, though our results suggest that spatially structured tau-PET signal remains detectable. Fourth, although FC captures aspects of communication relevant to tau-PET topography, structural connectivity (SC) offers a more direct account of axonal pathways and may be especially informative in early disease stages (Liu et al., 2012; de Calignon et al., 2012; Vogel et al., 2020; Raj et al., 2021; Vogel et al., 2023; Xiao et al., 2025). Though subject-specific SC modelling remains technically difficult to robustly estimate from diffusion MRI at the clinical scale, our individualized FC framework lays the groundwork for future multimodal models that integrate both functional dynamics and structural constraints, potentially yielding more mechanistic insight into FC–tau relationships. Finally, while our focus was on cross-sectional FC, longitudinal changes in FC and their relationship to tau-PET change remain important future directions for establishing temporal dynamics and causal relationships.

A further limitation is that the analyses used a group-derived Schaefer parcellation mapped to each participant’s cortical surface rather than an individually estimated functional parcellation. Accordingly, the subject-specific FC matrices captured inter-individual differences in connectivity strength within a common set of parcels, but did not model individual variability in functional network boundaries or cortical topography. This common atlas framework facilitated correspondence across participants, imaging modalities, atlas resolutions, and template FC models, but may have reduced sensitivity to individualized functional organization. We also did not explicitly adjust the regional analyses for cortical parcel volume or surface area. Although all modalities were sampled using the same subject-space parcellation, parcel size may influence within-parcel averaging, partial-volume effects, signal-to-noise ratio, and the reliability of regional FC estimates. The consistency of the principal findings across parcellations ranging from 100 to 1,000 regions, together with thresholding sensitivity analyses and degree-and strength-preserving FC null models, argues against the results being driven solely by parcel size or lower-order network properties; however, these analyses do not constitute a direct parcel-volume correction. Future studies could test whether individually estimated functional parcellations and explicit modelling of regional surface area or volume further improve the relationship between individual FC and tau-PET patterns.

In conclusion, our findings show that subject-specific functional brain architecture provides additional explanatory information about the spatial distribution and longitudinal pattern of tau pathology in Alzheimer’s disease. By demonstrating that individual FC improves explanatory power beyond template and canonical tau models, replicates across independent cohorts, and is associated with follow-up tau-PET topography, our work supports a precision-connectomics framework for characterizing tau-PET heterogeneity. These findings are consistent with the network-spread hypothesis but do not establish a causal propagation mechanism. Future research should integrate individual Aβ patterns, structural connectivity, longitudinal FC, and regionally specific vulnerability factors to refine individualized models of tau-PET trajectories. Moreover, extending these models to capture the distinction between tau presence and load, and testing directional, activity-based mechanisms, will be important for refining individualized biomarker models and evaluating future approaches for monitoring tau-PET progression before widespread cortical involvement occurs.

## 4. Methods

### BioFINDER-2 primary cohort

We used data from the Swedish BioFINDER-2 study (NCT03174938). Information regarding recruitment, diagnostic criteria, and Aβ positivity assessment for the BioFINDER-2 cohort (https://biofinder.se/) has been described in detail previously (Leuzy et al., 2020; Palmqvist et al., 2020). All participants provided written informed consent in accordance with the Declaration of Helsinki. Ethical approval for the study was obtained from the Ethics Committee of Lund University, Lund, Sweden, with additional permissions for PET imaging granted by the Swedish Medicines and Products Agency and the local Radiation Safety Committee at Skåne University Hospital, Sweden.

Our study cohort included 805 participants aged 50+, consisting of 398 cognitively normal (CN), 77 with subjective cognitive decline (SCD), 156 with mild cognitive impairment (MCI), and 174 with AD dementia (ADD). Individuals with SCD, MCI, or ADD were all Aβ+ based on CSF testing whereas the CN individuals were split into two groups based on their CSF Aβ status; a pre-established cutoff of 0.08 on the CSF Aβ42/Aβ40 ratio was used to define Aβ positivity based on Gaussian mixture modeling (Palmqvist et al., 2020), and CSF Aβ42 and Aβ40 were measured using the Elecsys immunoassays of Roche Diagnostics (Hansson et al., 2018). All subjects had baseline resting-state fMRI and tau-PET, whereas A*β*-PET was available for 627 participants and was not available for patients with ADD. For the longitudinal part of the study, we used follow-up tau-PET scans from 422 participants. Table 1 shows the demographic and clinical characteristics of the sample.

### BioFINDER-2 MRI: imaging and image processing

Structural and functional MRI was acquired using a Siemens 3T MAGNETOM Prisma scanner (Siemens Healthineers, Erlangen, Germany) with a 64-channel head coil. T1-weighted images (Magnetization Prepared, Rapid Gradient Echo, MPRAGE) were acquired with the following parameters: in-plane resolution = 1 *×* 1 mm^2^, slice thickness = 1 mm, repetition time (TR) = 1900 ms, echo time (TE) = 2.54 ms, flip-angle = 9°. Resting-state fMRI data (eyes closed) were acquired using a 3D echo-planar imaging (EPI) sequence with an in-plane resolution of 3 *×* 3 mm and slice thickness 3.6 mm; echo time = 30 ms, and flip-angle = 63°. Scan time was 7.85 minutes, with a multiband repetition time of 1020 ms, resulting in 462 frames per scan before processing and censoring. Preprocessing has briefly been described in (Berron et al., 2021). The processing was performed using a modified CPAC (Craddock et al., 2013) pipeline, building mostly on FSL (Jenkinson et al., 2012), AFNI (Cox, 1996) and ANTS (Avants et al., 2014). Skull-stripping was done with in-house code using the T2 structural image as a primer and Brain Surface Extractor from BrainSuite (Shattuck and Leahy, 2002). T1-weighted images were also processed with FreeSurfer v6.0 to reconstruct cortical surfaces used for subsequent PET/fMRI surface projection and Schaefer atlas mapping.

The fMRI preprocessing included slice-timing and motion correction. Physiological noise was regressed out using CompCor (Behzadi et al., 2007), alongside the removal of linear and quadratic trends. Additionally, regression of Friston’s 24-parameter motion correction (Friston et al., 1996), white matter and CSF signals were performed. Susceptibility distortion was corrected by unwarping the functional data using a nonlinear diffeomorphic transformation (performed with ANTS) of the mean functional image to the high-resolution (1*×*1*×*1 mm^3^) T2 structural image (Wang et al., 2017); The T2-weighted structural image was used as the undistorted anatomical target for this established BioFINDER preprocessing workflow. This transformation was then applied to each volume in the fMRI time-series. This fieldmap-less correction strategy was used because dedicated high-quality field maps or reversed phase-encoding EPI acquisitions were not available in the BioFINDER-2 rs-fMRI protocol. Registration-based EPI distortion correction, in which distorted EPI data are nonlinearly aligned to an undistorted anatomical target such as a T1- or T2-weighted image, has been used previously as an alternative when fieldmap-based correction is not feasible (Wang et al., 2017; Albi et al., 2018; Esteban et al., 2019). This preprocessing approach follows prior analyses of BioFINDER rs-fMRI data, including work on tau-related functional connectivity and large-scale AD-related FC alterations (Berron et al., 2021; Rittmo et al., 2026). We therefore used the same pipeline consistently across all BioFINDER participants and analyses. Finally, a bandpass filter (0.01-0.1 Hz) was applied. For each scan, frames were censored based on DVARS for that frame lying outside above and below the third and first quartiles, respectively 1.5*×* IQR (Power et al., 2012). To avoid distortion from the outlier frames, they were interpolated before bandpass filtering and then removed from the final 4D image. Finally, any participant with an average frame displacement (FD) > 0.5 mm or maximum FD > 3 mm over the entire sequence were filtered out before analysis.

### BioFINDER-2 PET: imaging and image processing

All study participants underwent tau PET scans, and a subset of them as described above underwent Aβ-PET scans, all on digital GE Discovery MI scanners (General Electric Medical Systems). BioFINDER PET scanning procedures have been previously described in detail for tau-PET (Smith et al., 2020) and Aβ-PET (Palmqvist et al., 2014). Briefly, for tau PET, participants were injected with an average of 365 *±* 20 MBq [^18^F]RO948, and emission data was acquired in LIST mode between 70 and 90 minutes post-injection, adjusted for the tracer’s pharmacokinetics; for Aβ PET, an average of 185 MBq [^18^F]flutemetamol was used as tracer and emission data was acquired between 90 and 110 minutes post-injection. Lowdose CT scans were conducted before PET scans for attenuation correction. PET data was reconstructed using the VPFX-S algorithm (ordered subset expectation maximization with time-of-flight and point spread function corrections). The LIST mode data was binned into 4 *×* 5-minute time frames, and images were motion-corrected, summed, and co-registered to corresponding T1-weighted MRI images. FreeSurfer segmentation (v6; https://surfer.nmr.mgh.harvard.edu) was used to define reference regions for SUVR normalization, with uptake normalized to the mean uptake in inferior cerebellar grey matter for [^18^F]RO948 and to the whole cerebellum for [^18^F]flutemetamol. The resulting PET SUVR images were retained in each participant’s anatomical space and later projected to the cortical surface and parcellated using the shared subject-space Schaefer workflow described below.

### ADNI replication cohort

For external replication, we used data from the Alzheimer’s Disease Neuroimaging Initiative (ADNI; https://adni.loni.usc.edu/). Participants were included based on the availability of clinical data, baseline A*β*-PET using either [^18^F]florbetapir or [^18^F]florbetaben, tau-PET using [^18^F]flortaucipir, and 3T resting-state fMRI. All baseline imaging data were required to have been acquired within a 6-month time window. The ADNI replication sample included 361 participants, consisting of 149 CN A*β*-, 91 CN A*β*+, 78 MCI A*β*+, and 43 ADD A*β*+ individuals. Table 2 shows the demographic and clinical characteristics of the sample.

Diagnostic status was determined by ADNI. CN participants had a Mini Mental State Examination (MMSE) score ≥24, Clinical Dementia Rating (CDR) score of 0, and no depression. MCI participants had an MMSE score ≥ 24, CDR score of 0.5, objective memory impairment based on education-adjusted Wechsler Memory Scale II criteria, and preserved activities of daily living. Dementia participants had an MMSE score of 20–26, CDR score *>*0.5, and met National Institute of Neurological and Communicative Disorders and Stroke/Alzheimer’s Disease and Related Disorders Association criteria for probable AD. A*β* status was determined from global amyloid-PET SUVR using tracer-specific cutoffs of 1.11 for [^18^F]florbetapir and 1.08 for [^18^F]florbetaben, as previously established in ADNI (Landau et al., 2012; Royse et al., 2021). All ADNI study procedures were conducted in accordance with the Declaration of Helsinki. Ethical approval was obtained by the ADNI investigators, and all participants provided written informed consent.

### ADNI MRI: imaging and image processing

Structural and functional MRI data were acquired on 3T Siemens, GE, and Philips scanners using ADNI acquisition protocols. T1-weighted structural images were acquired using an MPRAGE sequence with 1 *×* 1 *×* 1 mm^3^ voxel size and TR = 2300 ms. Resting-state fMRI data were acquired using an echoplanar imaging sequence with 200 volumes per participant, TR = 3000 ms, TE = 30 ms, flip angle = 90^*◦*^, and voxel size = 3.4*×*3.4*×*3.4 mm^3^.

All images were screened for artifacts before preprocessing. T1-weighted structural MRI scans were bias-corrected using the CAT12 toolbox (https://neuro-jena.github.io/cat12-help/). FreeSurfer cortical surface reconstruction was performed for each participant to support subject-space surface projection and parcellation. Resting-state fMRI images were slice-time corrected and realigned to the first volume for motion correction, followed by co-registration to the corresponding T1-weighted image. Using rigid-transformation parameters, T1-derived grey matter, eroded white matter, and eroded cerebrospinal fluid (CSF) segments were transformed to EPI space. To denoise the EPI time series, nuisance covariates were regressed out, including mean signals from eroded white matter and eroded CSF, six motion parameters, and their temporal and dispersion derivatives. This was followed by detrending and band-pass filtering between 0.01 and 0.08 Hz in native space. To further reduce motion artifacts that may affect functional connectivity estimates (Power et al., 2014), motion scrubbing was performed by removing volumes exceeding a frame-wise displacement threshold of 0.5 mm, together with one preceding and two subsequent volumes. All included participants had at least 5 minutes of resting-state fMRI data remaining after scrubbing (Franzmeier et al., 2018). Spatial smoothing was not performed to avoid artificially increasing functional connectivity through signal spillover between adjacent brain regions. Susceptibility distortion correction was not applied in ADNI because field maps or reversed phase-encoding acquisitions were not consistently available across participants.

The preprocessed resting-state fMRI data were retained in native subject space and later projected onto each participant’s FreeSurfer-derived cortical surface using the shared subject-space parcellation workflow described below.

### ADNI PET: imaging and image processing

PET data were acquired after intravenous injection of [^18^F]flortaucipir, [^18^F]florbetapir, or [^18^F]florbetaben according to ADNI acquisition protocols. For [^18^F]flortaucipir tau-PET, six 5-minute time frames were acquired 75–105 minutes post-injection. For A*β*-PET, four 5-minute time frames were acquired 50–70 minutes post-injection for [^18^F]florbetapir and 90–110 minutes post-injection for [^18^F]florbetaben.

Dynamically acquired PET images were realigned and averaged to obtain a single PET image for each tracer and participant. The resulting PET images were rigidly registered to the corresponding T1-weighted MRI scan. PET SUVR images were generated using inferior cerebellar grey matter as the reference region for [^18^F]flortaucipir tau-PET and the whole cerebellum as the reference region for [^18^F]florbetapir and [^18^F]florbetaben A*β*-PET (Baker et al., 2017; Franzmeier et al., 2017). In ADNI, A*β*-PET was used to define A*β* status and clinical A*β* groups, whereas the replication analyses focused on tau-PET model comparisons. The resulting tau-PET SUVR images were retained in each participant’s subject-specific anatomical space and later projected to the cortical surface using the shared subject-space parcellation workflow described below.

### A4 replication cohort

For additional external replication, we used data from the Anti-Amyloid Treatment in Asymptomatic Alzheimer’s Disease study (A4). Participants were included based on the availability of clinical data, baseline A*β*-PET using [^18^F]florbetapir, tau-PET using [^18^F]flortaucipir, and 3T resting-state fMRI. The A4 replication sample included 336 cognitively normal participants, consisting of 36 CN A*β*- and 300 CN A*β*+ individuals. Table 3 shows the demographic and clinical characteristics of the primary A4 replication sample.

A*β* status was determined using the available global composite A*β*-PET SUVR measure and a cutoff of 1.15, as previously suggested by the A4 imaging core (Langford et al., 2020) and used in prior A4 analyses (Roemer-Cassiano et al., 2025). All A4 study procedures were conducted in accordance with the Declaration of Helsinki and were approved by the relevant institutional review boards. All participants provided written informed consent.

### A4 MRI: imaging and image processing

Structural and functional MRI data in A4 were acquired across multiple sites using 3T GE, Philips, and Siemens scanners. Acquisition parameters differed slightly across sites and scanner manufacturers but were largely harmonized. T1-weighted images were commonly acquired using an accelerated sagittal MPRAGE/IR-SPGR sequence with TR = 2300 ms, TE = 2.95 ms, TI = 900 ms, flip angle = 9^*◦*^, field of view = 270 mm, matrix = 256 *×* 256, and slice thickness = 1 mm. Resting-state fMRI data were acquired using an eyes-open axial EPI sequence lasting 10 minutes, with TR = 3000 ms, TE = 30 ms, flip angle = 80^*◦*^, field of view = 220 mm, matrix = 64 *×* 64, slice thickness = 3.3 mm, 48 slices, in-plane resolution = 3.3125 *×* 3.3125 mm^2^, and 200 volumes.

Structural and functional MRI data were preprocessed using fMRIPrep version 25.1.1 (Esteban et al., 2019). T1-weighted preprocessing included correction for intensity non-uniformity using N4BiasFieldCorrection, skull stripping using `antsBrainExtraction.sh`, tissue segmentation using FSL FAST, cortical surface reconstruction using FreeSurfer `recon-all` version 7.3.2, and brain-mask estimation using Mindboggle (Klein et al., 2017). Functional preprocessing included head-motion correction using FSL MCFLIRT with respect to the BOLD reference generated by fMRIPrep, BOLD-to-T1w co-registration using boundary-based registration with six degrees of freedom, and resampling of the preprocessed BOLD time series into native T1-weighted anatomical space using cubic B-spline interpolation.

When T2-weighted images were available, susceptibility-induced EPI distortions were corrected before fMRIPrep processing using the Synth pipeline (Montez et al., 2023). Briefly, Synth generated a synthetic undistorted image with EPI-like contrast from the T1- and T2-weighted anatomical images. This synthetic image served as an undistorted reference for estimating the deformation between the distorted EPI image and the synthetic target, and the resulting deformation field was applied to unwarp susceptibility-induced geometric distortions in the EPI data. The distortion-corrected EPI images were then used as the initial BOLD inputs for the fMRIPrep workflow. Detailed descriptions of the preprocessing pipelines and scripts are available at https://github.com/DeMONLab-BioFINDER/a4-learn_fmri-processing.

Following fMRIPrep, additional temporal processing was applied to the A4 resting-state fMRI data. Friston’s 24 motion parameters, white matter signal, and CSF signal were regressed out, followed by band-pass filtering between 0.01 and 0.1 Hz. Participants were excluded from the A4 analyses if average frame displacement exceeded 0.5 mm or maximum frame displacement exceeded 3 mm over the entire resting-state fMRI sequence. No additional frame-level censoring or scrubbing was applied to the A4 resting-state fMRI data. Spatial smoothing was not performed. The resulting preprocessed resting-state fMRI data were retained in native subject space and later projected onto each participant’s FreeSurfer-derived cortical surface using the shared subject-space parcellation workflow described below.

### A4 PET: imaging and image processing

Tau-PET scans were acquired over 30 minutes, starting 75 *±* 5 minutes after intravenous injection of 10 mCi (370 MBq) *±* 10% [^18^F]flortaucipir, followed by a 15–20 mL saline flush. Images were acquired with matrix size = 128 *×* 128, field of view = 350 mm, and slice thickness = 4 mm. Data were reconstructed into six 5-minute frames using an iterative reconstruction algorithm, with LOR-RAMLA/3D-RAMLA used for Philips scanners. A*β*-PET data were acquired using [^18^F]florbetapir at the baseline study visit. Based on the congruent A4/ADNI imaging protocols, florbetapir A*β*-PET consisted of four 5-minute frames acquired 50–70 minutes after injection. A*β* status was defined using the available global composite A*β*-PET SUVR measure and a cut-off of 1.15, as previously suggested by the A4 imaging core (Langford et al., 2020).

Tau-PET images were processed using SPM12. The six 5-minute dynamic tau-PET frames were first motion-corrected by realigning all frames to the mean PET image. The motion-corrected frames were then summed to generate a single 3D tau-PET image in native PET space. The mean PET image was coregistered to the participant’s FreeSurfer-derived T1-weighted image using normalized mutual information, and the summed tau-PET image was resliced to the participant’s original T1-weighted image grid. Tau-PET SUVR images were generated by normalizing uptake to the mean uptake in FreeSurfer-derived cerebellar grey matter. The resulting tau-PET SUVR images were retained in the same subject-specific anatomical space as the MRI data and were subsequently processed using the shared subject-space cortical projection and parcellation workflow described below.

### Subject-space cortical parcellation of PET and fMRI

For each participant in BioFINDER, ADNI, and A4, we generated subject-space cortical parcellations at 10 spatial resolutions, ranging from 100 to 1000 cortical regions in steps of 100, using the Schaefer atlas (Schaefer et al., 2018). The Schaefer atlas is a group-derived cortical parcellation based on functional connectivity patterns and spatial contiguity. The atlas was not individually re-estimated from each participant’s functional data; rather, the group-derived atlas labels were projected to each participant’s own FreeSurfer-derived cortical surface, yielding a common atlas represented in subject space.

First, subject-specific cortical surfaces were reconstructed from T1-weighted scans using FreeSurfer (FS; https://surfer.nmr.mgh.harvard.edu/) within each cohort-specific preprocessing workflow; this procedure and all following surface-based steps were performed separately for each hemisphere. Next, the cortical surface annotation file for the Schaefer atlas at each spatial resolution, defined in `fsaverage` space, was projected to each participant’s FreeSurfer-derived surface using `mri_surf2surf`. The FreeSurfer-processed T1-weighted image, represented at 1-mm isotropic resolution, served as the common anatomical reference space for each participant.

Preprocessed tau-PET, BioFINDER A*β*-PET, and rs-fMRI data were co-registered to the participant’s FreeSurfer T1-weighted image and subsequently projected onto the cortical surface by sampling volumetric data at the midpoint of the cortical ribbon using `mri_vol2surf` with the `-projfrac 0.5` option. For rs-fMRI, each preprocessed time frame was projected to the surface. Cortical maps were then parcellated using the subject-space Schaefer annotation files by averaging surface-vertex values within each parcel, using `read_- annotation` together with in-house functions.

For PET data, this yielded subject-space parcellated SUVR maps that were used directly in downstream analyses. For rs-fMRI data, this yielded regional time series that were subsequently used to construct an individual functional connectome for each atlas resolution. Thus, although rs-fMRI and PET were acquired at different native spatial resolutions, both modalities were analysed within spatially corresponding cortical parcels in each participant’s own FreeSurfer anatomical space, without warping the final parcellated data to a common volumetric template.

### Functional connectivity estimation

For each atlas with *P* cortical parcels and each subject, pair-wise Pearson correlations were computed between the rs-fMRI time series of all parcels, resulting in a *P × P* symmetric correlation matrix. Matrix diagonals, corresponding to autocorrelations, and negative correlations were set to zero. Remaining positive FC values were Fisher Z-transformed to extend the dynamic range of the connectivity values.

No ad hoc thresholding was applied in the primary analyses. After setting diagonal elements and negative correlations to zero, all positive FC values were retained to avoid introducing an arbitrary sparsification parameter and to preserve distributed connectivity information. For each atlas and cohort, template FC was generated by averaging the FC matrices of cognitively unimpaired A*β*-negative individuals within that cohort. This group-derived FC matrix served as template FC throughout the corresponding BioFINDER, ADNI, or A4 analyses. Subject FC refers to the FC matrix estimated separately from each participant’s own rs-fMRI data.

As a sensitivity analysis, we repeated the hybrid FC construction and whole-brain regression analyses using density-thresholded FC matrices. For both subject-specific FC and template FC, only the strongest 30% of positive connections were retained, a threshold also used in prior related work (Franzmeier et al., 2024). Hybrid FC was then reconstructed from these thresholded FC matrices using the same regional weighting procedure as in the primary analysis.

### Analytical overview

All analyses were performed across the ten Schaefer atlas resolutions described above. The primary analyses were conducted in BioFINDER and then repeated, where applicable, in ADNI and A4. First, regional FC–tau regression models were used to test whether subject-specific FC provided explanatory information beyond template FC and to obtain region- and subject-specific coefficients for constructing hybrid FC. These regional analyses were accompanied by FC rewiring null models and Moran surrogate tau-PET maps. Second, hybrid FC, template FC, and subject FC were compared in whole-brain OLS models of individual tau-PET topography, with cross-validated ridge-regression and split-half coefficient-stability analyses used as robustness analyses. Third, hybrid FC was compared with cohort-derived canonical PET patterns, first for tau-PET and then for A*β*-PET. Finally, baseline hybrid FC was used to explain follow-up tau-PET patterns and to test whether model performance was associated with baseline pathology, longitudinal tau-PET change, and age.

### Explained variance and corrected *R*^2^

Across all in-sample regression analyses, model performance was summarized using corrected *R*^2^ to account for differences in the number of predictors across models. Corrected *R*^2^ corresponds to adjusted explained variance after accounting for model complexity. For regional models, observations corresponded to cortical parcels excluding the seed parcel. For whole-brain models, observations corresponded to cortical parcels within each hemisphere. Cross-validated ridge-regression analyses were instead evaluated using out-of-sample *R*^2^ on held-out parcels, as described below.

### Null models for subject FC and tau-PET spatial autocorrelation

To test whether the increased explained variance from adding subject-specific FC simply reflected the addition of another FC-like regressor or the capture of lower-order graph properties, we generated null-model FC matrices that preserved selected network properties while randomizing higher-order topology. The first approach used a degree-preserving rewiring algorithm (Maslov and Sneppen, 2002). In this method, edges of the subject FC matrix were repeatedly rewired while maintaining each node’s degree, thereby preserving the distribution of the number of connections per region but eliminating the specific wiring pattern. The second approach used a simulated annealing algorithm for weighted networks (Milisav et al., 2025). This method additionally preserved nodal strength, defined as the total connection weight of each region, while randomizing the arrangement of weights across edges. Each null FC matrix was then entered into the same regional regression pipeline as the empirical subject FC, allowing direct comparison of explained variance across empirical and null FC models. For each subject and atlas resolution, one degree-preserving null FC matrix and one degree- and strength-preserving null FC matrix were generated.

To test whether regional FC–tau associations could be explained by spatial autocorrelation in tau-PET maps alone, we generated Moran spectral surrogate tau-PET maps for each subject and atlas resolution. First, a cortical spatial weights matrix was constructed from inter-regional geodesic distances between parcel centroids on the cortical surface. Moran spectral randomization was then used to generate surrogate tau-PET maps that preserved the spatial autocorrelation structure of each empirical tau-PET map while randomizing its spatial organization. The regional FC–tau regression analyses were repeated using these surrogate tau-PET maps as outcomes, with the same template FC and subject FC regressors as in the empirical analyses. Explained variance from empirical tau-PET maps was then compared with explained variance from the Moran surrogate maps. For each subject and atlas resolution, 10 Moran surrogate tau-PET maps were generated.

### Regional-level regression of individual tau-PET patterns on functional connectivity

To assess the added value of subject-specific functional connectivity (FC) in explaining individual tau-PET topographies, we applied linear regression models at the regional level (Fig. 1a). For each subject and each cortical parcellation atlas with *P* parcels, we modelled the subject’s tau-PET signal using the FC profile of each region as a regressor. Specifically, for a given parcel *k*, we regressed the subject’s tau-PET values across the remaining *P* − 1 regions onto the *k*-th column of the FC matrix, excluding the diagonal element. This procedure was repeated for both subject-specific FC and the group-level template FC, yielding one explained variance (*R*^2^) value per region, per subject, and per FC modality.

To determine whether subject-specific FC provides explanatory power beyond the template FC, we fit two-predictor linear models that included both the regional FC vector from the template and a second regressor derived from either subject-specific FC, a degree-preserving null FC, or a degree- and strength-preserving null FC. Importantly, the second regressor was orthogonalized with respect to the template FC regressor using a projection-removal procedure. This step removes the component of subject FC that lies along the template FC direction, thereby isolating the unique or residual component of subject FC. In other words, it identifies the part of the subject FC that is independent of the group-level template, ensuring that both regressors explain distinct sources of variance in the tau-PET signal.

Both the template FC and the orthogonalized subject FC regressors were 2-norm normalized prior to model fitting. This normalization was essential to ensure that the regression coefficients (*β*_1_ for template FC and *β*_2_ for subject FC) are comparable and interpretable, since otherwise the scale differences between regressors would bias coefficient estimates. The corrected *R*^2^ values from these models provided a matched distribution of variance explained across regions, subjects, and FC types, enabling fair comparison of explanatory performance across models. The regional coefficients from the empirical template-plus-subject-FC model were subsequently used to construct hybrid FC profiles.

### Network-specific analyses

To examine whether FC–tau associations differed across canonical functional networks, we repeated model summaries within the seven Schaefer network assignments: visual, somatomotor, dorsal attention, ventral attention, limbic, frontoparietal, and default mode networks. For the network-specific whole-brain analyses, models were fitted separately using the set of parcels belonging to each network, and explained variance was summarized for each subject, atlas resolution, network, and FC representation. We additionally quantified the difference in explained variance between hybrid FC and template FC within each network.

For the regional analyses, we summarized regional FC– tau model fits within each network by identifying, for each subject, atlas resolution, and network, the parcel with the highest explained variance from the regional template-plus-subject-FC model. We then compared the explained variance of this best-fitting regional model with the corresponding template-only model. These analyses were used to assess whether the contribution of subject-specific FC varied by functional network and spatial scale.

### Hybrid functional connectivity profiles

To enable joint modelling of subject-specific and group-level FC effects in a tractable way, we constructed hybrid functional connectivity (hybrid FC) profiles for use in the whole-brain modelling analysis described below. While the regional regression models allowed us to estimate the separate contributions of subject FC and template FC, including both as predictors in whole-brain models would double the number of regressors. The hybrid FC approach addresses this by combining both FC sources into a single FC profile per seed region, preserving the dimensionality of the whole-brain model while integrating both shared and individual-specific connectivity information.

For each subject, atlas resolution, and seed region, hybrid FC was constructed using the regional regression coefficients estimated from the empirical template-plus-subject-FC model described above. The coefficient for template FC reflected the association between the template FC profile and the subject’s tau-PET pattern, whereas the coefficient for subject FC reflected the association between the orthogonalized subject-FC component and the tau-PET pattern. These coefficients were then used as region- and subject-specific weights to combine the original, non-orthogonalized subject FC vector with the template FC vector.

Specifically, for each region and subject, we constructed a linear combination of the subject’s original FC vector and the template FC vector, using the corresponding regional coefficients (*β*_1_ for template FC and *β*_2_ for subject FC) as weights. Both input vectors were 2-norm normalized before weighting, and the resulting hybrid FC vector was normalized again after combination. Thus, although *β*_2_ was estimated from the subject-FC component after orthogonalization with respect to template FC, the hybrid FC profile itself was constructed from the original subject FC vector and the template FC vector.

This procedure yields a single integrated FC profile per region that reflects weighted contributions of normative and individual-specific connectivity information. The template FC captures group-level connectivity structure, while the subject FC captures each participant’s own connectivity profile. Hybrid FC should therefore be interpreted as a dimension-preserving modelling representation that combines template and subject-specific FC information, rather than as a directly observed biological connectome. Because hybrid FC replaces the original subject and template regressors with weighted combinations derived from regional fits, improvement at the whole-brain level is not mathematically guaranteed for every individual; occasional parity or small decrements are expected. The primary cross-sectional hybrid-FC analyses were interpreted as explanatory model-comparison analyses rather than fully independent predictive validation, because the regional weights used to construct hybrid FC were estimated from the same baseline PET modality subsequently evaluated in the whole-brain models. The purpose was therefore to compare whether an individualized FC representation better characterized subject-specific PET topography than template FC alone, rather than to estimate absolute predictive performance in an independent dataset.

For the A*β*-PET analyses, hybrid FC was reconstructed using *β* coefficients estimated specifically from A*β*-PET data rather than reusing coefficients derived from tau-PET analyses. Accordingly, the hybrid FC representations used in tau-PET and A*β*-PET analyses were estimated independently for each modality.

### Whole-brain OLS regression of individual tau-PET patterns on functional connectivity

To extend the analysis beyond regional models and evaluate the whole-cortex relationship between FC and tau-PET topography, we fit ordinary least-squares (OLS) linear models using the full FC profiles of each subject (Fig. 2a). For each FC representation, i.e., subject FC, template FC, and hybrid FC, all regional connectivity profiles within a hemisphere were used simultaneously to explain the subject’s tau-PET distribution in the same hemisphere. Self-connections were excluded, and each FC regressor vector was 2-norm normalized prior to inclusion. To reduce model dimensionality and keep model fitting within-hemisphere, models were fitted separately for the left and right hemispheres, and the resulting corrected *R*^2^ values were averaged across hemispheres for each subject, atlas resolution, and FC representation. This yielded one whole-brain explained-variance estimate per subject and FC model. These OLS analyses were used for in-sample model comparison; out-of-sample performance was assessed separately using the cross-validated ridge-regression analyses described below. This analysis provides a complementary, system-level perspective to the regional models and enables direct comparison across different FC representations.

### Cross-validated ridge-regression prediction of held-out tau-PET parcels

Because the whole-brain OLS models included many correlated FC predictors, we performed an additional regularized analysis to assess whether the main model-comparison findings generalized to held-out cortical parcels within each subject. For each subject, atlas resolution, hemisphere, and FC representation, we fit ridge-regression models in which tau-PET values across cortical parcels were modelled from the corresponding whole-brain FC profiles. The same FC representations were tested as in the OLS analyses: subject FC, template FC, and hybrid FC. Self-connections were excluded, and FC predictor vectors were 2-norm normalized before model fitting.

Model performance was evaluated using nested crossvalidation across parcels. In the outer loop, cortical parcels were divided into five held-out folds using network-stratified sampling, so that each fold preserved representation of canonical functional networks. Ridge models were trained on the remaining parcels and evaluated on the held-out parcels. In the inner loop, the ridge penalty parameter was selected using five-fold cross-validation within the training parcels. Candidate *λ* values were evaluated on a logarithmically spaced grid of 30 values from 10^−3^ to 10^3^, and the *λ* minimizing the inner-loop mean squared error was selected. The same outer partition seed was used across FC representations to ensure matched held-out folds for subject FC, template FC, and hybrid FC. Out-of-sample performance was quantified as cross-validated *R*^2^ for held-out tau-PET parcels.

Ridge-regression analyses were performed separately for the left and right hemispheres, and out-of-sample *R*^2^ values were averaged across hemispheres for each subject, atlas resolution, and FC representation. These analyses were used only as a robustness analysis of within-subject held-out-parcel prediction and were kept distinct from the primary in-sample OLS explained-variance analyses.

### Split-half stability of ridge-regression coefficients

To assess the reproducibility of the coefficient patterns estimated in the high-dimensional ridge-regression models, we performed a split-half stability analysis. For each subject, atlas resolution, hemisphere, and FC representation, cortical parcels were divided into two non-overlapping halves using network-stratified sampling, ensuring that both halves contained parcels from the same canonical functional networks. Ridge-regression models were then fitted independently in each half using the same FC representation, normalization procedure, and ridge-penalty selection framework described above.

Coefficient stability was quantified as the Pearson correlation coefficient between the two resulting ridge-regression coefficient vectors, excluding the intercept. This procedure was repeated across five network-stratified split-half repetitions using a fixed random seed, and stability values were averaged across repetitions and hemispheres for each subject, atlas resolution, and FC representation. The analysis was used to assess whether ridge-regression coefficient patterns were reproducible across independent parcel subsets, rather than to rank FC representations by coefficient stability.

### Canonical on- and off-target tracer binding PET patterns

Building on prior work (Vogel et al., 2020), we applied two-component Gaussian mixture modelling to PET SUVR values for each cortical parcel and atlas. Models were fit to the full cohort: 805 patients (CN A*β*-, CN A*β*+, SCD A*β*+, MCI A*β*+, ADD A*β*+) for tau-PET and 627 participants (CN A*β*-, CN A*β*+, SCD A*β*+, MCI A*β*+) for A*β*-PET. Each model yielded two overlapping Gaussian curves per modality, atlas, and parcel, representing estimated off-target and on-target binding distributions; see Fig. S8 for a schematic overview of the process.

While previous studies used these distributions to convert SUVRs to PET positivity probabilities (Vogel et al., 2020; Xiao et al., 2025), we instead aimed to derive canonical binding patterns that summarize cohort-level on- and off-target signals. These patterns allowed us to assess how well individual PET profiles align with canonical distributions versus more complex, connectivity-based models. The right-shifted Gaussian (higher mean) was designated as on-target binding. The means defined canonical off- and on-target values, generating two parcellated cortical maps per atlas (Figs. S9 and S10, left). Standard deviations captured uncertainty, revealing low spatial variability in off-target binding but substantial variability in on-target binding, positively correlated with the mean patterns (Figs. S9 and S10, right).

The resulting canonical PET patterns were used as cohort-derived predictor maps in the model-comparison analyses. For the ADNI replication analysis, ADNI-specific canonical tau-PET patterns were estimated from the ADNI tau-PET data using the same procedure.

### Model comparisons with canonical PET and individual A*β*-PET patterns

To compare FC-based models with canonical PET-pattern models, we used the same whole-brain regression framework described above. For each subject and atlas resolution, individual tau-PET values were modelled using canonical off-target tau-PET, canonical on-target tau-PET, canonical on- and off-target tau-PET, hybrid FC, or hybrid FC combined with canonical on- and off-target tau-PET. For FC-based models, predictors corresponded to regional FC profiles, as described above; for canonical PET models, predictors corresponded to cohort-derived regional PET-pattern maps sampled in the same atlas space. All predictor vectors were 2-norm normalized before model fitting, and self-connections were not applicable for canonical PET predictors because these were regional PET-pattern maps rather than FC profiles. Corrected *R*^2^ was used to compare models with different numbers of predictors.

In BioFINDER participants with available A*β*-PET, we additionally tested whether each individual’s A*β*-PET pattern explained tau-PET topography beyond canonical tau-PET patterns. For these analyses, individual parcellated A*β*-PET values were entered together with canonical tau-PET predictors and compared with hybrid FC. To assess modality specificity, the same canonical-pattern modelling framework was also applied with A*β*-PET as the outcome, comparing hybrid FC with canonical A*β*-PET patterns.

### External replication analyses

The ADNI and A4 replication analyses followed the same subject-space parcellation, FC estimation, template-FC construction, hybrid-FC construction, and whole-brain model-comparison workflow as the primary BioFINDER analyses. In both cohorts, we compared subject FC, template FC, and hybrid FC across atlas resolutions. In ADNI, we additionally compared hybrid FC with the ADNI-specific canonical tau-PET models described above. In A4, the replication analyses focused on the FC model comparisons because the cohort included only cognitively unimpaired participants.

### Non-parametric estimation of effect sizes, confidence intervals, and p-values

To assess differences in model performance across spatial scales and clinical groups, we used a non-parametric resampling-based procedure to estimate effect sizes (Cohen’s *d*), 95% confidence intervals (CIs), and permutation-based p-values. All comparisons were conducted on paired data, comparing model performance for the same set of individuals. Cohen’s *d* was computed as the mean difference in explained variance between models, divided by the standard deviation of the within-subject differences. Confidence intervals for the effect size were obtained via bootstrapping with 10,000 iterations. On each iteration, a bootstrap sample was drawn with replacement from the vector of within-subject differences, and Cohen’s *d* was recomputed. The 95% CI was defined as the 2.5th and 97.5th percentiles of the resulting distribution of bootstrap estimates. Permutation-based p-values were computed using a sign-flipping approach: for each of 10,000 iterations, the signs of the within-subject differences were randomly flipped to simulate the null distribution. The two-tailed p-value was then defined as the proportion of permutations where the absolute mean difference was greater than or equal to the observed value. This non-parametric approach avoids assumptions of normality, controls for subject-level variability, and yields robust estimates of effect size and statistical significance across spatial scales. Because effect sizes and their confidence intervals are estimated via resampling, the approach naturally accommodates differences in group sample sizes, allowing for consistent interpretation of Cohen’s *d* across clinical groups.

For between-group comparisons, where performance was compared across independent subject groups (e.g., CN Aβ-vs. CN Aβ+), we applied the same resampling-based framework adapted for unpaired data. Cohen’s *d* was computed as the difference in group means divided by the pooled standard deviation. Bootstrapped confidence intervals for the effect size were generated using 10,000 iterations, resampling with replacement separately within each group. Permutation-based p-values were obtained by permuting group labels and recomputing the mean difference on each iteration to construct a null distribution. These procedures allow valid statistical inference without assumptions of equal variance or normality, and remain robust even in the presence of unequal group sizes.

### Longitudinal tau-PET analyses

For participants with available follow-up tau-PET, we tested whether FC representations derived from baseline data explained tau-PET topography at follow-up. Hybrid FC was constructed from baseline data only, using baseline rs-fMRI and the regional regression coefficients estimated from baseline tau-PET. Follow-up tau-PET data were not used in the construction of hybrid FC. Baseline template FC and baseline hybrid FC were then entered into the same whole-brain OLS framework described above to explain follow-up tau-PET patterns, with corrected *R*^2^ averaged across hemispheres for each subject, atlas resolution, and FC representation.

To compare baseline and follow-up model performance within the same individuals, we repeated the whole-brain analyses for baseline tau-PET using the subset of participants who also had follow-up tau-PET. We then compared explained variance for baseline and follow-up tau-PET and quantified the correlation between baseline and follow-up explained variance across subjects.

### Association between baseline pathology and explained variance of baseline and follow-up tau-PET

To test whether baseline pathology influenced the explanatory performance of hybrid FC, we fit linear models relating baseline A*β*-PET, early tau-PET (Braak I–IV), longitudinal tau-PET change, or age to the explained variance of tau-PET at baseline and follow-up (Fig. 6c–e; Fig. S18d), using hybrid FC derived from each individual’s baseline resting-state fMRI. The analysis aimed to assess how explained variance in baseline and follow-up tau-PET relates to baseline pathological measures, longitudinal tau-PET change, and age. All predictors and outcomes were z-scored to obtain standardized effect sizes, and sex and, for pathology models, age were included as covariates. To evaluate robustness, the analysis was repeated across spatial scales.

For the analyses shown in Fig. 6c–e, CN A*β*-individuals were excluded to avoid leverage from consistently low biomarker levels. For A*β*-PET analyses, patients with ADD were excluded because A*β*-PET was not available in this group. Longitudinal tau-PET change was quantified from Braak I–IV tau-PET values between baseline and follow-up.

We additionally performed mediation analyses to test whether baseline A*β*-PET mediated the association between baseline early tau-PET burden and hybrid-FC explained variance at baseline or follow-up (Fig. S19). These models used baseline A*β*-PET as the mediator, baseline Braak I–IV tau-PET as the independent variable, and hybrid-FC explained variance as the outcome, with age and sex included as covariates.

## Data availability

The primary BioFINDER-2 data are not publicly available for download because they contain sensitive human participant data and are governed by ethical and legal restrictions. Anonymized BioFINDER-2 data may be shared upon reasonable request from a qualified academic investigator, provided that the proposed data transfer is compatible with European Union General Data Protection Regulation requirements, approvals from the Swedish Ethical Review Authority and Region Skåne, and an appropriate data/material transfer agreement.

ADNI data used in this study are available through the Alzheimer’s Disease Neuroimaging Initiative database (https://adni.loni.usc.edu/) upon registration and compliance with the ADNI data-use agreement. A4 data used in this study are available through the LONI Image and Data Archive / A4 data access procedures, subject to registration, approval, and applicable data-use agreements. Processed derivatives generated from controlled-access imaging data are subject to the same cohort-specific access restrictions.

## Code availability

MATLAB code supporting the core methodological analyses is available at https://github.com/aitchbi/taufc. The repository provides documented implementations and example workflows for the main analytical components, including subject-space Schaefer parcellation, surface-based projection and parcellation of rs-fMRI and PET data, FC–PET regression models, hybrid FC construction, canonical PET-pattern modelling, null-model analyses, and cross-validated ridge-regression analyses. The repository does not include controlled-access BioFINDER-2, ADNI, or A4 imaging data. Additional code details required to reproduce specific manuscript panels can be made available from the corresponding authors upon reasonable request, subject to cohort-specific data-use restrictions.

## Acknowledgements

We would like to express our gratitude to the research volunteers who participated in the studies from which these data were obtained and their supportive families. Work at the authors’ research center was supported by the National Institute of Aging (R01AG083740), European Research Council (ADG-101096455, StG-101221737), Alzheimer’s Association (ZEN24-1069572, SG-23-1061717, ALZSI-26-1523522), GHR Foundation, Swedish Research Council (2022-00775, 2021-02219, 2024-03642), ERA PerMed (ERAPERMED2021-184), Knut and Alice Wallenberg foundation (2022-0231), the SciLifeLab & Wallenberg Data Driven Life Science Program (grant: KAW 2020.0239), Strategic Research Area MultiPark (Multidisciplinary Research in Parkinson’s disease) at Lund University, Swedish Alzheimer Foundation (AF-980907; AF-981041, AF-1011799), Swedish Brain Foundation (FO2021-0293, FO2025-0055), Parkinson foundation of Sweden (1412/22), Cure Alzheimer’s fund, Rönström Family Foundation, Konung Gustaf V:s och Drottning Victorias Frimurarestiftelse, Skåne University Hospital Foundation (2020-O000028), Regionalt Forskningsstöd (2022-1259, 2025-2026-2024-2428), WASP and DDLS Joint call for research projects (WASP/DDLS22-066), GRIP Global Research and Imaging Platform, Michael J Fox Foundation (MJFF-025507), Swedish federal government under the ALF agreement (2022-Projekt0080, 2022-Projekt0107), Grant 2021-613 of the Strategic Focus Area “Personalized Health and Related Technologies (PHRT)” of the ETH Domain (Swiss Federal Institutes of Technology), and Swiss National Science Foundation under Grant 205321-163376. The funding sources had no role in the design and conduct of the study; in the collection, analysis, interpretation of the data; or in the preparation, review, or approval of the manuscript. The precursor of [^18^F]flutemetamol was sponsored by GE Healthcare. The precursor of [^18^F]RO948 was provided by Roche. L.E.C. is supported by the MSCA Postdoctoral fellowship (101108819), Alzheimer Association Research Fellowship (23AARF-1029663) and Alzheimer Nederland (WE.25-2024-01) grants. H.H.B. received funding from the European Union’s Horizon Europe Research and Innovation Program under Marie Sklodowska-Curie action grant agreement number 101153323 and travel grants from the Strategic Research Area MultiPark (Multidisciplinary Research in Parkinson’s Disease) at Lund University.

---

**Fig. S1.**
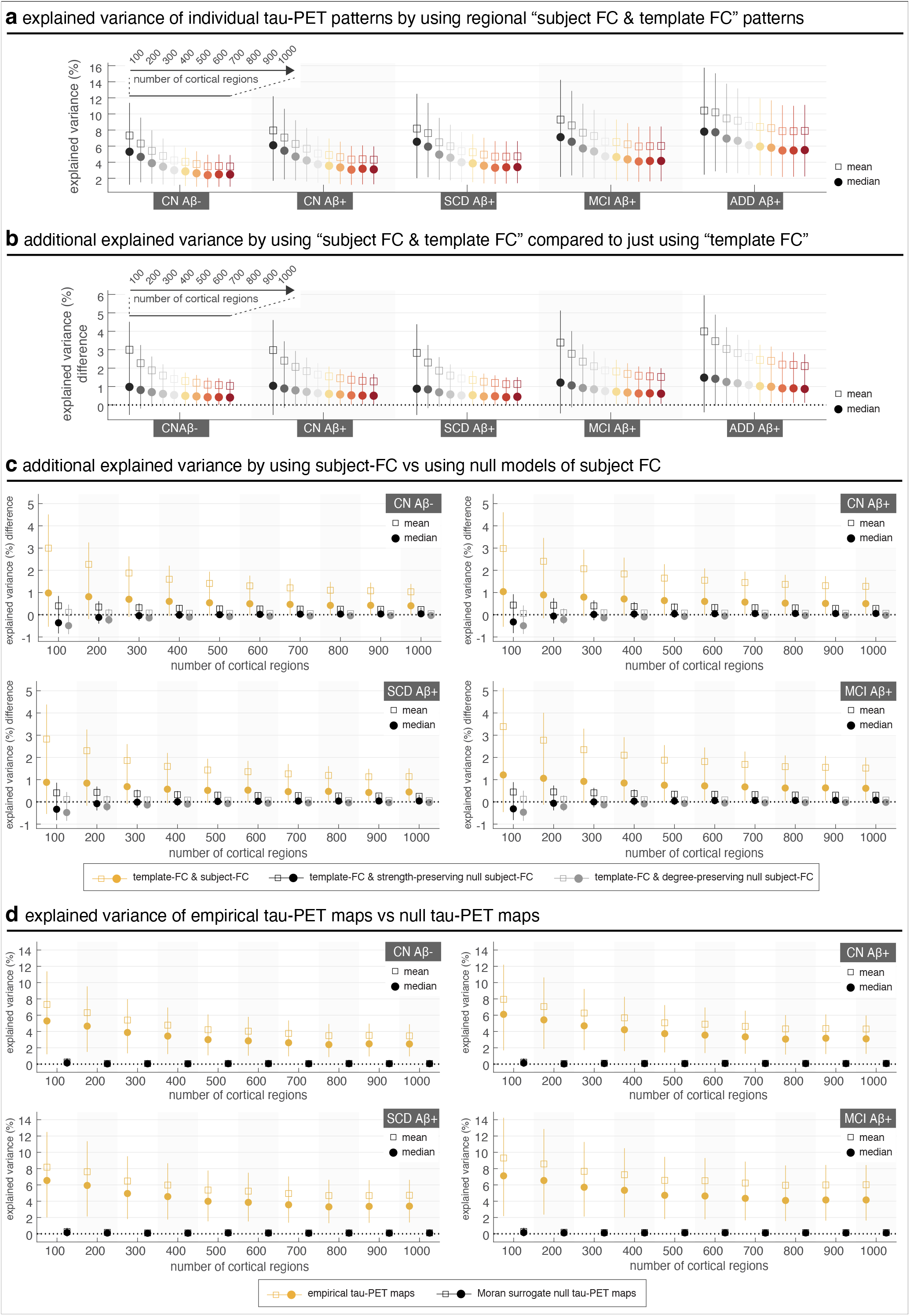
Explained variance of individual tau-PET by using both regional subject FC and template FC, across spatial scales and across groups. (a)–(b) Overall explained variance (a) and the additional explained variance compared to using just template FC (b) is greater at later stages and decreases as a function of spatial granularity of the design (i.e., cortical area covered by regions decreased). It is, however, important to note that as the number of regions in the atlases increases, FC of each region entails information of a smaller fraction of the entire FC; thus, the explained variance by regional FC decreases. (c) Additional explained variance from including subject FC, compared to two null models where subject FC was rewired to preserve either nodal degree (Maslov and Sneppen, 2002) or both nodal degree and strength (Milisav et al., 2025). Empirical subject FC produced consistently greater explained variance than the null subject FC models, indicating that the added explanatory value of subject FC depended on its specific wiring pattern rather than preserved degree or strength alone. (d) Explained variance obtained when regional FC–tau regression models were fitted to empirical tau-PET maps versus Moran surrogate tau-PET maps. Moran surrogate maps preserved the spatial autocorrelation structure of each empirical tau-PET map while randomizing its spatial organization. Explained variance values for surrogate maps were close to zero across clinical groups and atlas resolutions, indicating that the observed regional FC–tau associations were not attributable to spatial autocorrelation alone. In all panels, whiskers represent the 25th to 75th percentiles; markers indicate the mean and median, computed across regions and subjects in each group.

**Fig. S2.**
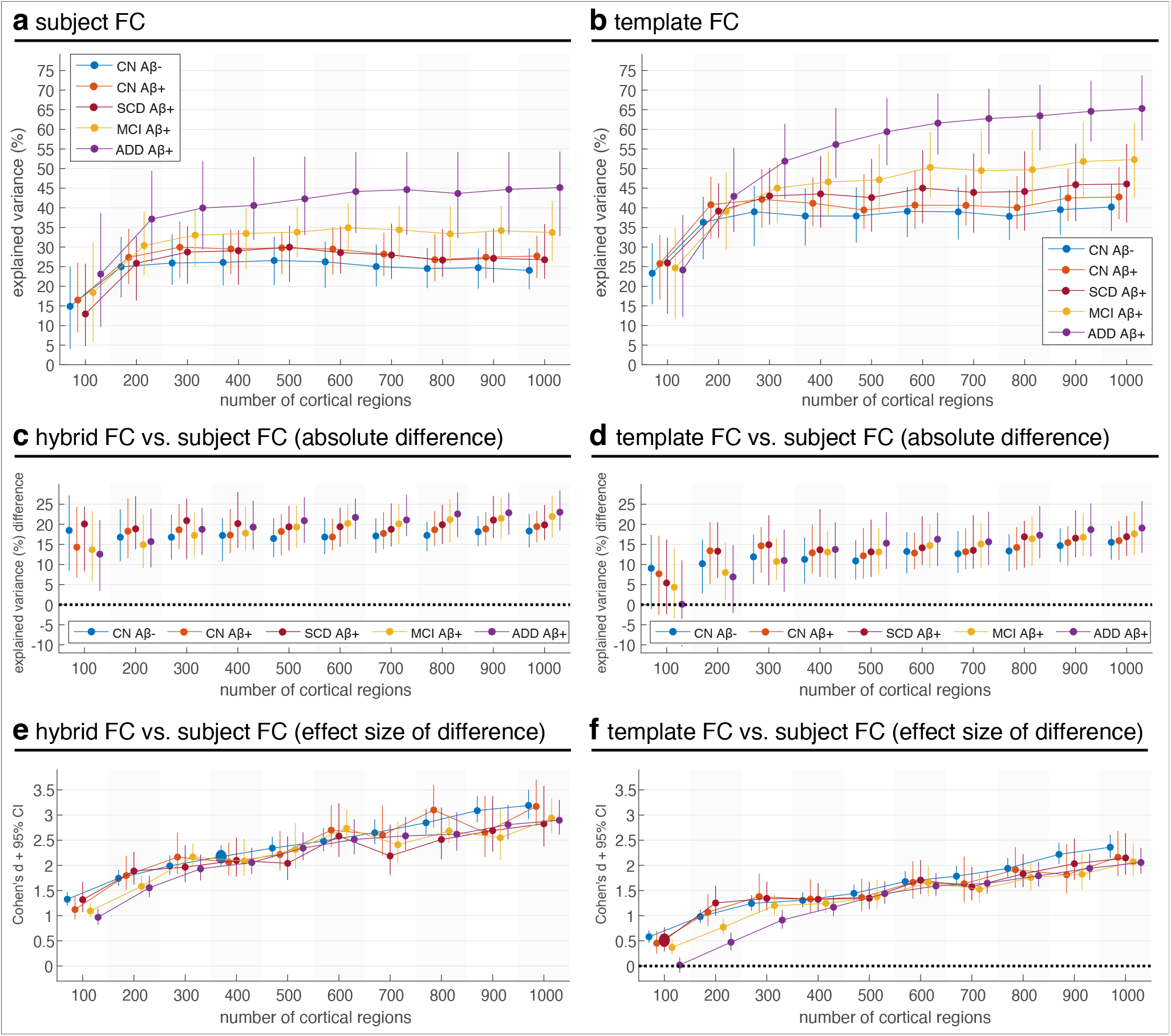
Hybrid FC consistently outperforms subject FC and template FC in explaining individual tau-PET patterns across spatial scales and clinical stages. (a) Explained variance (*R*^2^) from subject FC increases modestly with spatial granularity, peaking in ADD patients. (b) Template FC explains more variance than subject FC across groups, especially at finer scales and in ADD, echoing Fig. 2c (top). (c) Hybrid FC outperforms subject FC, with the largest gains in MCI and ADD, particularly at finer parcellations. (d) These gains exceed those from template FC vs. subject FC across groups and scales. (e–f) Corresponding effect sizes (Cohen’s *d*) for panels (c) and (d), showing that hybrid FC’s advantage over subject FC grows with granularity and surpasses that of template FC. These results support Fig. 2, showing that hybrid FC (a combination of subject- and template FC) outperforms both components alone. While subject FC explains little variance early in disease, hybrid FC significantly improves performance, especially in MCI and ADD, and remains robust across scales. The superior performance of template FC at high resolutions (panel b vs. a) explains hybrid FC’s increasing advantage with granularity (panel c). Overall, hybrid FC better captures tau-PET variability across disease stages and spatial resolutions. In panels a–d, whiskers show 25th–75th percentiles; markers indicate median. In panels e–f, whiskers denote 95% CI around Cohen’s *d*; markers indicate point estimates. Template FC refers to the group-derived functional connectivity matrix estimated from amyloid-negative cognitively healthy controls; subject FC was estimated separately from each participant’s rs-fMRI data. Hybrid FC combines template and subject-specific FC using region- and subject-specific weights.

**Fig. S3.**
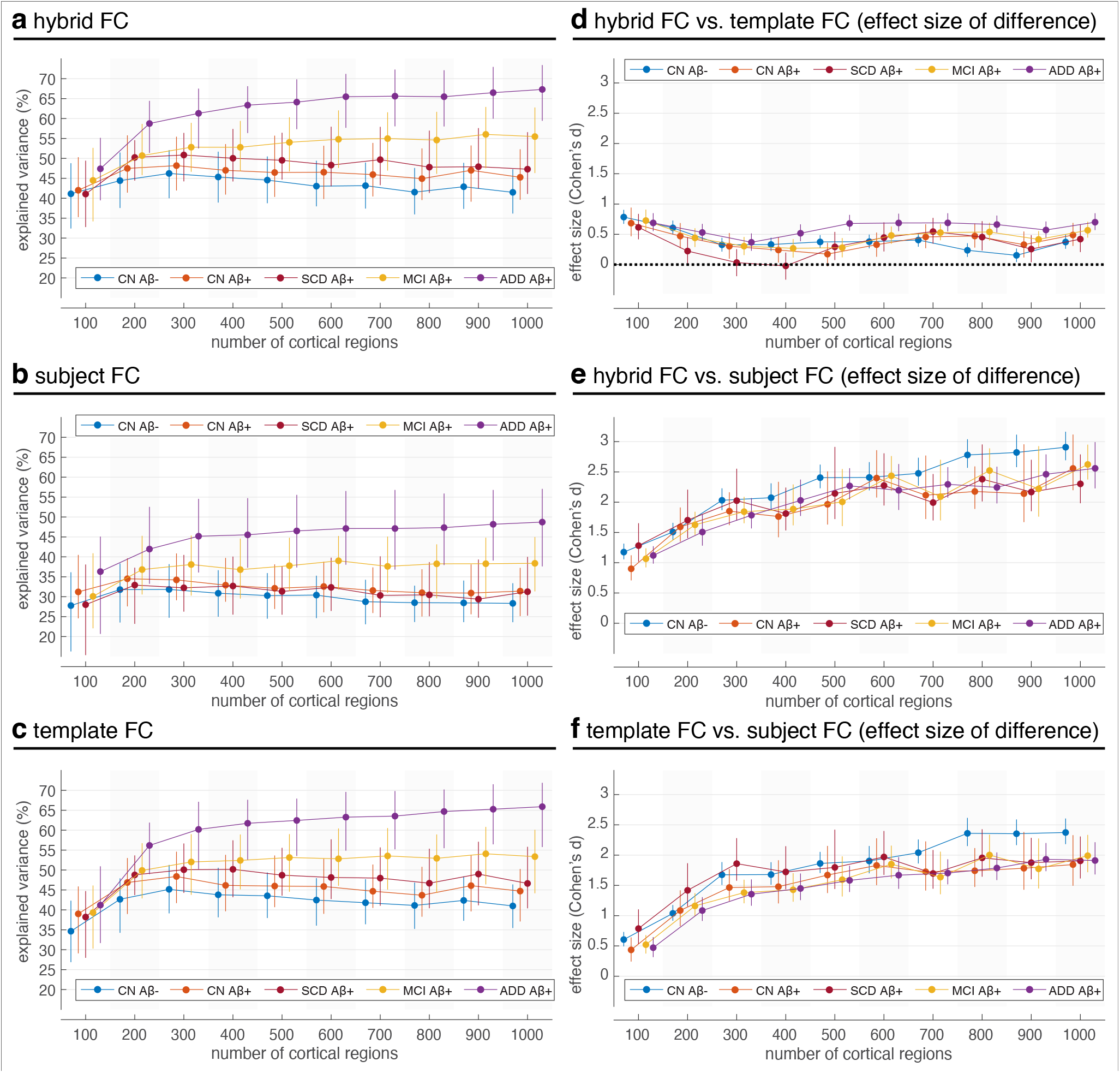
Sensitivity analysis using density-thresholded FC matrices. To assess whether the main findings depended on fully connected, unthresholded FC matrices, we repeated the hybrid FC construction and whole-brain regression analyses after sparsifying both subject-specific and template FC matrices to retain the strongest 30% of positive connections. Hybrid FC was reconstructed from the thresholded subject-specific and template FC matrices using the same regional weighting procedure as in the primary analysis. (a–c) Explained variance of individual tau-PET patterns using thresholded hybrid FC, subject FC, and template FC, respectively. (d) Standardized difference in explained variance between thresholded hybrid FC and thresholded template FC models. (e) Standardized difference in explained variance between thresholded hybrid FC and thresholded subject FC models. (f) Standardized difference in explained variance between thresholded template FC and thresholded subject FC models. Positive Cohen’s *d* values indicate better performance of the first model named in each panel title. Points and error bars show group-level mean *±* 95% confidence interval across subjects for each clinical group and atlas resolution. Template FC refers to the group-derived functional connectivity matrix estimated from amyloid-negative cognitively healthy controls; subject FC was estimated separately from each participant’s rs-fMRI data.

**Fig. S4.**
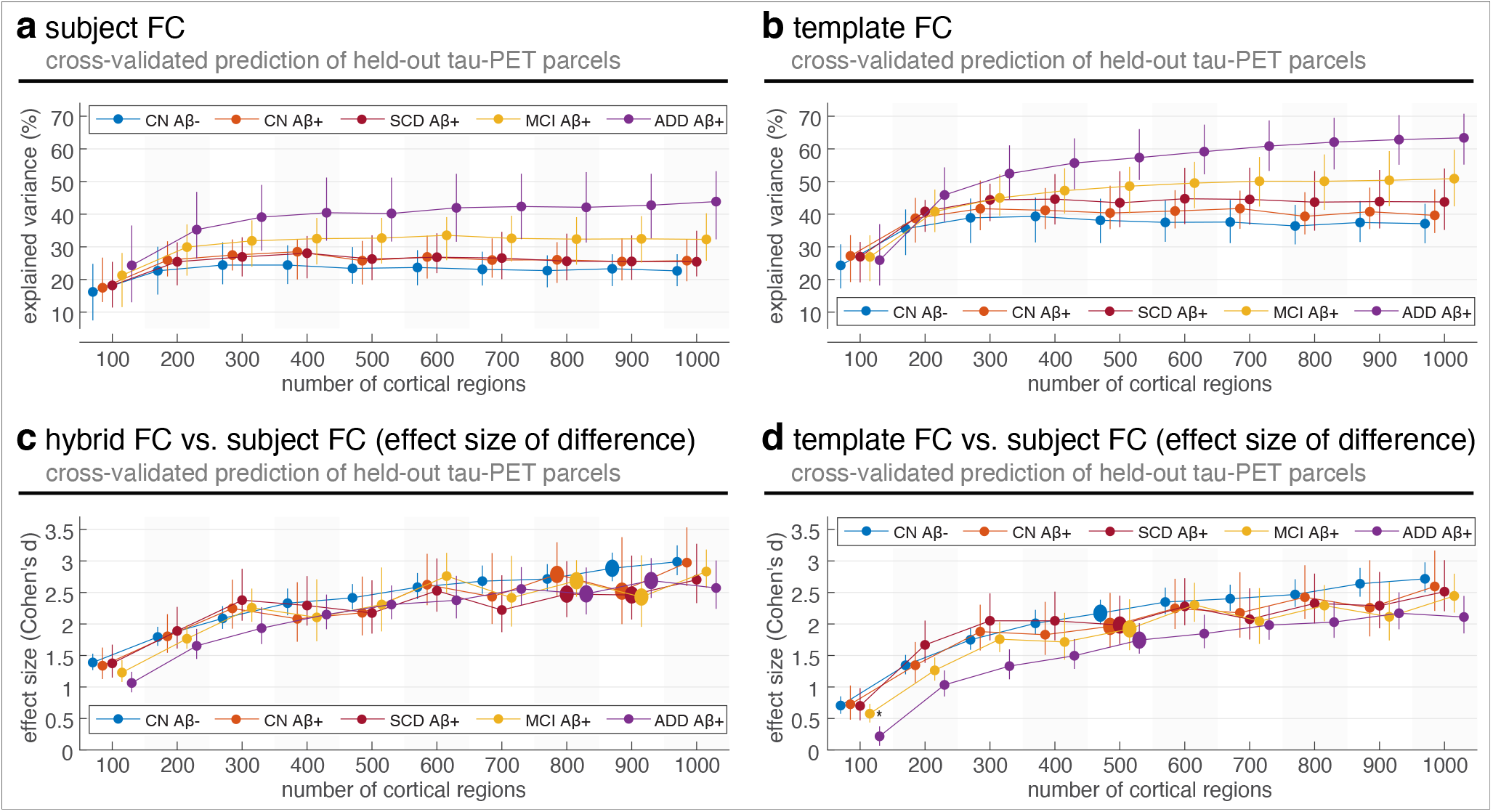
Cross-validated ridge-regression prediction of held-out tau-PET parcels across FC representations, clinical groups, and spatial scales. (a) Out-of-sample explained variance (*R*^2^) from subject FC increased modestly with spatial granularity, with highest values generally observed in symptomatic groups. (b) Out-of-sample explained variance from template FC showed stronger performance than subject FC across groups and atlas resolutions. (c) Hybrid FC outperformed subject FC in cross-validated prediction, with standardized differences quantified using Cohen’s *d*. (d) Template FC also outperformed subject FC, with standardized differences quantified using Cohen’s *d*. Together with Fig. 2e–f, these results show that the superior performance of hybrid FC was not limited to in-sample OLS explained variance but was also observed under regularized out-of-sample prediction. In all panels, ridge models were trained and evaluated using nested, network-stratified cross-validation across held-out tau-PET parcels. Points and error bars show group-level mean *±* 95% confidence interval across subjects for each clinical group and atlas resolution. Template FC refers to the group-derived functional connectivity matrix estimated from amyloid-negative cognitively healthy controls; subject FC was estimated separately from each participant’s rs-fMRI data.

**Fig. S5.**
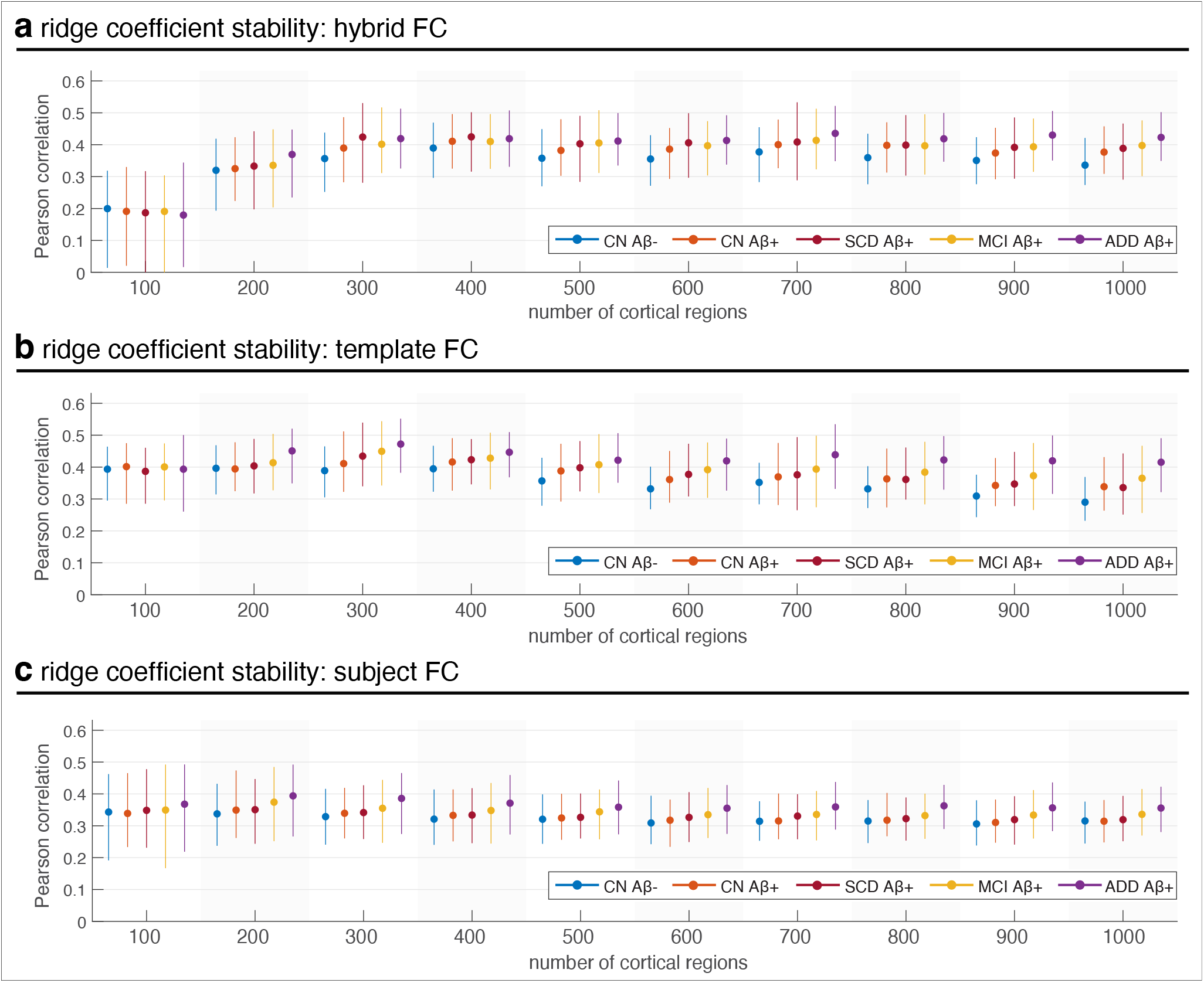
Split-half stability of ridge-regression coefficients in whole-brain FC models. *β*-coefficient stability was assessed for whole-brain ridge-regression models fitted to individual tau-PET patterns using (a) hybrid FC, (b) template FC, and (c) subject FC. For each subject, atlas resolution, hemisphere, and FC representation, cortical parcels were divided into two network-stratified halves, ridge models were fitted independently in each half, and stability was quantified as the Pearson correlation coefficient between the resulting *β* coefficient vectors, excluding the intercept. Correlations were averaged across split-half repetitions and hemispheres. Points and error bars show group-level mean *±* standard deviation across subjects for each clinical group and atlas resolution. Consistently positive correlations indicate reproducible coefficient patterns under regularized estimation across FC representations. Template FC refers to the group-derived functional connectivity matrix estimated from amyloid-negative cognitively healthy controls; subject FC was estimated separately from each participant’s rs-fMRI data.

**Fig. S6.**
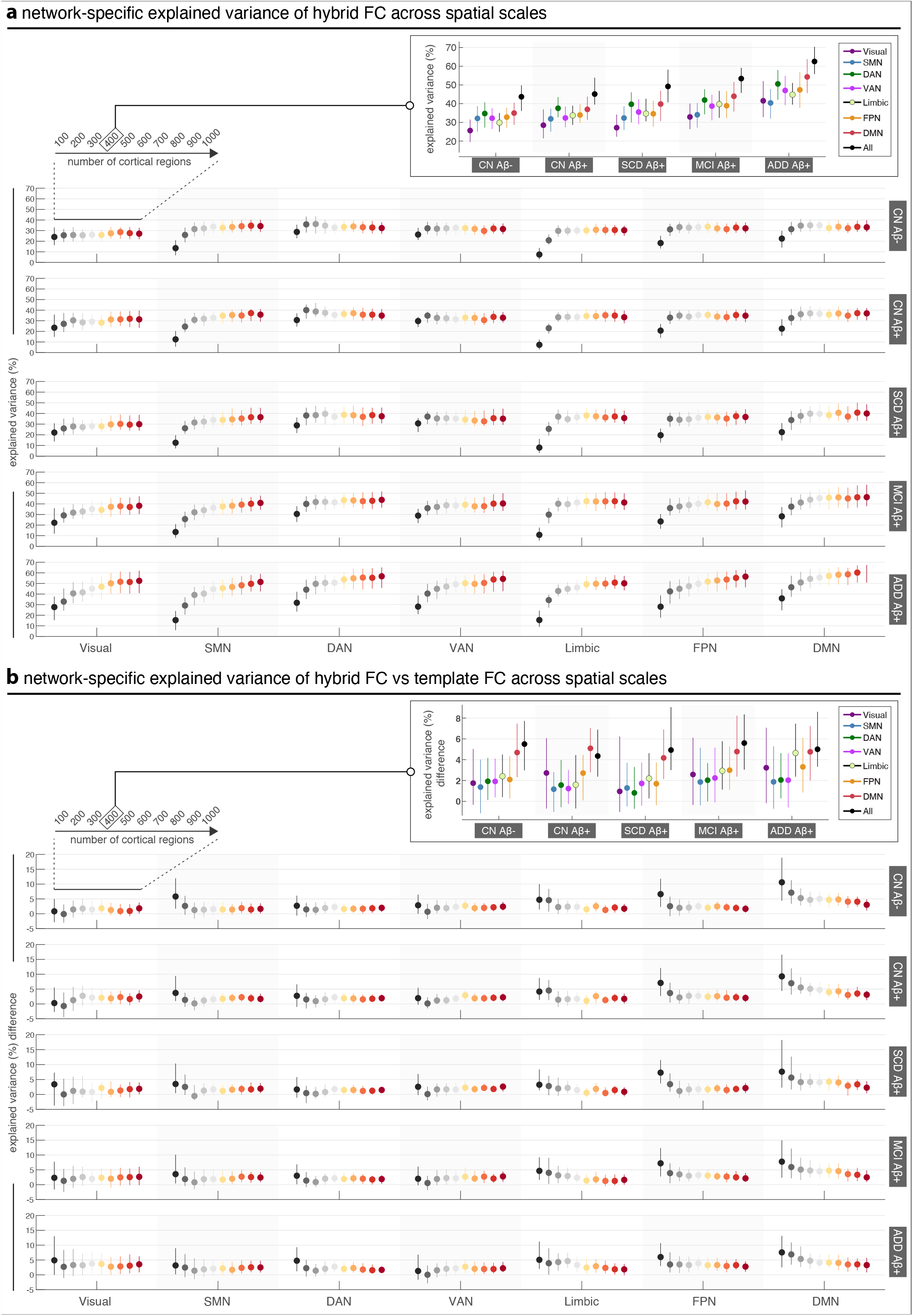
Network-specific model fits across spatial scales using regions spanning entire networks. (a) Explained variance of individual tau-PET patterns using hybrid FC, fit separately within each canonical network, across spatial scales and clinical groups. For each network, models were fit using the regional FC profiles of all regions within the network. The DMN and FPN consistently showed the highest explained variance, particularly in MCI A*β*+ and ADD groups, whereas the limbic network was stronger in earlier A*β*+ groups but plateaued or declined at finer parcellations. Visual and limbic networks also showed sharper declines in performance at higher resolutions, in contrast to the DMN and FPN, which remained stable. (b) Additional explained variance of hybrid FC relative to template FC, computed separately for each network, spatial scale, and group. Gains in the DMN and FPN were robust across scales, while benefits in the limbic and visual networks peaked at intermediate resolutions (400–600 regions). These findings emphasize that the contribution of individual FC varies by both network and disease stage, with the DMN and FPN most consistently capturing tau-related variability. Whiskers represent the 25th to 75th percentiles across subjects; markers indicate the median. Template FC refers to the group-derived functional connectivity matrix estimated from amyloid-negative cognitively healthy controls; subject FC was estimated separately from each participant’s rs-fMRI data.

**Fig. S7.**
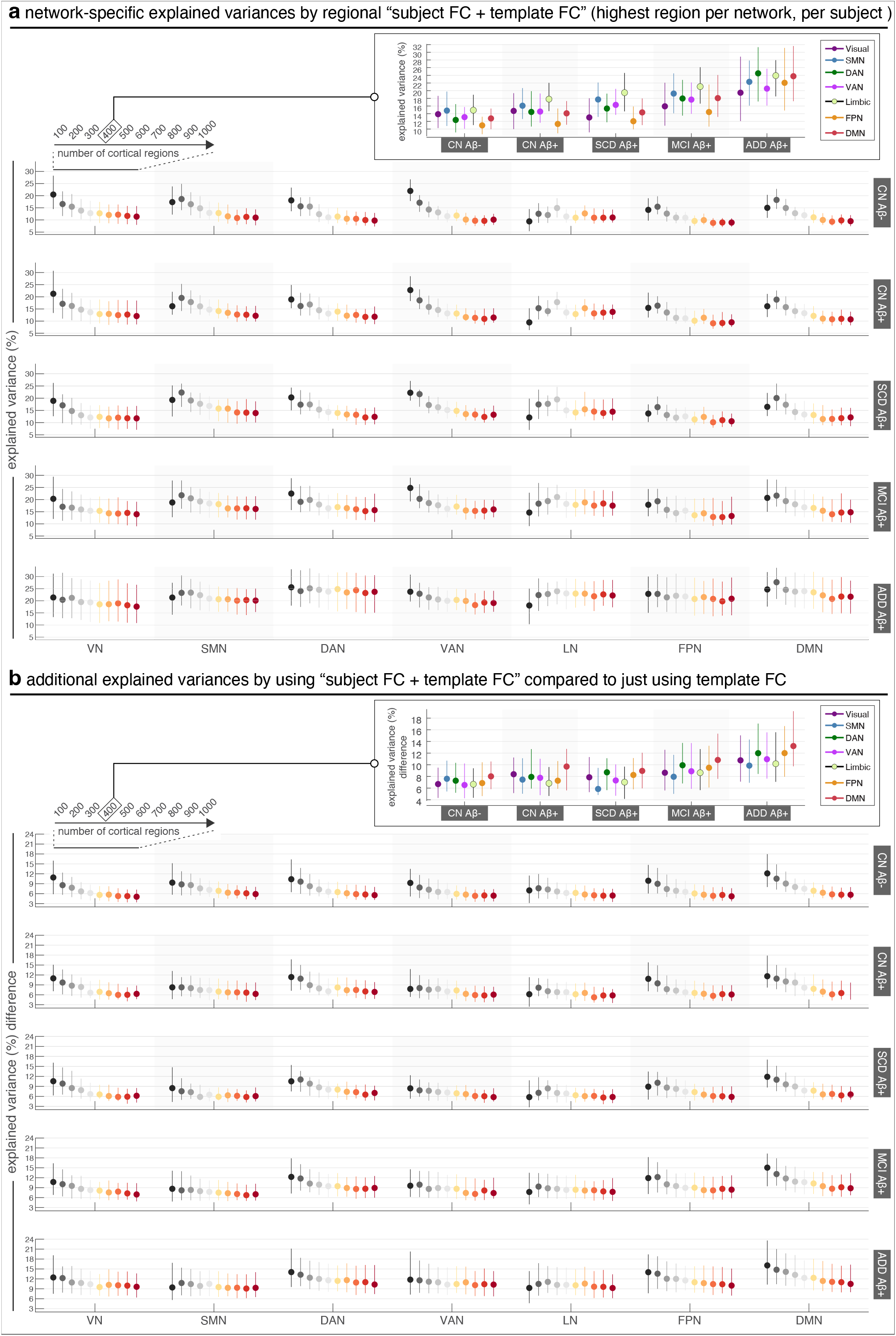
Network-specific model fits across spatial scales using regional FC. (a) Explained variance of individual tau-PET patterns using regional “subject FC + template FC”. For each subject and network, the region with the highest explained variance was selected. Performance varied across networks and disease stages, with the Default Mode (DMN) and Frontoparietal (FPN) networks generally yielding the strongest fits, especially in symptomatic A*β*+ groups. The limbic network also showed notable contributions in earlier A*β*+ groups, though its performance was more sensitive to spatial resolution, peaking at intermediate atlas sizes (400 regions). (b) Additional explained variance gained by including both subject and template FC compared to template FC alone. Gains were most consistent in the DMN and FPN, while other networks (e.g., limbic and visual) showed scale- and group-dependent improvements. Together, these results highlight that individual FC contributes explanatory power beyond group-level FC, and that the most informative regions for capturing tau variability often lie within networks previously implicated in AD progression. Whiskers represent the 25th to 75th percentiles across subjects; markers indicate the median. “Subject FC + template FC” denotes a joint regression model in which subject-specific FC and template FC were entered simultaneously as separate predictors. This differs from hybrid FC, which was constructed as a weighted combination of the two FC profiles. Template FC refers to the group-derived functional connectivity matrix estimated from amyloid-negative cognitively healthy controls; subject FC was estimated separately from each participant’s rs-fMRI data.

**Fig. S8.**
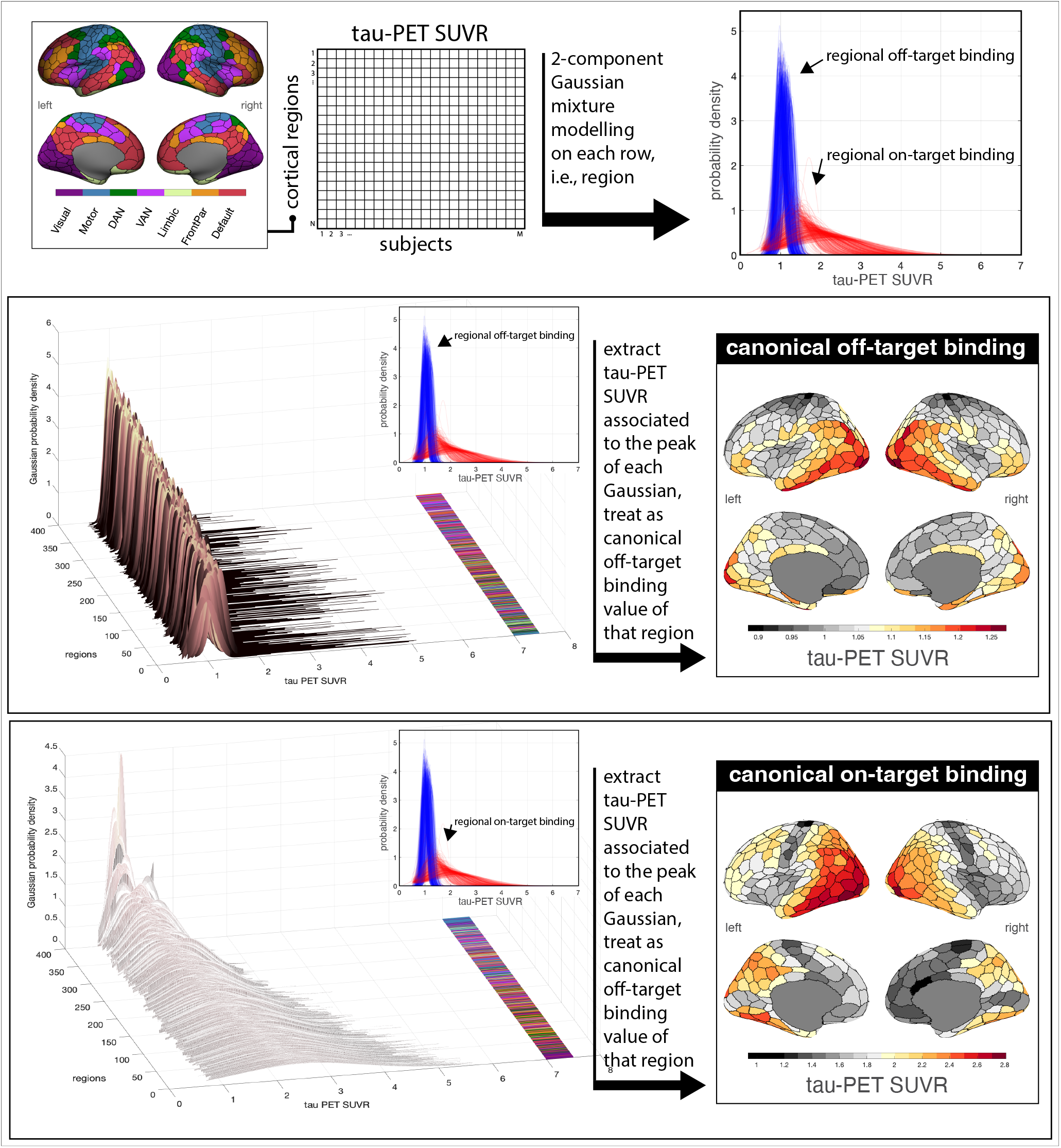
Derivation of canonical PET patterns. For each cortical region, a two-component Gaussian mixture model was fitted on tau-PET SUVR values of that region in the cohort (Vogel et al., 2020). The mean of left- and right-hand Gaussian are treated as the canonical off- and on-target tau-PET value of that region, respectively; the standard deviation of each Gaussian can also be used to reflect the uncertainty of each canonical estimate. This analysis results in an off- and on-target binding tau-PET spatial map defined at the resolution of the atlas used, with the resulting spatial map for each atlas being shown in Fig. S9 and Fig. S10, respectively.

**Fig. S9.**
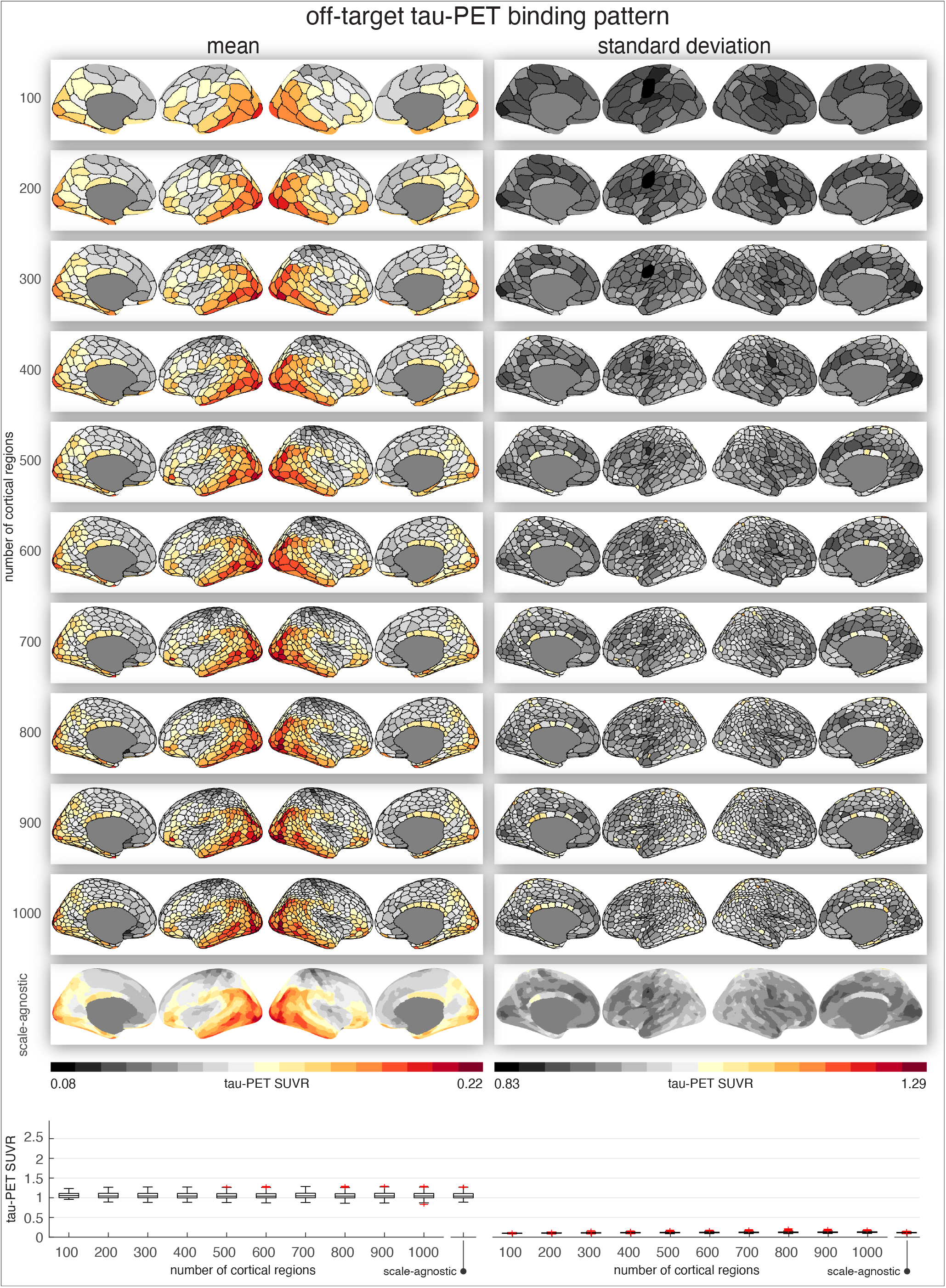
Canonical off-target binding tau-PET patterns across spatial scales. A unit set of off-target tau-PET binding value is obtained for each region defined by an atlas at a given resolution, resulting in a spatial pattern at the resolution of the atlas (see the first 10 rows of the first column); the second column shows the standard deviation of the estimated off-target binding values for each region. The last row in the surface plots show the scale-average off-target binding tau-PET maps and its standard deviation, obtained by averaging the 10 atlas-resolution maps. The bottom panel compares boxplots of the distribution of values across all the displayed surface maps using a fixed y-axis range.

**Fig. S10.**
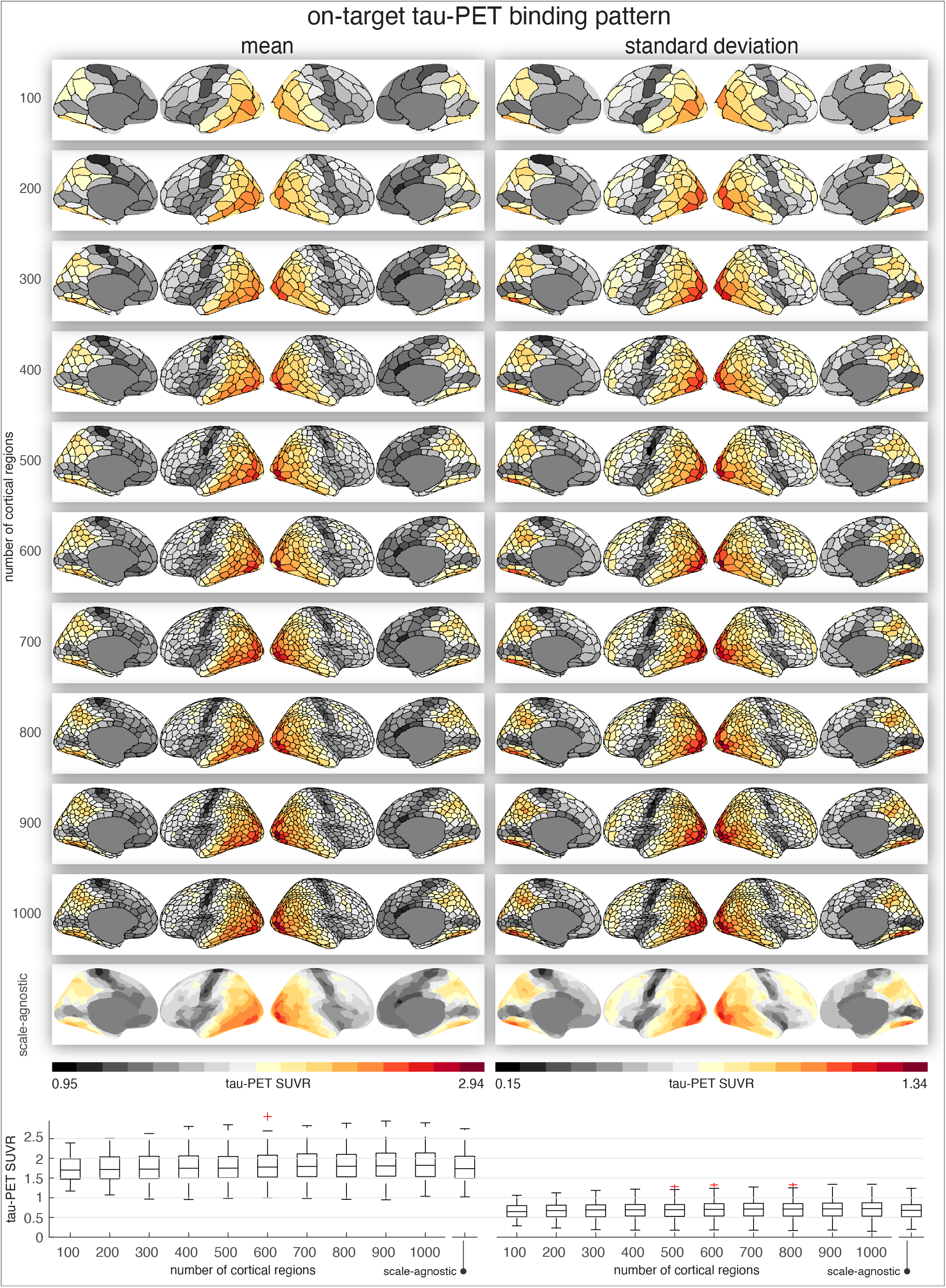
Canonical on-target binding tau-PET patterns across spatial scales. A unit set of on-target tau-PET binding value is obtained for each region defined by an atlas at a given resolution, resulting in a spatial pattern at the resolution of the atlas (see the first 10 rows of the first column); the second column shows the standard deviation of the estimated on-target binding values for each region. The last row in the surface plots show the scale-average on-target binding tau-PET maps and its standard deviation, obtained by averaging the 10 atlas-resolution maps. The bottom panel compares boxplots of the distribution of values across all the displayed surface maps using a fixed y-axis range.

**Fig. S11.**
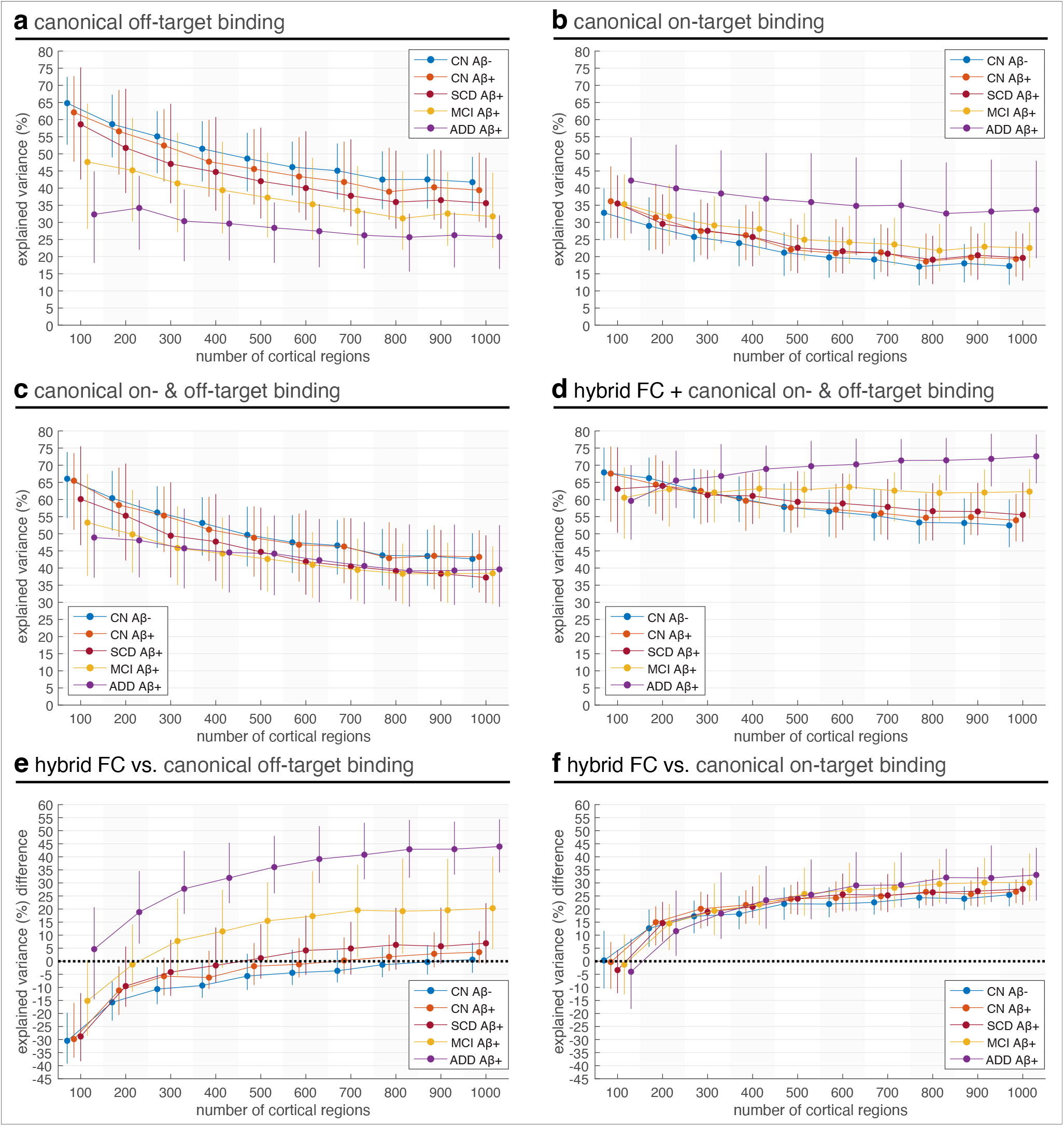
Explained variance of individual tau-PET using canonical tau-PET regressors: (a) canonical off-target binding tau-PET pattern, (b) canonical on-target binding tau-PET pattern, (c) canonical off- and on-target tau-PET binding patterns, and (d) hybrid FC and canonical off- and on-target binding patterns, across scales and across groups. Difference in explained variance of individual tau-PET patterns using hybrid FC compared to using (e) canonical off-target tau-PET binding pattern and (f) canonical on-target tau-PET binding pattern. In (a)–(f), explained variances represent corrected *R*^2^ to facilitate comparison of explained variance using models with different numbers of regressors. In all panels, whiskers represent the 25th to 75th percentiles across subjects; markers indicate the median. Template FC refers to the group-derived functional connectivity matrix estimated from amyloid-negative cognitively healthy controls. Hybrid FC combines this template FC representation with each participant’s subject-specific FC using region- and subject-specific weights.

**Fig. S12.**
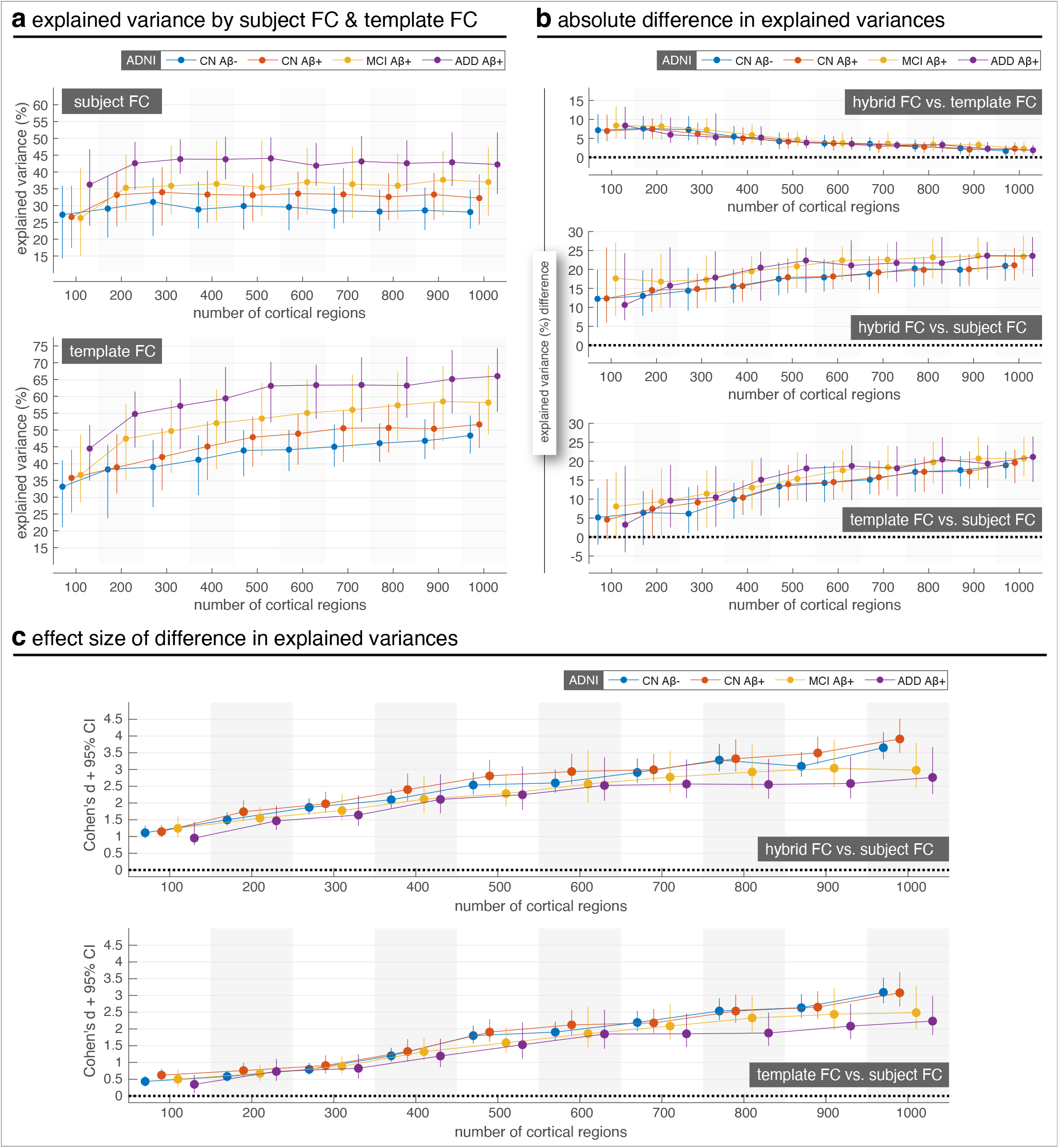
External replication of hybrid FC performance relative to subject FC and template FC in ADNI. (a) Explained variance of individual tau-PET patterns using subject FC and template FC across atlas resolutions. (b) Absolute differences in explained variance between hybrid FC and template FC, hybrid FC and subject FC, and template FC and subject FC. Positive values indicate greater explained variance for the first FC representation named in each comparison. (c) Standardized differences in explained variance for hybrid FC versus subject FC and template FC versus subject FC, quantified using Cohen’s *d*. Positive Cohen’s *d* values indicate greater explained variance for the first FC representation named in each comparison. Points and error bars show group-level mean *±* 95% confidence interval across subjects for each clinical group and atlas resolution. Template FC refers to the group-derived functional connectivity matrix estimated from amyloid-negative cognitively healthy controls; subject FC was estimated separately from each participant’s rs-fMRI data. CN: cognitively normal; MCI: mild cognitive impairment; ADD: Alzheimer’s disease dementia.

**Fig. S13.**
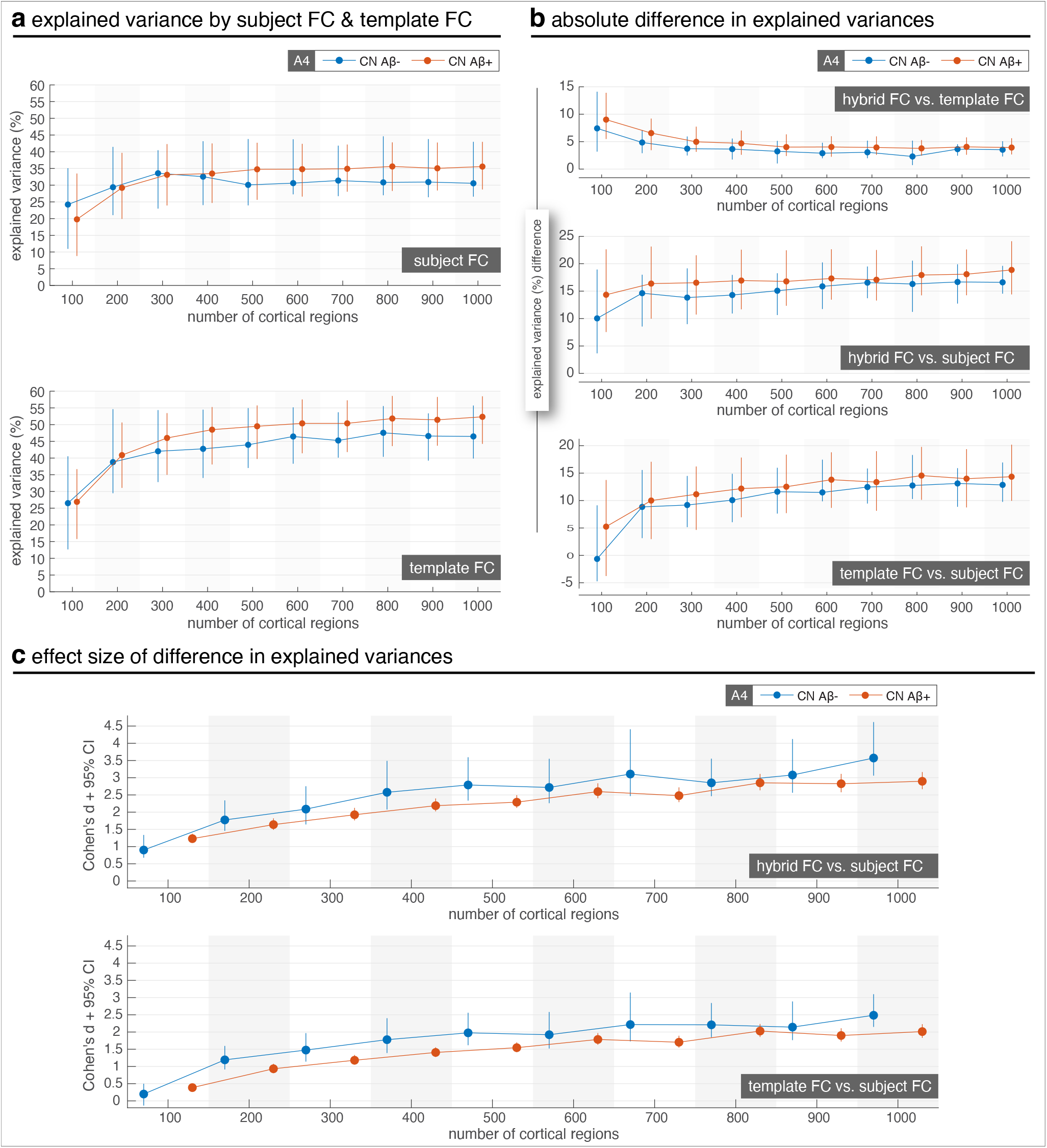
External replication of hybrid FC performance relative to subject FC and template FC in A4. (a) Explained variance of individual tau-PET patterns using subject FC and template FC across atlas resolutions. (b) Absolute differences in explained variance between hybrid FC and template FC, hybrid FC and subject FC, and template FC and subject FC. Positive values indicate greater explained variance for the first FC representation named in each comparison. (c) Standardized differences in explained variance for hybrid FC versus subject FC and template FC versus subject FC, quantified using Cohen’s *d*. Positive Cohen’s *d* values indicate greater explained variance for the first FC representation named in each comparison. Points and error bars show group-level mean *±* 95% confidence interval across subjects for each clinical group and atlas resolution. Template FC refers to the group-derived functional connectivity matrix estimated from amyloid-negative cognitively healthy controls; subject FC was estimated separately from each participant’s rs-fMRI data. CN: cognitively normal.

**Fig. S14.**
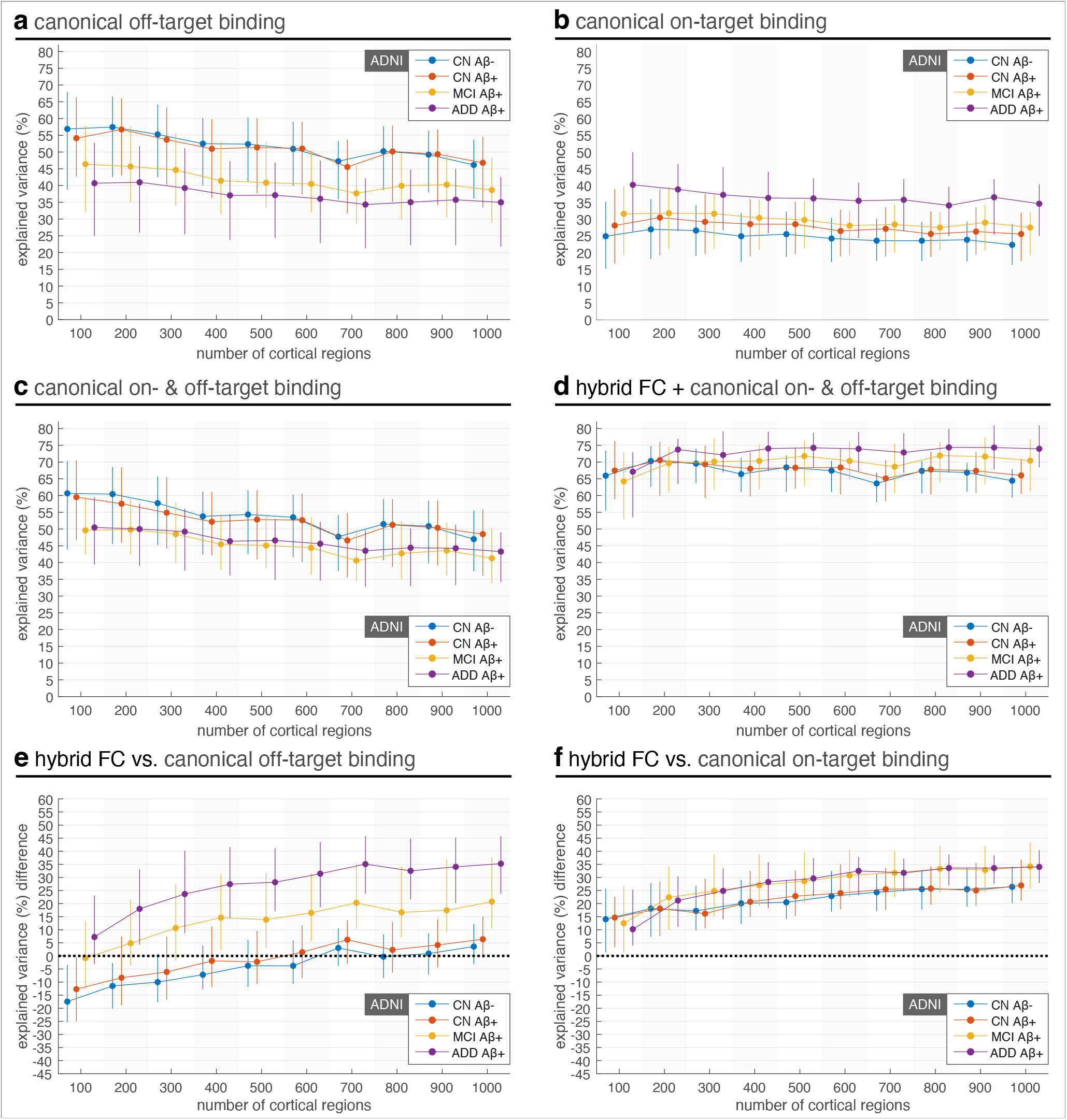
External replication of hybrid FC performance relative to canonical tau-PET patterns in ADNI. (a–c) Explained variance of individual tau-PET patterns by canonical off-target binding tau-PET, canonical on-target binding tau-PET, and canonical on- and off-target binding tau-PET across atlas resolutions. (d) Explained variance of individual tau-PET patterns by a combined model including hybrid FC together with canonical on- and off-target binding tau-PET. (e–f) Absolute differences in explained variance between hybrid FC and canonical off-target binding tau-PET, and between hybrid FC and canonical on-target binding tau-PET. Positive values indicate greater explained variance by hybrid FC. Points and error bars show group-level mean *±* 95% confidence interval across subjects for each clinical group and atlas resolution. CN: cognitively normal; MCI: mild cognitive impairment; ADD: Alzheimer’s disease dementia.

**Fig. S15.**
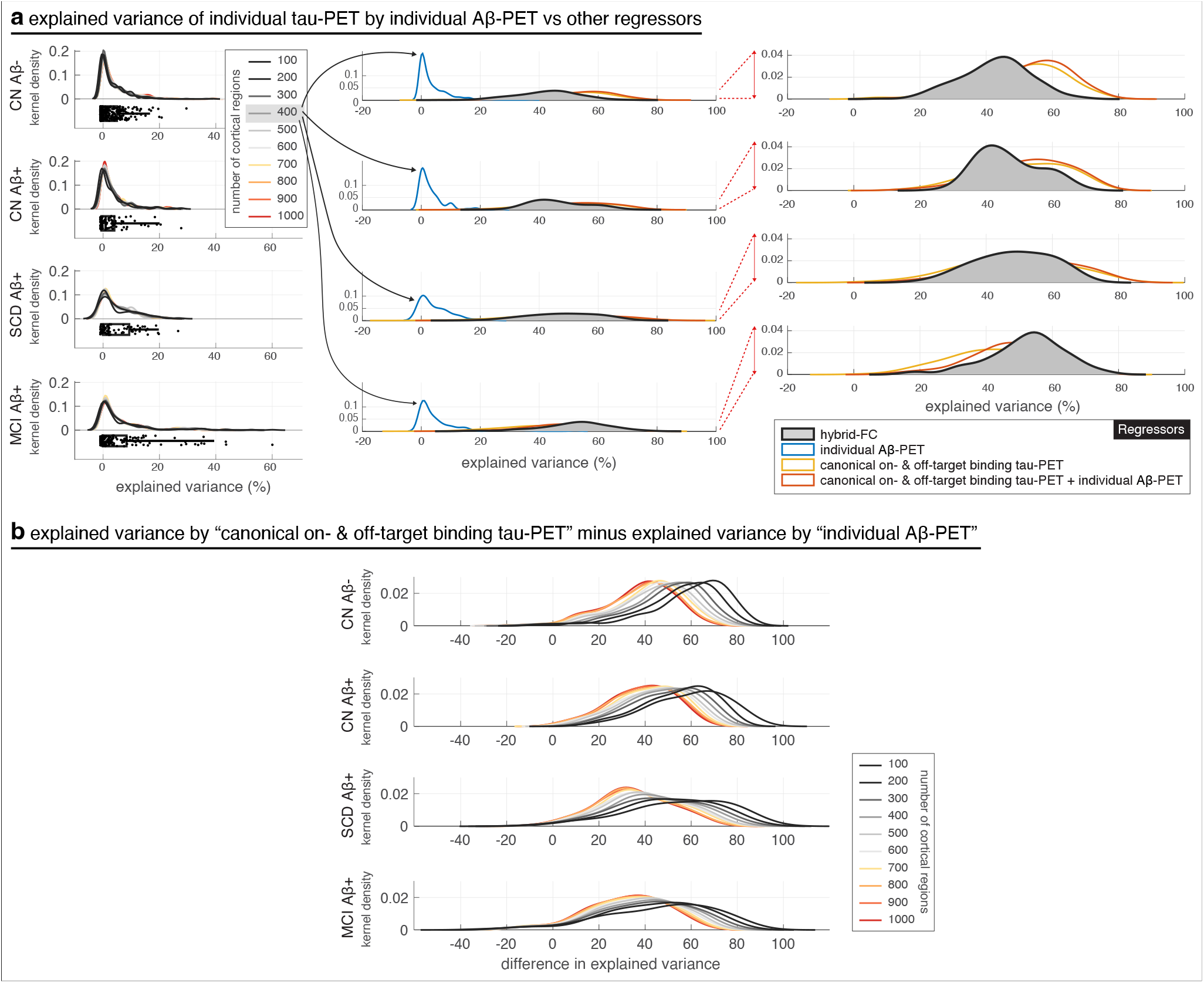
Comparison of performance of individual Aβ-PET patterns in explaining individual tau-PET patterns relative to the performance of canonical tau-PET patterns and individual hybrid FC. Patients with ADD are not included in results in this figure since they did not have an Aβ-PET scan. (a) Explained variances of individual tau-PET patterns using individual Aβ-PET are compared to other regressors; performance of individual Aβ-PET across spatial scales is shown on the left and on the right its performance is compared against other regressors for cortical parcellation with 400 regions. Individual Aβ-PET on its own minimally explains the spatial profile of individual tau-PET patterns, across spatial scales. Furthermore, when combined with canonical tau-PET patterns, the explained variance only minimally increases in CN individuals. In patients with MCI, the increase is slightly larger, yet, the performance remains below the performance of hybrid FC in explaining individual tau-PET. (b) The difference in explained variance of individual tau-PET by canonical-tau-PET patterns relative to individual Aβ-PET patterns is shown across spatial scales. The difference in performance decreases as the spatial resolution increases across the three groups. The differences in explained variances are quite in par across the three groups at high spatial resolutions whereas it is slightly lower at low spatial resolution in patients with MCI. All in-sample explained-variance values are reported as corrected *R*^2^.

**Fig. S16.**
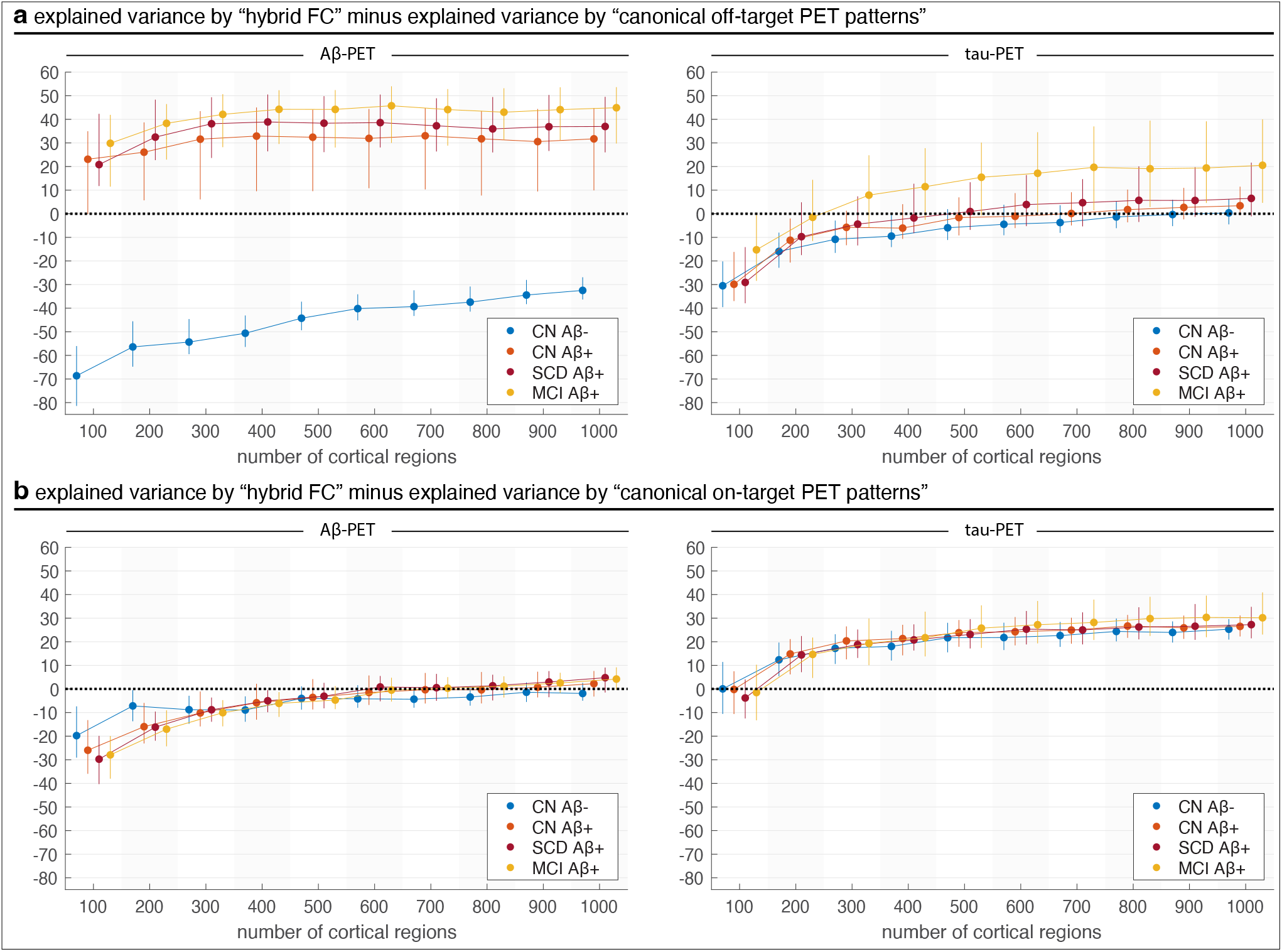
Comparison of differences in explained variance between hybrid FC and canonical PET pattern models for individual tau-PET and Aβ-PET, across spatial scales. (a) Hybrid FC vs. canonical off-target patterns: Aβ-PET is better explained by canonical patterns in CN Aβ+ and MCI, while hybrid FC outperforms for tau-PET across all groups. (b) Hybrid FC vs. canonical on-target patterns: Similar Aβ-PET trend as in (a), with narrowing differences at finer scales; tau-PET remains better captured by hybrid FC. These results, together with those shown in Fig. 5b, underscore a modality-specific dissociation: Aβ-PET is better explained by canonical spatial patterns, whereas tau-PET is more accurately explained by functional connectivity-based models, particularly at finer spatial resolutions. In all panels, whiskers represent the 25th to 75th percentiles; markers indicate the median, across subjects in each group and at each scale.

**Fig. S17.**
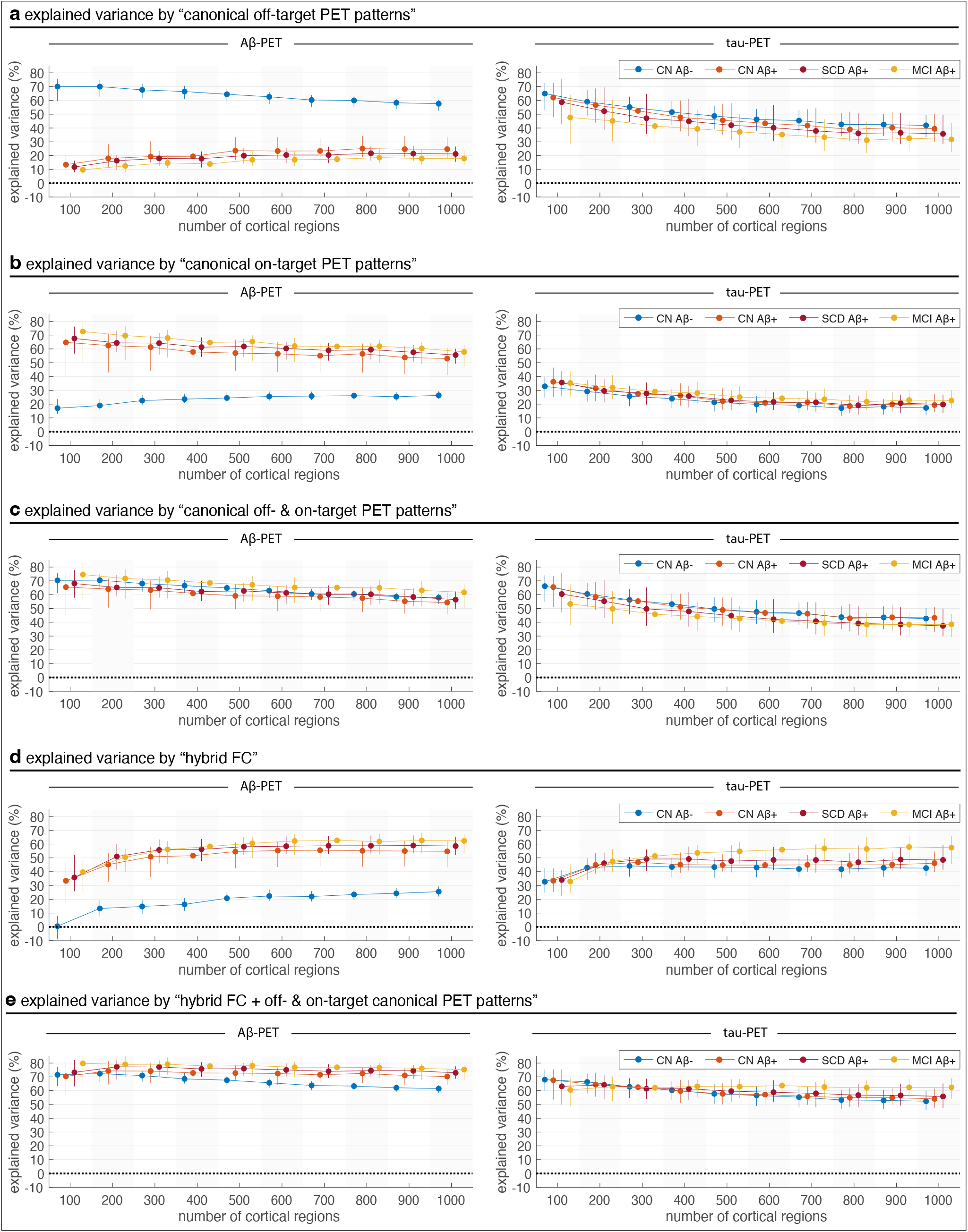
Explained variance of Aβ-PET and tau-PET patterns by different regressors across spatial scales. (a) Canonical off-target PET patterns, (b) canonical on-target PET patterns, (c) combined canonical off- & on-target PET patterns, (d) hybrid functional connectivity (FC), and (e) hybrid FC + canonical PET patterns. Canonical Aβ-PET patterns explain high Aβ-PET variance in Aβ+ groups with stable performance across scales. Tau-PET variance explained by canonical models declines with resolution and remains lower overall. Hybrid FC better explains tau-PET (especially in Aβ+ groups) than Aβ-PET, with improvements at finer scales. For Aβ-PET, hybrid FC adds little beyond canonical models. For tau-PET, combining hybrid FC with canonical PET patterns improves performance at higher resolutions. Results highlight modality-specific patterns: Aβ-PET aligns with predefined spatial maps; tau-PET is better captured by FC-based models. Whiskers: 25th–75th percentiles; markers: median across subjects. All in-sample explained-variance values are reported as corrected *R*^2^.

**Fig. S18.**
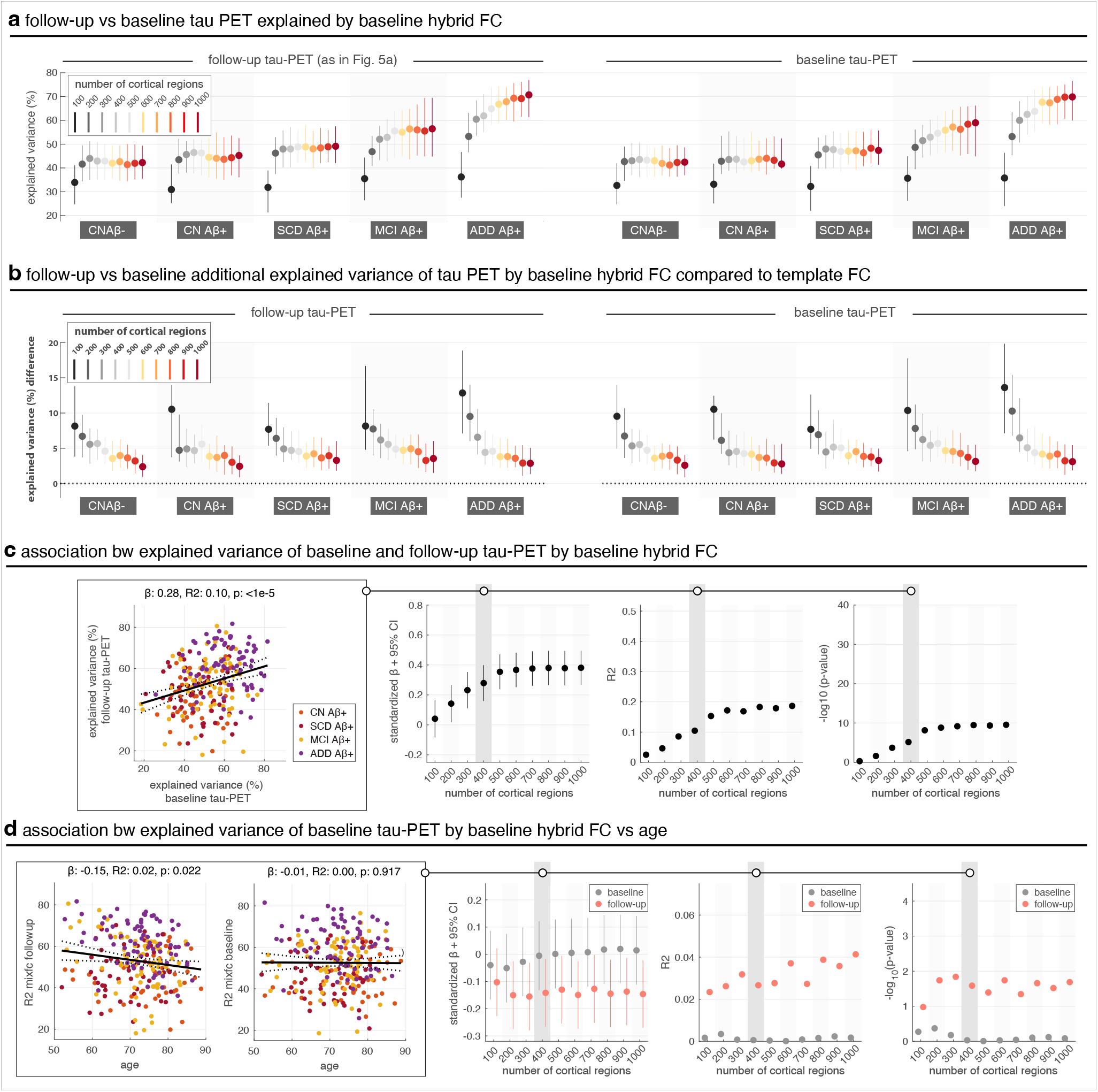
Comparison of explained variance in baseline vs. follow-up tau-PET by hybrid FC. (a) Baseline tau-PET patterns were explained by hybrid FC with a similar degree of accuracy as follow-up tau-PET, showing consistent increases with finer parcellations across all clinical groups. (b) Hybrid FC provided added explanatory value beyond template FC, especially at finer spatial scales and in MCI Aβ+ and ADD groups. (c) Although group-level variance explained was comparable between baseline and follow-up, individual-level associations between explained variance at the two timepoints were modest, with standardized effect sizes peaking around 400– 600 cortical regions but with little of the overall variance explained. (d) Association between age and hybrid FC explained variance at baseline and follow-up. Age was not associated with baseline explained variance, whereas a small negative association was observed for follow-up explained variance. Together, these results suggest group-level stability in FC–tau alignment, but also underscore subject-specific variability in how tau pathology evolves over time relative to baseline FC architecture.

**Fig. S19.**
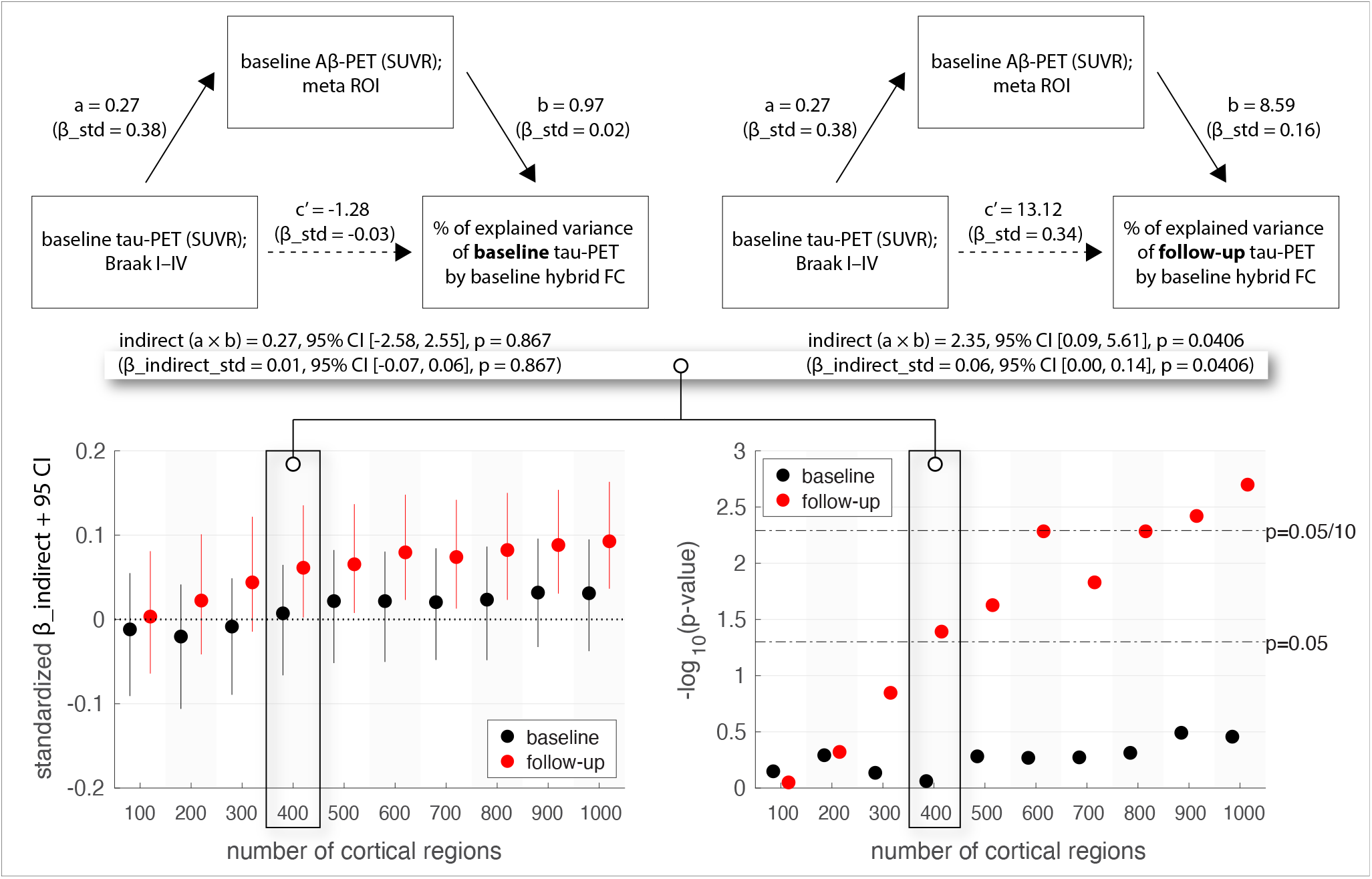
Mediation of baseline Aβ-PET between early aggregate tau burden and explained variance (EV) of tau-PET patterns by FC (models adjust for age and sex). Top: Mediation at a 400-region parcellation (unstandardized coefficients; standardized *β* in parentheses). Aβ significantly mediates the association between baseline Braak I–IV tau and EV follow-up tau-PET patterns explained by baseline hybrid FC, but not EV of baseline tau-PET patterns. Bottom: Across spatial scales, the mediated effect of Aβ for EV of follow-up tau-PET patterns strengthens with granularity and becomes significant from ~400 regions onward, whereas EV of baseline tau-PET patterns shows no mediation at any scale. Same longitudinal sample as used in Fig. 6, where patients with ADD are excluded as they lacked Aβ-PET scans. For Aβ-PET, the meta ROI SUVR is the average SUVR calculated from a global neocortical ROI, including prefrontal, lateral temporal, parietal, anterior cingulate and posterior cingulate/precuneus.

